# Optimizing Biophysical Large-Scale Brain Circuit Models With Deep Neural Networks

**DOI:** 10.1101/2025.04.07.647497

**Authors:** Tianchu Zeng, Fang Tian, Shaoshi Zhang, Xin Li, Ai Peng Tan, Bart Larsen, Mansour L. Sina, Fang Ji, Joanna Su Xian Chong, Kwong Hsia Yap, Christopher Chen, Nicolai Franzmeier, Sebastian N. Roemer-Cassiano, Sidhant Chopra, Carrisa V. Cocuzza, Justin T. Baker, Juan Helen Zhou, Marielle V. Fortier, Yap Seng Chong, Michael J. Meaney, Xi-Nian Zuo, Nagaendran Kandiah, Woon-Puay Koh, Eric Kwun Kai Ng, Voon Hao Lew, Fiona Jia Wen Goh, Alzheimer’s Disease Neuroimaging Initiative, Ruben C. Gur, Raquel E. Gur, Tyler M. Moore, Theodore D. Satterthwaite, Gustavo Deco, Avram J. Holmes, B.T. Thomas Yeo

## Abstract

Biophysical modeling provides mechanistic insights into brain function, spanning single-neuron dynamics to large-scale circuit models. These models are governed by biologically meaningful parameters, many of which can be experimentally measured. Some parameters are unknown, and optimizing them improves fit to experimental data, enhancing biological plausibility. However, existing methods require repeated, computationally expensive numerical integration of differential equations, limiting scalability to population-level datasets. Here, we introduce DELSSOME (DEep Learning for Surrogate Statistics Optimization in MEan field modeling), a framework that bypasses numerical integration by directly predicting whether parameter sets produce realistic brain dynamics. Across three large-scale circuit models, DELSSOME achieves a 1500-8000× speedup over numerical integration in predicting model realism. When embedded within an evolutionary optimization strategy, DELSSOME enables 50-100× faster parameter estimation without sacrificing agreement with numerical integration. Because of computational constraints, most studies simulate large-scale circuit models only at the group level. DELSSOME enables efficient individual-level optimization of the feedback inhibition control model. By collating 12,005 individuals across 14 datasets, we derive – for the first time – normative trajectories of cortical E/I ratio across the lifespan, revealing new insights into sex differences and network-specific patterns. This acceleration enables population-scale mechanistic modeling and unlocks new opportunities for understanding brain function.

## 1 Introduction

Biophysical modeling is a powerful approach for deriving mechanistic principles of brain function. The Nobel Prize-winning Hodgkin–Huxley model exemplifies this approach by mathematically explaining action potential generation in single neurons (Hodgkin & Huxley, 1952). Similarly, mean field models (MFMs) of local circuit function – such as feedback inhibition control (FIC) models – have yielded biophysically-plausible accounts of perceptual decision making (Brunel & Wang, 2001; Wong & Wang, 2006). Integrating MFMs into large-scale circuit models has enabled the bridging of neuroscientific research across multiple scales, from transcriptomics (Deco et al., 2021) and cellular density (Wang et al., 2019), through local circuits and whole-brain dynamics (Honey et al., 2009; Breakspear et al., 2010), to behavior (Froudist-Walsh et al., 2021).

The dynamics of biophysical brain models are governed by biologically meaningful parameters (e.g., membrane resting potentials), many of which can be measured from cellular neurophysiology or histology. However, some parameters are unknown and require tuning. The tuning process usually involves iteratively integrating biophysical differential equations and adjusting parameters to maximize similarity between simulated and empirical brain activity, such as resting-state functional MRI (rs-fMRI). Exhaustive search is the most common approach for optimizing model fit (Kringelbach et al., 2020; Müller et al., 2023; Pang et al., 2023), but is not scalable to multiple parameters. Gradient descent methods have been proposed (Deistler et al., 2025b), yet they require a differentiable cost function.

In contrast, evolutionary and Bayesian optimization iteratively sample candidate parameters from a proposal distribution, which is then updated based on their goodness of fit (Demirtaş et al., 2019; Kong et al., 2021). Applying these strategies to fit a large-scale FIC circuit model to rs-fMRI produces markedly more realistic brain dynamics (Demirtaş et al., 2019; Zhang et al., 2024). The FIC model explicitly captures the balance of excitation and inhibition in cortical regions (Deco et al., 2021) – a core principle of cortical function critical to development (Larsen et al., 2022), aging (Richardson et al., 2013) and neuropsychiatric disorders (Anticevic & Lisman, 2017; Lauterborn et al., 2021). The fitted FIC models can thus generate a marker of whole-cortex excitation/inhibition (E/I) ratio, which has been validated with pharmacological fMRI and shows strong spatial convergence with positron emission tomography (Zhang et al., 2024).

A key limitation of previous approaches is the computational cost of numerically integrating differential equations. Emerging deep neural networks (DNNs) learn the mapping from equation parameters to simulated time courses (Raissi et al., 2019; Li et al., 2021; Kovachki et al., 2023). A feedforward pass through a trained DNN is extremely fast, bypassing costly numerical integration. However, DNNs predicting full time courses are typically designed for deterministic systems and replace only hundreds of numerical integration steps (Li et al., 2021; Kovachki et al., 2023; Ghafourpour et al., 2025). In contrast, large-scale brain models rely on stochastic differential equations requiring over 100,000 steps (Lam et al., 2022; Pang et al., 2023; Zhang et al., 2024). Another DNN-based framework, simulation-based inference (SBI; Cranmer et al., 2020; Gonçalves et al., 2020; Tolley et al., 2024) predicts model parameters directly from simulated data. SBI is effective when simulations closely match empirical data, but degrades substantially when mismatches occur (Boelts et al., 2025; Deistler et al., 2025a). For large-scale circuit models, close correspondence between simulated and empirical in-vivo brain imaging data is rarely achievable.

Here, we propose DELSSOME (DEep Learning for Surrogate Statistics Optimization in MEan field modeling), a framework for optimizing large-scale circuit models governed by stochastic differential equations. Unlike SBI (Gonçalves et al., 2020; Tolley et al., 2024), DELSSOME is trained on both simulated and empirical data, so it does not require the biophysical model to fully replicate the biological system. Rather than predicting entire time courses (Li et al., 2021; Ghafourpour et al., 2025), DELSSOME directly predicts whether a parameter set produces realistic brain dynamics based on a rich set of neuroscience-informed surrogate statistics, thus reducing the prediction targets from millions of variables to several key metrics.

When embedded within an evolutionary optimization strategy, DELSSOME achieves a 50-100× speedup over numerical integration for FIC (Deco et al., 2014), MFM (Deco et al., 2013) and Hopf (Ponce-Alvarez & Deco, 2024) models. DELSSOME’s parameter estimates closely match numerical integration, whereas SBI (Gonçalves et al., 2020; Tolley et al., 2024) produces substantially less accurate parameters. Due to computational constraints, the vast majority of studies usually simulate large-scale circuit models at the group level (Demirtaş et al., 2019; Müller et al., 2023; Pang et al., 2023). DELSSOME enables efficient fitting of the FIC model at the individual-level. Using data from 12,005 individuals across 14 datasets, we derive normative trajectories of cortical E/I ratio across the lifespan. This massive acceleration allows biophysical models to be deployed at population scale, opening the door to new mechanistic insights into brain function.

## 2 Results

### 2.1 Optimizing the feedback inhibition control (FIC) model with numerical integration

The FIC model (Deco et al., 2014) is a neural mass model derived through a mean-field reduction of a spiking neuronal network model (Brunel & Wang, 2001; Wong & Wang, 2006). Mean field models approximate the collective activity of many neurons within a region, rather than simulating each neuron individually. The FIC model comprises ordinary differential equations (ODEs) that capture the dynamics of excitatory and inhibitory neuronal populations within each cortical region (Fig. 1a; Supplementary Methods S1). Local dynamics are driven by recurrent interactions within the excitatory and inhibitory populations, as well as between excitatory and inhibitory populations. Higher excitatory-to-excitatory recurrent strength (w_EE_) and lower inhibitory-to-excitatory connection strength (w_IE_) increases activity of the excitatory population. Similarly, higher excitatory-to-inhibitory connection strength (w_EI_) and lower inhibitory-to-inhibitory recurrent strength (w_II_) enhance synaptic currents in the inhibitory population. The noise amplitude (σ) controls neuronal noise in each cortical region. Finally, the excitatory populations in different cortical regions are connected via a structural connectivity (SC) matrix, modulated by a global coupling constant G.

**Figure 1.**
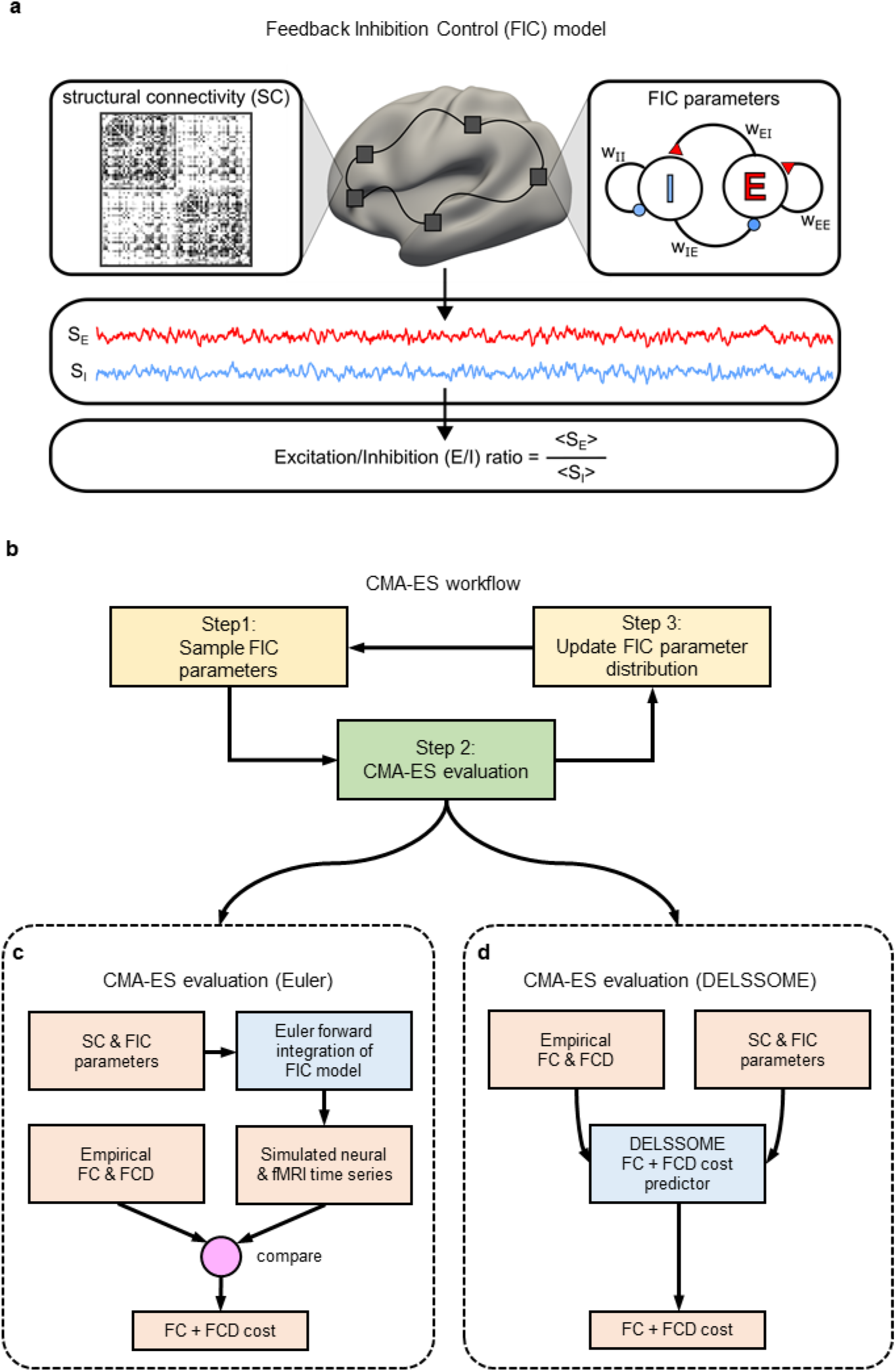
Overview of the Feedback Inhibition Control (FIC) model and its optimization with the covariance matrix adaptation evolution strategy (CMA-ES). **a.** The FIC model (Deco et al., 2014) comprises differential equations describing the neural dynamics of excitatory (“E”) and inhibitory (“I”) neuronal populations within each cortical region (right panel). Excitatory connections are represented by red triangles, and inhibitory connections are represented by blue circles. Connection strengths between neuronal populations are denoted as “w_XY_”, where x is the source population and y is the target population (e.g., w_EI_ represents the connection strength from the excitatory to the inhibitory population). Cortical regions are interconnected via excitatory connections parameterized by a structural connectivity (SC) matrix. For a given set of model parameters, time courses of excitatory (S_E_) and inhibitory (S_I_) synaptic gating variables, which represent the fraction of open channels, can be simulated. Specifically, S_E_ represents the fraction of open NMDA receptor channels on excitatory (pyramidal) neurons. The faster AMPA component is approximated at steady state owing to its rapid decay kinetics. S_I_ represents the fraction of open GABA receptor channels on inhibitory interneurons (Brunel & Wang, 2001; Deco et al., 2014). The excitation-inhibition ratio (E/I ratio) is defined as the ratio of the temporal average of S_E_ to S_I_ (Zhang et al., 2024). **b.** 100 sets of FIC parameters are sampled from an initial Gaussian distribution (Step 1). The 100 sets of FIC parameters are then evaluated (Step 2). The 10 sets of FIC parameters with the best evaluation metric are then used to update the Gaussian distribution (Step 3). **c.** To evaluate a set of FIC parameters, the FIC model is numerically integrated, resulting in simulated neural and fMRI time courses. Simulated fMRI time courses are then evaluated by computing a cost function that compares simulated and empirical functional connectivity (FC), as well as simulated and empirical functional connectivity dynamics (FCD), which we will refer to as FC+FCD cost. A lower FC+FCD cost indicates more realistic simulated fMRI time courses. **d.** The DELSSOME (Deep Learning for Surrogate Statistics Optimization in Mean Field Modeling) cost predictor will predict the FC+FCD cost without numerical integration. Throughout the current study, numerical integration was performed using the Euler method.

In our previous study (Zhang et al., 2024), a FIC model with spatially heterogeneous local circuit parameters was fitted to empirical fMRI data using the covariance matrix adaptation evolution strategy (CMA-ES; Hansen, 2006). CMA-ES is an optimization algorithm inspired by evolution, in which candidate parameter sets (“children”) are sampled, evaluated for fitness, and the best-performing sets guide the next generation. In the case of the FIC model, there were 10 unknown parameters (see Supplementary Methods S2 for details). The optimized FIC model was then used to generate excitatory and inhibitory synaptic gating variable time courses S_E_ and S_I_. The E/I ratio estimate was defined as the ratio of the temporal average of S_E_ and S_I_ (Fig. 1a).

To optimize the FIC model, CMA-ES samples 100 sets of candidate parameters from a randomly initialized 10-D Gaussian distribution (Step 1 in Fig. 1b). The 10-D Gaussian corresponds to the fact that we are trying to optimize 10 parameters. Each set of candidate parameters is then used to compute an evaluation metric that measures the realism of the resulting FIC model (Step 2 in Fig. 1b). The 10 sets of candidate parameters with the best evaluation metric are then used to update the sampling distribution for the next epoch (Step 3 in Fig. 1b). These three steps constitute one epoch of the CMA-ES algorithm.

Each CMA-ES epoch involves 100 children, so with 100 epochs, we need to evaluate 10,000 sets of FIC parameters (Step 2 in Fig. 1b), which is the most computationally expensive step of CMA-ES. More specifically, for a given set of FIC parameters, neural and fMRI timecourses are simulated via numerical integration of the FIC differential equations (Fig. 1c). Numerical integration is a method for accurately approximating the solution of differential equations over time when exact solutions are not available. Euler integration is the simplest such approach, which estimates dynamics by stepping forward in small increments. Throughout the current study, we always use the Euler method to perform numerical integration.

The simulated fMRI time courses were then evaluated by comparison with empirical (real) fMRI time courses using three evaluation metrics (Fig. 1c). First, the simulated and empirical time courses can be used to compute simulated and empirical functional connectivity (FC) matrices respectively. Each FC element reflects the synchronicity of the time courses of a given pair of brain regions. The similarity of the static and empirical FC was computed using the Pearson correlation (*r*). Because *r* does not account for differences in scale, we also computed the absolute difference (*d*) between the means of the empirical and simulated FC matrices (Demirtaş et al., 2019), where smaller *d* indicates greater similarity. The inclusion of *d* was necessary to prevent overly synchronized fMRI signals (Zhang et al., 2024).

Finally, the simulated and empirical fMRI time courses were also used to compute simulated and empirical functional connectivity dynamics (FCD) matrices respectively. While FC measures the overall synchronicity of a pair of brain regions over the entire fMRI session, FCD measures how the synchronicity of a pair of brain regions fluctuates throughout the fMRI session. Dissimilarity between the FCD matrices was computed using the Kolmogorov–Smirnov (KS) distance (Hansen et al., 2015; Kong et al., 2021; Zhang et al., 2024) of the FCD cumulative distribution functions (FCD-CDFs). The overall FC+FCD cost function was defined as *(1 – r) + d + KS*. A lower cost indicates greater realism in the simulated fMRI time courses. More details about CMA-ES are found in Supplementary Methods S2.

### 2.2 DELSSOME achieves 2000× speed-up for evaluating FIC model realism

To bypass the computationally intensive numerical integration in the CMA-ES evaluation (Fig. 1c), we propose the DELSSOME cost predictor to directly predict the FC+FCD cost (Fig. 1d). DELSSOME employs a transformer network (Devlin et al., 2019) to encode the dynamics of the FIC model. In parallel, multilayer perceptrons (MLPs) embed the FC and FCD. The resulting FIC model encoding and FC/FCD embeddings are then combined to predict the FC+FCD cost. More details are found in Sections 4.2 and 4.3.

To evaluate whether DELSSOME can effectively predict FIC model realism, we divided 1029 Human Connectome Project Young Adult (HCP-YA) participants into training (N = 680), validation (N = 180) and test (N = 169) sets. 50 participants were repeatedly sampled from the training set to generate 64 distinct group-level SC, FC and FCD-CDF samples. For each triplet of SC, FC and FCD-CDF, CMA-ES was then run to generate 10,000 sets of FIC model parameters and corresponding FC+FCD costs, resulting in 640,000 training samples for training DELSSOME. This procedure was repeated for the validation and test sets, yielding 140,000 and 130,000 samples respectively. The validation samples were used to tune the hyperparameters. The final trained DELSSOME cost predictor was then applied to the test samples.

DELSSOME predicted the static FC correlation cost (*1 - r*) with *r* = 0.97 (Fig. 2a), the static FC mean absolute difference cost *d* with *r* = 0.86 (Fig. 2b), and the FCD *KS* cost with *r* = 0.92 (Fig. 2c). Fig.2 was obtained based on the 68-region Desikan-Killiany cortical parcellation (Desikan et al., 2006). Similar performance (Fig. S1) was obtained using a 100-region cortical parcellation (Yan et al., 2023). Overall, our results suggest that DELSSOME can predict whether the FIC model will generate realistic neural and fMRI dynamics without Euler integration. Because a single feedforward pass through the DELSSOME deep neural network is extremely fast, DELSSOME offers a 2000× speed up over Euler integration for evaluating FIC model realism (Figs. 2d & S1).

**Figure 2.**
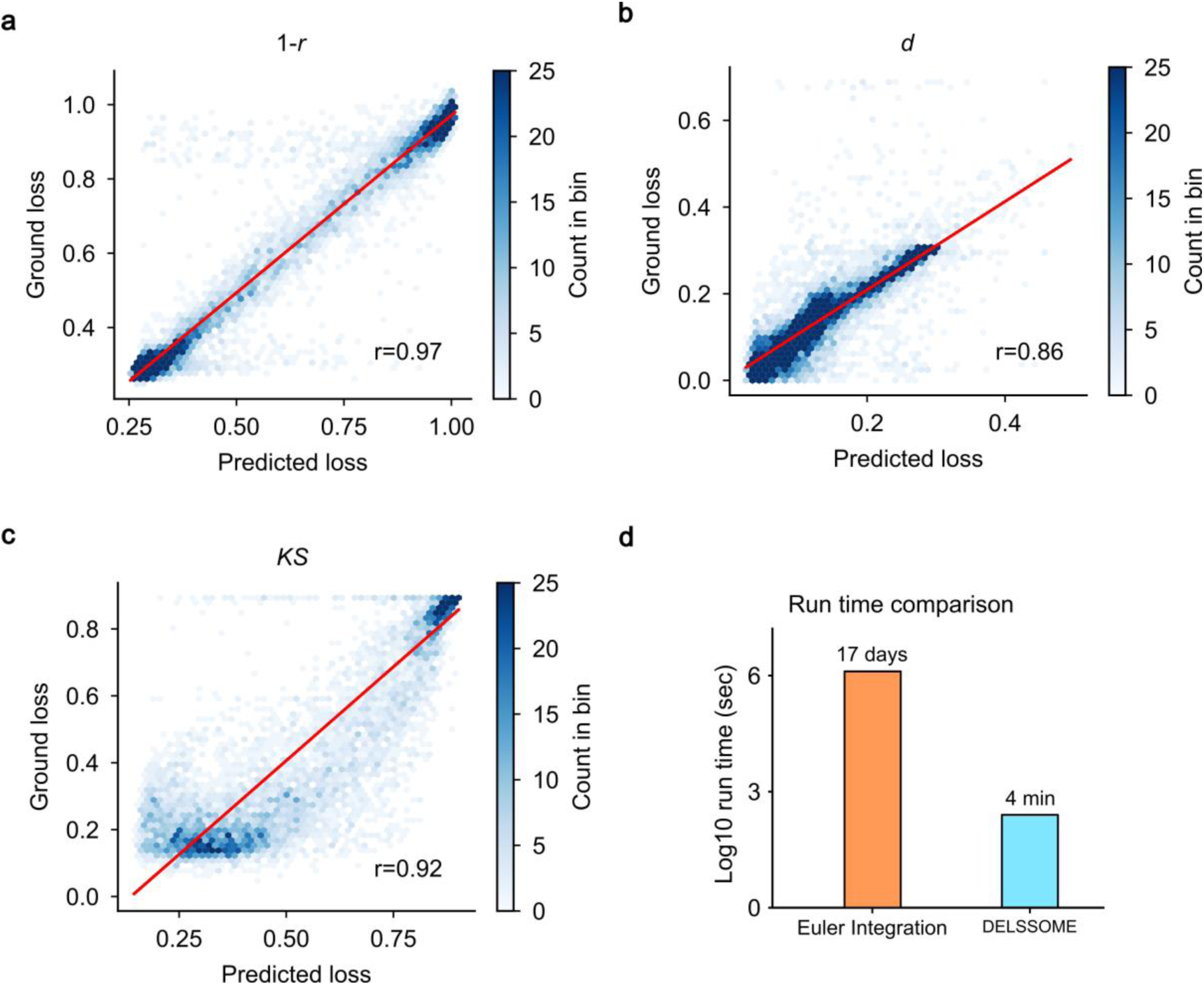
Test performance of DELSSOME FC+FCD cost predictor. **a.** Test performance of DELSSOME prediction of static FC cost (*1 - r*). **b.** Test performance of DELSSOME prediction of static FC cost (*d*). **c.** Test performance of DELSSOME prediction of FCD cost (*KS*). The Pearson’s correlation between the predicted and ground truth cost were at least 0.86. In all the analyses, ground truth was defined based on Euler integration, while the DELSSOME models avoided the Euler integration. **d.** Run time (log scale) of DELSSOME versus Euler integration in evaluating FIC model realism. DELSSOME offers a 2000× speed up over Euler integration.

### 2.3 DELSSOME achieves 50× speed-up for optimizing the FIC model

Although DELSSOME exhibited high accuracies (Fig. 2), small discrepancies between the DELSSOME predictions and ground truth (from Euler integration) might compound if DELSSOME was used to optimize the FIC model. Therefore, we tested whether DELSSOME could replace Euler integration in the CMA-ES algorithm (Fig. 1d). Here, we only considered the HCP-YA test participants since they were not used to train DELSSOME. The HCP-YA test participants were divided into the FIC model inversion training, validation and test sets (Fig. 3a).

**Figure 3.**
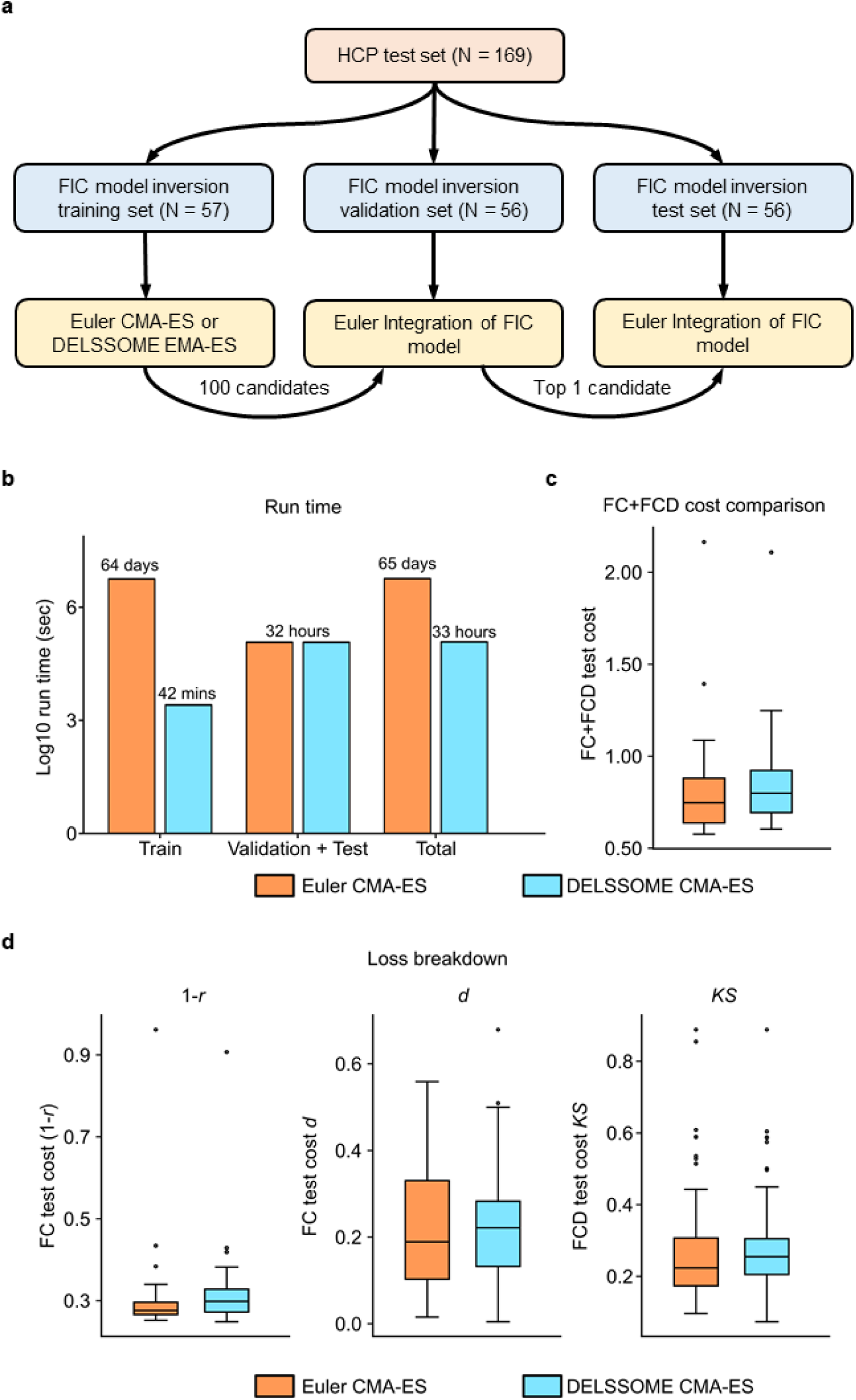
Comparison of Euler CMA-ES and DELSSOME CMA-ES in the HCP-YA test set. **a.** Split of the HCP-YA test set into FIC model inversion training, validation and test sets. Euler CMA-ES was run on the FIC model inversion training set and the 100 candidate parameter sets were evaluated in the FIC model inversion validation set. The best candidate parameter set was then evaluated in the test set. The procedure was repeated with DELSSOME CMA-ES replacing Euler CMA-ES in the training set. This comparison was repeated 50 times with different random initialization of the FIC model parameters each time. **b.** Run time (log scale) of DELSSOME CMA-ES versus Euler CMA-ES. DELSSOME CMA-ES offers a 2000× speed up over Euler CMA-ES in the training phase. If we also included validation and test phases in the run time, DELSSOME CMA-ES offers a 50× speed up over Euler CMA-ES. **c.** Total FC+FCD test cost comparison between DELSSOME CMA-ES and Euler CMA-ES (two-sample t-test, p = 0.38). Each boxplot contains 50 data points corresponding to the 50 repetitions of the procedure in panel (a). **d.** Breakdown of the FC+FCD test cost from panel **c** into the two FC costs (*1-r* and *d*) and one FCD cost (*KS*). DELSSOME significantly sped up the estimation of the FIC model parameters without any degradation in estimation quality.

Euler CMA-ES was run on the training set for 100 epochs to generate candidate parameter sets, which were then ranked on the validation set. The best parameter set was evaluated in the test set. The same procedure was repeated with DELSSOME replacing Euler integration in the FIC model inversion training set, although Euler integration was still used in the validation and test sets to preserve fidelity (Fig. 3a). The whole procedure was repeated 50 times with different random initializations.

During the training phase, DELSSOME CMA-ES was 2000 times faster than Euler CMA-ES (Fig. 3b). However, during the validation and test phases, the run time was the same since both approaches utilize Euler integration (Fig. 3b). Therefore, when accounting for all training, validation and test phases, DELSSOME CMA-ES was around 50× faster than Euler CMA-ES (Fig. 3b).

Importantly, FC+FCD costs between DELSSOME CMA-ES and Euler CMA-ES were similar (two-sample t-test p = 0.38; Fig. 3c), suggesting that the FIC parameters were equally good. Fig. 3 was obtained based on the 68-region Desikan-Killiany parcellation (Desikan et al., 2006). Similar performance (Fig. S2) was obtained using a 100-region cortical parcellation (Yan et al., 2023).

Replacing Euler integration with DELSSOME during both training and validation phases increased acceleration to ~100×, while maintaining comparable parameter estimates to Euler CMA-ES (two-sample t-test p = 0.24; Fig. S3). However, to be conservative, we recommend that Euler integration be retained in the validation phase.

Finally, while DELSSOME’s parameter estimates were comparable to full Euler integration, simulation-based inference (Gonçalves et al., 2020; Tolley et al., 2024) resulted in substantially worse parameters than Euler CMA-ES (Fig. S4a). The worse SBI performance arose because of mismatch between simulated and empirical data (Fig. S4b). Overall, the speed up from simulation-based inference came at the expense of model realism.

### 2.4 DELSSOME generalizes to a new dataset without further tuning

We note that DELSSOME not only encodes properties of the biophysical model, but also learns to predict the agreement between model simulations and empirical rs-fMRI data. Therefore, an important question is whether DELSSOME requires retraining for a new rs-fMRI dataset with different acquisition parameters and population characteristics. To evaluate whether DELSSOME CMA-ES generalizes to a new dataset, and also whether DELSSOME CMA-ES can extract similar neurobiological information as Euler CMA-ES, we replicated key findings of our previous study showing that E/I ratio decreases with age during neurodevelopment (Zhang et al., 2024). More specifically, the DELSSOME models trained from the HCP-YA dataset (previous section) were applied directly to the Philadelphia Neurodevelopment Cohort (PNC) dataset (Satterthwaite et al., 2014; Calkins et al., 2015) without any further tuning.

The PNC dataset comprised 885 participants aged 8 to 23 years old (Fig. 4a). Participants were sorted according to age (in months) and divided into 29 age groups (with 30 or 31 participants in each group). Within each age group, 15 participants were randomly selected as the validation set, while the remaining participants were assigned to the training set. Group-level FC and FCD-CDF were obtained by averaging across training and validation participants separately. For each age group, DELSSOME CMA-ES was applied to the training set for 50 epochs. The procedure was repeated five times with different random initializations, resulting in 250 candidate parameter sets. The 250 parameter sets were evaluated in the validation set with Euler integration. The best candidate parameter set was then used to generate an excitation/inhibition (E/I) ratio map using Euler integration. The same procedure was repeated with Euler CMA-ES. More details can be found in Supplementary Methods S4.

**Figure 4.**
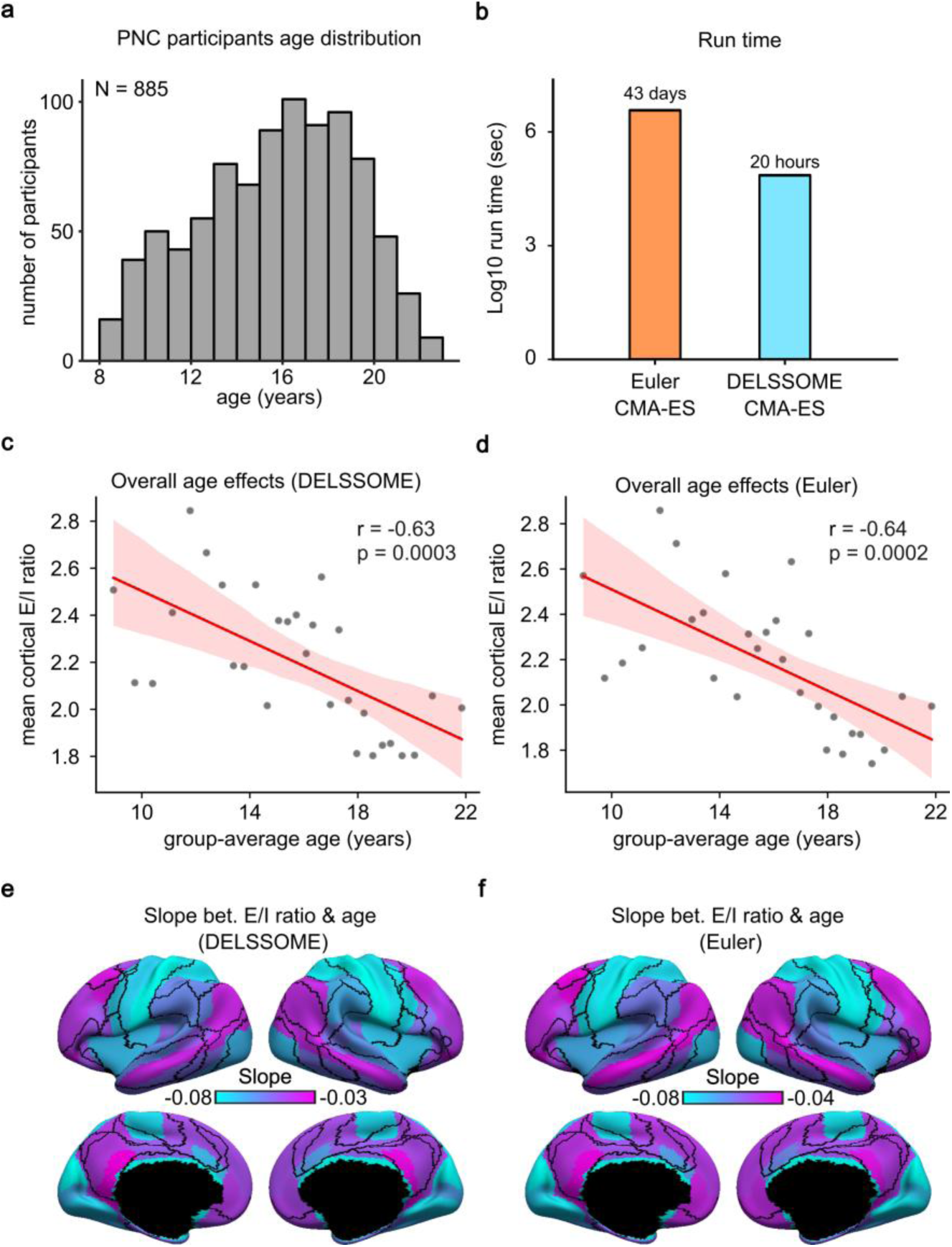
DELSSOME CMA-ES generalizes to the Philadelphia Neurodevelopmental Cohort (PNC) without further tuning. **a.** Age distribution of 885 PNC participants (mean = 15.66, std = 3.36, min = 8.17, max = 23). **b.** Run time comparison between DELSSOME CMA-ES and Euler CMA-ES. DELSSOME CMA-ES offers a 50× speed-up over Euler CMA-ES. **c.** Correlation between age and mean cortical E/I ratio estimated by DELSSOME CMA-ES. **d.** Correlation between age and mean cortical E/I ratio estimated by Euler CMA-ES. **e.** Regression slope between age and regional E/I ratio estimated by DELSSOME CMA-ES. **f.** Regression slope between age and regional E/I ratio estimated by Euler CMA-ES. All slopes in panels **e** and **f** are negative and significant after multiple comparisons correction with false discovery rate (FDR) q < 0.05.

DELSSOME CMA-ES was ~50 times faster than Euler CMA-ES (Fig. 4b). Consistent with the previous study (Zhang et al., 2024), both DELSSOME CMA-ES and Euler CMA-ES revealed a decrease in mean cortical E/I ratio with age (Figs. 4c,d). Pearson’s correlation between the 29 pairs of mean cortical E/I ratio (Figs. 4c,d) was 0.96. The decrease in E/I ratio was also more pronounced in sensory-motor regions than association cortex for both DELSSOME CMA-ES and Euler CMA-ES (Figs. 4e,f).

Since E/I ratio decreases with age, a lower E/I ratio might be an indicator of greater brain maturity. Therefore, another key result from our previous study was that among youth of the same age, lower E/I ratio was associated with better cognitive performance (Zhang et al., 2024). Here, we repeated the same analysis using the trained DELSSOME model from the HCP-YA dataset, yielding highly similar results to Euler CMA-ES (Fig. S5).

The previous analyses (Figs. 4 and S5) were performed using the 68-region Desikan-Killiany parcellation. Similar results were obtained with the 100-region Yan parcellation (Figs. S6 and S7). Overall, these results underscore the robustness of the DELSSOME CMA-ES approach in replicating previous findings relating E/I ratio and cognitive development, suggesting that DELSSOME CMA-ES can be used to replace Euler CMA-ES in future neuroscientific studies involving the FIC model.

### 2.5 DELSSOME generalizes to other large-scale circuit models

To evaluate the generalizability of the DELSSOME framework, we applied DELSSOME to two other large-scale circuit models with different dynamical regimes – the mean-field model (MFM; Deco et al., 2013; Kong et al., 2021) and the Hopf model (Ponce-Alvarez & Deco, 2024). The Hopf model has been used in neuroscience (Moon et al., 2015), and also other domains such as coupled lasers (Sciamanna et al., 2012). Details of MFM and Hopf can be found in Supplementary Methods S7 and S8 respectively.

For each large-scale circuit model, we trained the DELSSOME cost predictor using the same setup as for the FIC model (Sections 2.2 and 2.3). For MFM, DELSSOME CMA-ES achieved a 50× speedup over Euler CMA-ES (Fig. 5a), while producing parameter estimates of comparable quality (two-sample t-test p = 0.12; Fig. 5b). For the Hopf model, DELSSOME CMA-ES was 95× faster than Euler CMA-ES (Fig. 5d), again with no significant difference in estimation quality (two-sample t-test p = 0.28; Fig. 5e).

**Figure 5.**
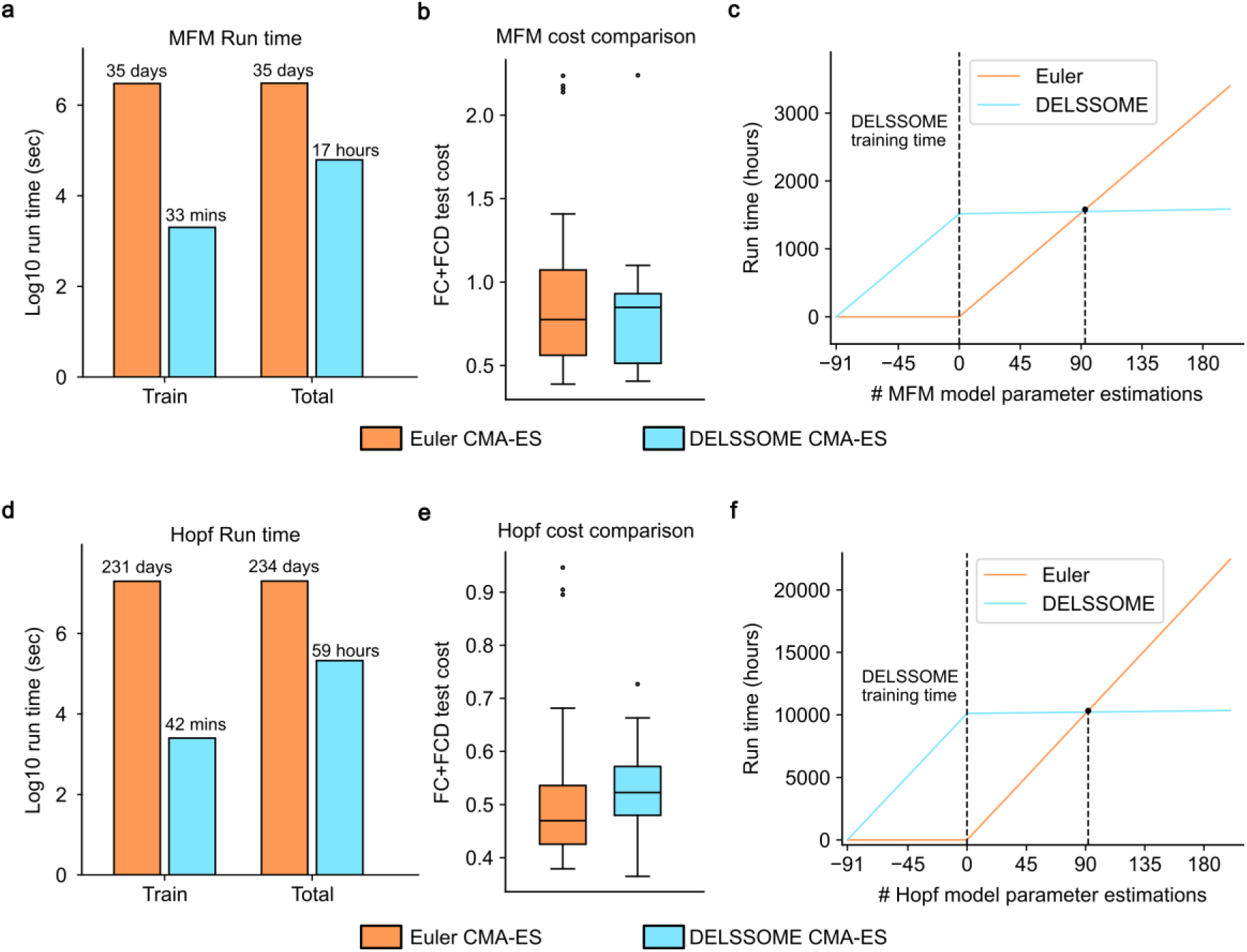
Comparison of Euler CMA-ES and DELSSOME CMA-ES applied to MFM and Hopf models in the HCP-YA test set. **a.** Run time (log scale) of DELSSOME CMA-ES versus Euler CMA-ES when applied to the mean field model (MFM). DELSSOME CMA-ES offers a 1500× speedup during training and 50× overall speedup versus Euler CMA-ES. **b.** Total FC+FCD test cost comparison between DELSSOME CMA-ES and Euler CMA-ES when applied to MFM (two-sample t-test p = 0.12). Each boxplot contains 50 data points corresponding to the 50 repetitions of the benchmarking procedure. **c.** Run time comparison taking into account the overhead of training DELSSOME for a new model, i.e., MFM. The breakeven point is 91 instances of MFM parameter estimation. **d.** Run time (log scale) of DELSSOME CMA-ES versus Euler CMA-ES when applied to the Hopf model. DELSSOME CMA-ES offers a 8000× speedup during training and 50× overall speedup versus Euler CMA-ES. **e.** Total FC+FCD test cost comparison between DELSSOME CMA-ES and Euler CMA-ES when applying to Hopf model (two-sample t-test p = 0.28). Each boxplot contains 50 data points corresponding to the 50 repetitions of the benchmarking procedure. **f.** Run time comparison taking into account the overhead of training DELSSOME for a new model, i.e., Hopf. The breakeven point is 91 instances of Hopf model parameter estimation.

Accounting for the one-time computational overhead required to train DELSSOME for a new large-scale circuit model (Figs. 5c,f), we estimate that studies involving more than 91 instances of model parameter estimation would realize net computational savings. For example, a study with 200 parameter estimation instances would achieve roughly 50% total runtime reduction. These results indicate that DELSSOME is likely to be advantageous even for moderate-sized studies.

Across all three models – FIC, MFM, and Hopf – we used the same transformer architecture, hyperparameters, and training procedure, without any model-specific architectural or training-level optimization. This suggests that DELSSOME may generalize to other large-scale brain circuit models without extensive model-specific modifications, with additional tuning potentially providing further gains.

### 2.6 DELSSOME enables efficient individual-level optimization of the FIC model

To move beyond group-average dynamics and assess whether DELSSOME can enable new biological insights, we applied DELSSOME to optimize FIC model parameters at the individual-level. To complement HCP-YA and PNC, we added 10 new datasets, yielding a total of 12 datasets. For each dataset, we randomly selected 40, 20 and 20 participants for the training, validation and test sets respectively. In the case of the HCP-YA dataset, care was taken to only select participants from the HCP-YA test set (Fig. 3a), which was not previously used to train the group-level DELSSOME model (Section 2.2).

The individual-level DELSSOME cost predictor utilized the same transformer architecture as group-level DELSSOME. The training set was used to train DELSSOME, while the validation set was used for hyperparameter tuning. Finally, the trained DELSSOME model was evaluated in the test set. Further details can be found in Supplementary Methods S6.

Across all 12 datasets, DELSSOME CMA-ES accelerated parameter optimization by approximately 50-fold while achieving comparable parameters to Euler CMA-ES (Figs. 6a and S8). We also applied the trained DELSSOME model, without further tuning, to two new datasets, where the quality of estimated parameters was again comparable with Euler CMA-ES (Figs. 6b,c), demonstrating strong out-of-distribution generalization of the trained DELSSOME cost predictor.

**Figure 6.**
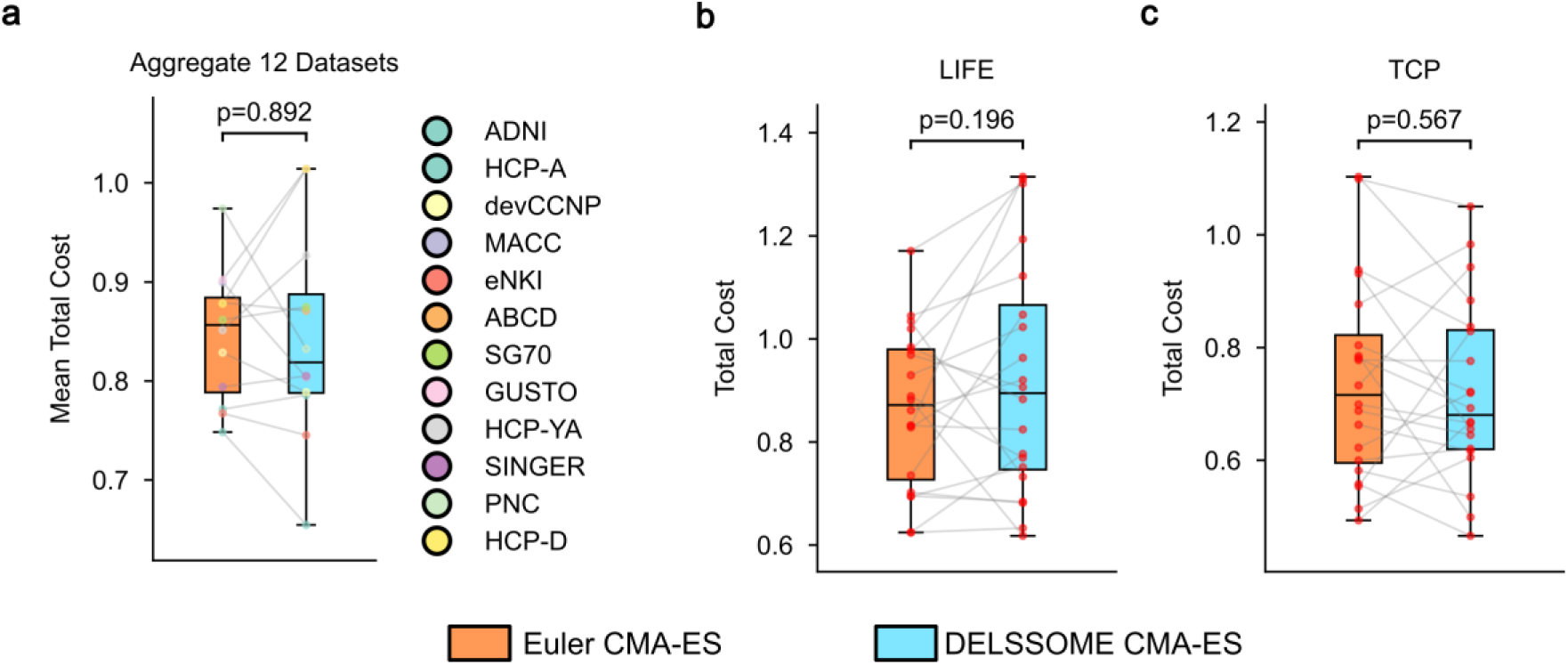
Comparison of Euler CMA-ES and DELSSOME CMA-ES for individual-level FIC parameter optimization. **a**. Across 12 datasets, the quality of estimated parameters was equivalent between DELSSOME CMA-ES and Euler CMA-ES (p = 0.892). There are 12 colored dots in each boxplot, corresponding to the 12 datasets. Each colored dot represents the averaged cost across 20 test participants. **b**. In the new unseen LIFE dataset, the quality of the estimated parameters was equivalent between DELSSOME CMA-ES and Euler CMA-ES (p = 0.196). c. In the new unseen TCP dataset, the quality of the estimated parameters was equivalent between DELSSOME CMA-ES and Euler CMA-ES (p = 0.567). P-values were computed based on two-sided paired-sample t-tests.

### 2.7 Normative cortical E/I ratio trajectory across lifespan

Previous studies of E/I balance have been limited to hundreds of individuals (Zhang et al., 2024; Saberi et al., 2025), precluding the characterization of population-level variability and sex differences across the lifespan. Leveraging the scalability afforded by DELSSOME, we collated 12,005 individuals from 14 datasets in North America and Asia, spanning 4.5 to 98.4 years old.

DELSSOME CMA-ES was used to estimate the cortical E/I ratio for each individual. Following previous studies (Bethlehem et al., 2022; Sun et al., 2025), we fitted the generalized additive model for location, scale, and shape (GAMLSS; Stasinopoulos & Rigby, 2008) to derive a normative trajectory of mean cortical E/I ratio (Fig. 7), and the E/I ratio of each cortical region (Fig. 8). Further details can be found in Supplementary Methods S6.

**Figure 7.**
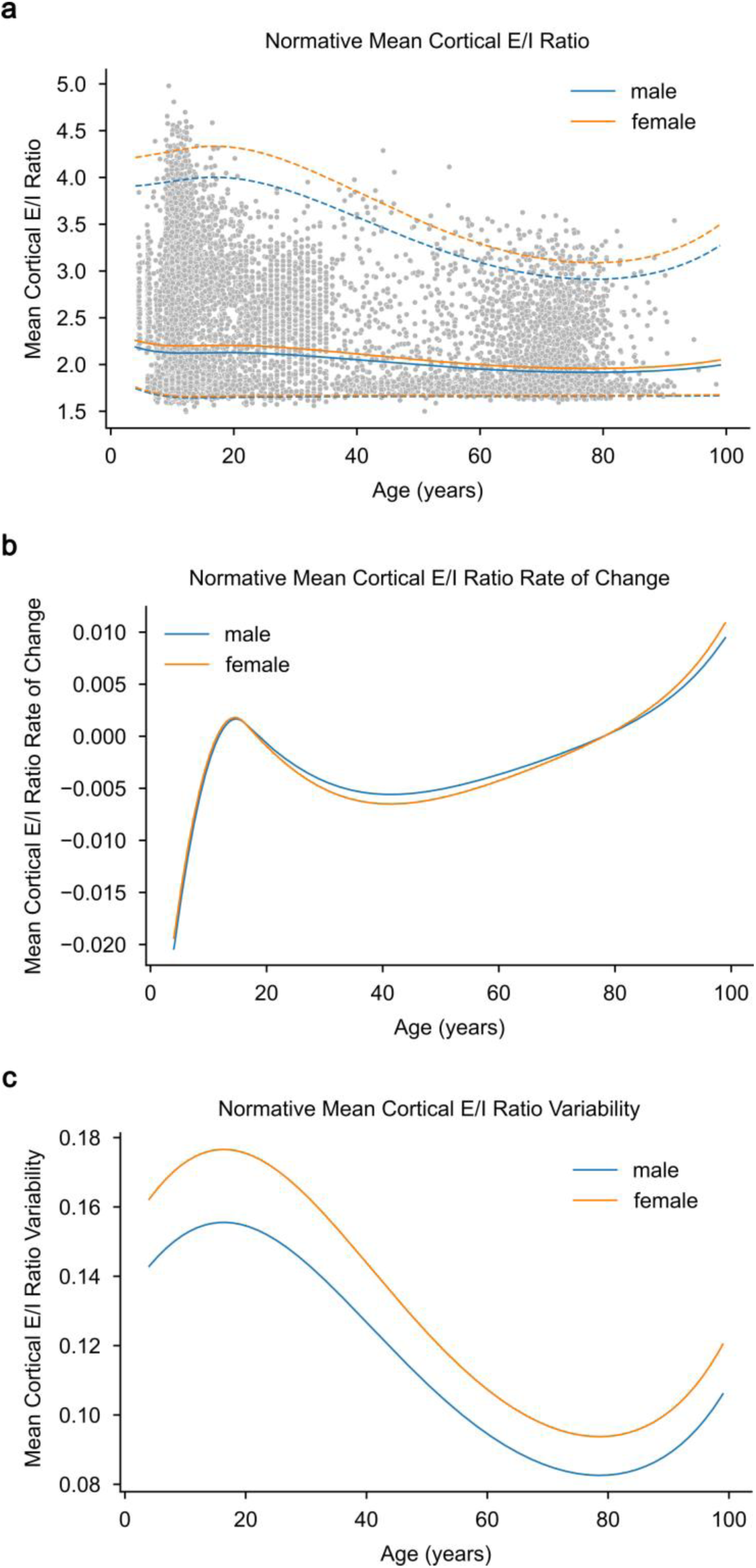
Normative trajectories of cortical excitation–inhibition (E/I) ratio across the human lifespan. **a.** Normative curves for E/I ratio estimated in 12,005 individuals using DELSSOME CMA-ES and modelled with the generalized additive model for location, scale, and shape (GAMLSS). These individuals spanned the age range from 4.5 to 98.4 years. Grey points denote individual estimates after site correction with GAMLSS. Solid lines show the median (50th centile) for males (blue) and females (orange). Dashed lines indicate the 2.5th and 97.5th centiles. **b.** Rate of change of the median E/I trajectory (first derivative of the median curve with respect to age), shown separately for males and females. Negative values indicate decreasing E/I ratio with age and positive values indicate increasing E/I ratio. **c.** Trajectories of between-individual variability in E/I ratio (scale parameter σ from the fitted GAMLSS) in males and females.

**Figure 8.**
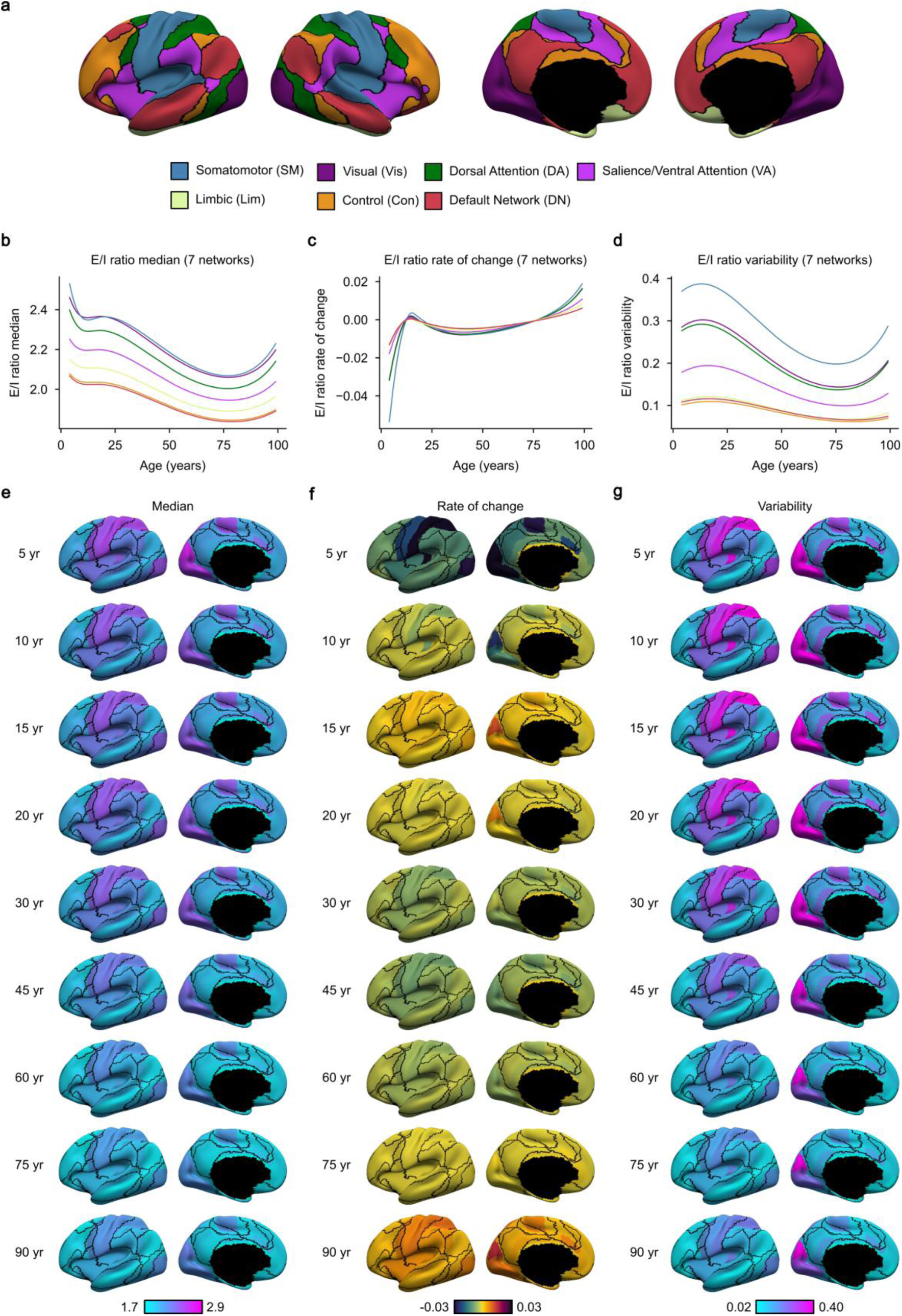
Normative trajectories of E/I ratio across cortical regions and networks. **a.** Seven large-scale resting-state networks (Yeo et al., 2011). **b.** Normative trajectories of E/I ratio (median) for each cortical network in 12,005 individuals using DELSSOME CMA-ES and the generalized additive model for location, scale, and shape (GAMLSS). **c.** Rate of change of the median E/I trajectories (first derivative of the median curve with respect to age) for each cortical network. **d.** Trajectories of between-individual variability in E/I ratio (scale parameter σ from the fitted GAMLSS) for each cortical network. **e.** Age-resolved cortical maps (shown for left hemisphere) summarizing the spatial distribution of median E/I ratio across the lifespan. **f.** Age-resolved cortical maps (shown for left hemisphere) summarizing the spatial distribution of rate of change of E/I ratio across the lifespan. **g.** Age-resolved cortical maps (shown for left hemisphere) summarizing the inter-individual variation in E/I ratio across the lifespan.

Mean cortical E/I ratio exhibited a nonlinear trajectory – declining during childhood, stabilizing around adolescence, continuing to decrease into early-to-mid adulthood, and increasing again in late life (Figs. 7a,b). This temporal motif was conserved across cortical networks (Fig. 8b). Across much of the lifespan, median E/I ratio was highest in early sensorimotor cortex—most prominently within the visual and somatomotor networks—whereas association cortex, including default and control networks, showed lower median E/I ratios (Fig. 8e).

Age-related modulation of E/I balance was likewise network dependent. The magnitude of change was greatest in sensorimotor systems and attenuated in association cortex (Figs. 8c,f), closely tracking the sensory–association neurodevelopmental axis (Sydnor et al., 2021; Larsen et al., 2023). Network differences in the rate of change were most pronounced during early development and late life (Figs. 8c,f).

Inter-individual variability also followed a nonlinear trajectory: dispersion was greatest in childhood, declined through adulthood, and increased again in late life (Fig. 7c). This pattern was shared across networks, but remained consistently higher in early sensorimotor networks than association networks (Figs. 8d,g), mirroring the previously observed sensory–association gradient in median E/I ratios and rate of E/I ratio change.

Males and females exhibited qualitatively similar lifespan trajectories and sensory-association gradient (Fig. S9). Nevertheless, clear sex differences were evident: females showed higher median E/I ratios (Fig. 7a) and greater inter-individual variability than males across the lifespan (Fig. 7c).

Together, these findings demonstrate that normative cortical E/I balance is structured along the sensory–association axis: early sensorimotor networks show higher E/I ratios, greater inter-individual variability, and stronger age-related modulation than transmodal association networks.

Importantly, these analyses were only computationally feasible using DELSSOME CMA-ES. Parameter optimization with Euler CMA-ES would have required 25 CPU hours per participant, translating to roughly 300,000 CPU hours – or 60 days using 200 CPUs – for all 12,005 participants. In contrast, generating the simulated data to train DELSSOME required 18,000 CPU hours, and training the cost predictor took 65 GPU hours on an RTX 3090. Once trained, DELSSOME CMA-ES was applied to all 12,005 participants in 6,000 CPU hours. Altogether, with 200 CPUs and a single GPU, the full DELSSOME pipeline completed in ~185 hours (or ~8 days), representing a massive reduction in computational cost relative to standard optimization.

## 3 Discussion

In this work, we introduced DELSSOME (Deep Learning for Surrogate Statistics Optimization in Mean-Field Modeling), a scalable framework for fitting large-scale brain circuit models. Rather than performing full numerical integration, DELSSOME predicts whether a candidate parameter set produces realistic brain dynamics using a compact set of surrogate statistics, achieving a 1,500–8,000× speedup across three large-scale circuit models. When paired with evolutionary optimization, it accelerates parameter estimation by 50–100× for the FIC (Deco et al., 2014), MFM (Deco et al., 2013), and Hopf (Ponce-Alvarez & Deco, 2024) models, while closely matching results of numerical integration. By dramatically reducing computational cost, DELSSOME enables practical individual-level model fitting. Leveraging data from 12,005 participants across 14 datasets, we characterized normative trajectories of cortical E/I balance across the lifespan.

### 3.1 Surrogate-statistics optimization

A growing literature uses deep learning to accelerate differential-equation–based modeling. Some methods learn solution operators or predict trajectories, e.g., physics-informed neural networks (Raissi et al., 2019) or neural operators (Li et al., 2021; Ghafourpour et al., 2025). Other methods learn inverse mapping from data to parameters, e.g., SBI (Gonçalves et al., 2020; Tolley et al., 2024). These approaches work well in deterministic systems or when the simulations closely match empirical data.

However, large-scale circuit models present two additional challenges: (1) they are stochastic and require long integrations to obtain stable statistics and (2) the optimization objective is rarely trajectory matching but instead focuses on summary statistics. DELSSOME addresses these challenges by aligning the surrogate’s prediction target with the optimization goal. Rather than predicting high-dimensional time courses, it evaluates whether parameters generate realistic brain dynamics. This enables accurate prediction of the FC+FCD cost and stable iterative search within CMA-ES, without accumulating surrogate errors that degrade performance.

DELSSOME also differs from SBI in its robustness to simulation-reality gaps. SBI can amortize inference, but its performance depends on the simulated data manifold covering empirical observations. Mismatches can substantially reduce performance (Boelts et al., 2025; Deistler et al., 2025a). By contrast, DELSSOME directly learns the agreement between candidate simulations and empirical brain data through the cost function itself. This makes it more resilient when simulations and reality diverge. Empirically, our benchmarking confirms this conceptual advantage: DELSSOME produces substantially more accurate parameter estimates than SBI (Fig. S4).

### 3.2 Normative E/I ratio trajectory across the lifespan

By leveraging DELSSOME’s computational efficiency, we provide the first estimates of normative E/I ratio trajectories across the lifespan (Figs. 7 and 8). Studies have reported decreases in E/I ratio during development (Larsen et al., 2022; Zhang et al., 2024; Saberi et al., 2025). With a substantially larger sample, we were able to fit more complex trajectories with GAMLSS, refining previous observations. We found that E/I ratio declines during childhood, stabilizes in adolescence, and continues to decrease into early adulthood. Extending prior work, we observed further decline in middle adulthood, followed by a late-life increase.

Ex-vivo rodent and in-vivo human studies have suggested age-related decline in GABA (Caspary et al., 2013; Gao et al., 2013; Porges et al., 2017; Simmonite et al., 2019; Chamberlain et al., 2021), but E/I ratio changes remained unclear since only GABA was measured. One in-vivo study indicated that GABA declined more than Glx (glutamine + glutamate), suggesting E/I ratio might rise in older adults (Petitet et al., 2021). However, these previous studies focused on single brain regions, leaving cortex-wide trajectories largely unknown. Our results fill this gap, revealing the full lifespan pattern of E/I ratio decline and late-life increase.

Interestingly, the spatial pattern of E/I ratio remains consistent across the lifespan. Sensorimotor networks consistently show higher E/I ratio than association networks, extending the sensory-association axis of neurodevelopment (Sydnor et al., 2021; Larsen et al., 2023) into adulthood and aging. This spatial motif was also evident in individual variability and rate of change. Finally, we observed that females have higher E/I ratio and greater inter-individual variability than males.

Beyond neuroscientific insights into E/I ratio evolution over the lifespan, these normative trajectories might also be useful for studying neuropsychiatric disorders. E/I imbalance has been implicated in conditions, ranging from schizophrenia (Anticevic & Lisman, 2017) to Alzheimer’s disease dementia (Lauterborn et al., 2021). Analogous to pediatric growth charts, the normative E/I trajectories we provide could help identify individuals whose cortical E/I balance deviates from typical patterns.

### 3.3 Use cases for DELSSOME

Pretrained DELSSOME models from this study are publicly available. Code for training DELSSOME and tuning hyperparameters is also provided for researchers interested in models beyond the pretrained versions. In our analyses, the same transformer architecture performed well for FIC, Hopf, and MFM models (Fig. 5), suggesting that researchers can adopt the same architecture. However, depending on the application, hyperparameter tuning may be necessary. For example, individual-level FC+FCD was noisier than group-level FC or FCD, so hyperparameter tuning was required for individual-level FIC parameter estimation.

Even accounting for the overhead of generating training data and training DELSSOME, studies with more than 91 parameter estimation instances would see net computational savings. For instance, a study with 200 instances could reduce total runtime by roughly 50%. DELSSOME is therefore advantageous even for moderate-sized studies, not just large datasets.

Furthermore, repeated parameter estimation is a routine feature of rigorous computational workflows, driven by statistical procedures (e.g., bootstrapping and permutation testing), sensitivity and control analyses, and the iterative refinement of modeling decisions and robustness checks. Consequently, computational efficiency is valuable not only for “big data”, but also for small datasets.

For example, in a previous study (Zhang et al., 2024), permutation testing for E/I ratio differences between alprazolam and placebo sessions (N = 45) was limited to 100 permutations due to the computational burden of Euler-based CMA-ES. With DELSSOME, substantially more permutations (e.g., 1000) would have been feasible, enabling more precise statistical inference even when sample size is small.

Beyond running existing studies faster and with more robustness, DELSSOME can potentially enable the community to answer new scientific questions that were previously impractical. In this study, we applied DELSSOME to individual-level modeling of 12,005 participants to estimate lifespan E/I trajectories. However, substantial opportunities remain to deepen our understanding of individual differences. For example, the FIC model bridges structural connectivity and emergent functional dynamics via local E/I mechanisms and long-range inter-regional interactions. Population-scale parameter estimation may further clarify how structural architecture constrains functional organization across individuals.

DELSSOME may also make higher-resolution modeling practical. In the current study, we used parcellations of 68 and 100 regions, but 1,000-region parcellations (Schaefer et al., 2018; Yan et al., 2023) may become tractable, capturing finer topography of E/I balance. Applying DELSSOME to a new parcellation requires regenerating training data (which scales linearly with the number of parcels) and adjusting input dimensionality of the model.

Nevertheless, our success across three large-scale circuit models suggests that the overall architecture and training procedure would remain unchanged.

### 3.4 Limitations & future work

A limitation of DELSSOME is the absence of formal uncertainty quantification. Although CMA-ES maintains a multivariate Gaussian search distribution, it reflects local sensitivity and parameter correlations arising from adaptive optimization rather than probabilistic inference, and should not be interpreted as a Bayesian posterior.

SBI methods naturally provide posterior distributions (Gonçalves et al., 2020; Tolley et al., 2024). However, because these models are typically only trained on simulations, the posteriors can be poorly calibrated on empirical data, leading to inaccurate parameter estimates (Fig. S4). Future work could integrate SBI ideas with DELSSOME to better characterize parameter uncertainty while remaining robust to the mismatch between simulations and empirical data.

Our model realism cost function relies on established FC and FCD metrics (Honey et al., 2009; Hansen et al., 2015; Pang et al., 2023). Because resting-state fMRI lacks temporal correspondence across sessions, individuals, and simulations, metrics comparing FCD matrices should be invariant to temporal ordering. This is achieved by collapsing FCD matrices into cumulative distributions and applying the KS statistic (Hansen et al., 2015; Pang et al., 2023; Zhang et al., 2024). However, this discards information about temporal structure, such as sequential state transitions. Developing richer, temporally invariant metrics that retain more FCD structure is an important direction for future work.

Finally, DELSSOME’s architecture is agnostic to the specific cost function being optimized. The framework could in principle be applied to other modalities – such as task-evoked activation patterns, magnetoencephalography, or electrophysiological recordings – by defining appropriate cost functions and simulating corresponding training data. Multi-objective fitting that incorporates spectral, temporal, or multimodal targets may further improve biological realism and broaden the scope of population-scale biophysical modeling.

### 3.5 Conclusions

In this work, we propose and validate the DELSSOME framework for speeding up the estimation of parameters in large-scale brain circuit models. The massive speedup provided by DELSSOME will enable the large-scale deployment of mechanistic models in population-level neuroscience.

## 4 Methods

### 4.1 Datasets

#### 4.1.1 Datasets demographics

We analysed resting-state fMRI data from multiple publicly available and locally acquired cohorts spanning early childhood to late adulthood. Unless stated otherwise, we applied cohort-specific quality control (QC) and inclusion/exclusion criteria, and report the sample sizes and demographics below. Site, scanner and scan protocol information are summarised in Supplementary Table S1.

Human Connectome Project young adults (HCP-YA) S1200. We used data from 1,029 participants from the HCP-YA WU–Minn S1200 release (Glasser et al., 2013; Van Essen et al., 2013). Participants were aged 22.0–37.0 years (mean 28.7, s.d. 3.7); 470 were male and 559 females.

Philadelphia Neurodevelopmental Cohort (PNC). Neuroimaging data were acquired from the PNC cohort of 1,601 youth (Satterthwaite et al., 2014; Calkins et al., 2015). A single resting-state fMRI scan was collected per participant. Consistent with our previous work (Zhang et al., 2024), we included 885 participants for analysis (age 8.2–23.0 years; mean 15.66, s.d. 3.36); 378 were male and 498 females.

HCP Development (HCP-D). We used resting-state fMRI data from the HCP-D dataset (Harms et al., 2018) and retained 612 participants out of 652 participants after QC (framewise displacement (FD) ≤ 0.2 and DVARS ≤ 75). Participants were aged 8.1–21.9 years (mean 14.7, s.d. 3.9); 284 were male and 328 female.

HCP Aging (HCP-A). We used resting-state fMRI data from HCP-A (Harms et al., 2018) and included 712 of 725 participants after excluding individuals with recorded age ≥ 100 years (ages in this range are top-coded and insufficiently precise for lifespan modelling). The retained sample spanned 36.0–89.8 years (mean 59.6, s.d. 14.9); 316 were male and 396 female.

Enhanced Nathan Kline Institute (eNKI). We used resting-state fMRI from eNKI (Nooner et al., 2012) and retained 689 participants of 1,333 by restricting to scans acquired with protocol R-mfMRI (TR = 1,400 ms; 2 mm isotropic voxels; 10 min duration). Participants were aged 6.0–85.0 years (mean 35.0, s.d. 19.1); 265 were male and 424 female.

Adolescent Brain Cognitive Development (ABCD). We analysed resting-state fMRI from 4,609 participants from the ABCD study (Casey et al., 2018; Jernigan et al., 2018; Volkow et al., 2018) after QC (FD ≤ 0.3; DVARS ≤ 50; boundary-based registration (BBR) ≥ 0.6). Participants were aged 8.9–13.8 years (mean 11.0, s.d. 1.2); 2,392 were male and 2,217 female.

GUSTO. We used resting-state fMRI from the GUSTO cohort (Soh et al., 2014) and retained 527 participants after QC. QC included mean FD ≤ 0.5, BBR ≥ 0.7, and at least 4 min of data remaining after censoring (censoring thresholds: FD 0.5 and DVARS 80). Participants were aged 4.5–11.3 years (mean 7.8, s.d. 2.3); 255 were male and 272 female.

The SINgapore GERiatric Intervention Study to Reduce Cognitive Decline and Physical Frailty (SINGER). We used resting-state fMRI from the SINGER cohort; preprocessing procedures followed our previous study (Ooi et al., 2025). From 898 participants, we retained 859 after excluding individuals with neurological disorders (Parkinson’s disease/other parkinsonism, seizures/epilepsy, traumatic brain injury with loss of consciousness ≥ 5 min or chronic deficits, and other neurological conditions), psychiatric disorders (active depression within the last 2 years and other psychiatric disorders), or a history of brain tumour. Participants were aged 60.1–80.7 years (mean 69.0, s.d. 4.8); 413 were male and 446 female.

SG70. We used resting-state fMRI from the SG70 dataset (n = 1,101). After removing participants with missing age information or missing imaging files, 942 participants remained. Participants were aged 68.8–80.8 years (mean 74.8, s.d. 2.6); 436 were male and 506 female.

SG-Lifespan (LIFE). We used resting-state fMRI from the LIFE dataset. Of 272 participants, 240 were retained after removing those with missing age information or missing imaging files. Participants were aged 21.0–73.0 years (mean 40.0, s.d. 14.9); 93 were male and 147 female.

Alzheimer’s Disease Neuroimaging Initiative (ADNI). We used resting-state fMRI from ADNI (Jack et al., 2008; Petersen et al., 2010) and retained 491 participants of 914 by restricting to healthy control individuals and retaining the baseline session only. Participants were aged 50.6–98.4 years (mean 72.6, s.d. 7.6); 205 were male and 286 female. Data used in the preparation of this article were obtained from the Alzheimer’s Disease Neuroimaging Initiative (ADNI) database (adni.loni.usc.edu). The ADNI was launched in 2003 as a public-private partnership, led by Principal Investigator Michael W. Weiner, MD. The primary goal of ADNI has been to test whether serial magnetic resonance imaging (MRI), positron emission tomography (PET), other biological markers, and clinical and neuropsychological assessment can be combined to measure the progression of mild cognitive impairment (MCI) and early Alzheimer’s disease (AD).

MACC Harmonization cohort. We used data from the MACC Harmonization dataset (Hilal et al., 2020) and retained 115 participants of 712 by restricting to healthy control individuals and retaining the baseline session only. Participants were aged 50.0–89.0 years (mean 68.6, s.d. 7.6); 52 were male and 63 female.

devCCNP. We used resting-state fMRI data from the devCCNP dataset (Zuo & Consortium, 2023) and included all 204 participants from the baseline session. Participants were aged 6.0–18.2 years (mean 10.6, s.d. 2.8); 112 were male and 92 female.

Transdiagnostic Connectome Project (TCP). We used resting-state fMRI from the TCP dataset (Chopra et al., 2025) and retained 91 participants of 245 by restricting to healthy control individuals and excluding one participant with non-binary gender to enable binary sex-stratified summaries. Participants were aged 18.1–68.3 years (mean 32.8, s.d. 13.1); 41 were male and 50 female.

#### 4.1.2 Preprocessing

For the HCP dataset, preprocessing of the HCP fMRI data is detailed in the HCP S1200 manual (Glasser et al., 2013; Van Essen et al., 2013). We utilized preprocessed (MSMAll) resting-fMRI data, already projected to fsLR surface space, denoised with ICA-FIX, and smoothed by 2 mm.

For the PNC dataset, preprocessing followed previous studies (Larsen et al., 2022; Zhang et al., 2024). The steps are as follows. (1) Slice time correction. (2) Motion correction. (3) Correcting for susceptibility-induced spatial distortion. (4) Alignment with structural image using boundary-based registration (Greve & Fischl, 2009). (5) Five aCompCor components (Behzadi et al., 2007; Chai et al., 2012), six motion parameters and their temporal derivatives were regressed from the functional data. (6) The data underwent bandpass filtering (0.009–0.08 Hz). (7) Lastly, the data were then projected onto fsLR surface space.

For the HCP-D dataset, the minimally processed fsLR surface data with MSMAll was used (Harms et al., 2018). We additionally processed the minimally processed data with the following steps. (1) Five CompCor components were extracted from cerebrospinal fluid (CSF) and white matter (WM) masks. In total, 17 regressors were jointly regressed from the BOLD time series, including 6 head motion parameters and their temporal derivatives, and the top 5 aCompCor components. (2) The data underwent bandpass filtering (0.01–0.1 Hz).

For the HCP-A dataset, the minimally processed fsLR surface data with MSMAll was used (Harms et al., 2018). We additionally processed the minimally processed data with the following steps. (1) Nuisance regression the same as that of HCP-D. (2) The data underwent bandpass filtering (0.01–0.1 Hz).

For the eNKI dataset (Nooner et al., 2012), we processed the functional data with the following steps. (1) Removal of the first four frames. (2) Slice time correction. (3) Motion correction. (4) Alignment with structural image using boundary-based registration. (5) Five aCompCor components, six motion parameters and their temporal derivatives were regressed from the functional data. (6) Lastly, the data were then projected onto FreeSurfer fsaverage6 surface space.

For the ABCD dataset, the minimally processed resting state scans were used (Hagler Jr et al., 2019). We additionally processed the minimally processed data with the following steps. (1) The functional images were aligned to the T1 images using boundary-based registration. (2) Respiratory pseudomotion filtering was performed by applying a bandstop filter of 0.31–0.43 Hz. (3) Five aCompCor components, six motion parameters and their temporal derivatives were regressed from the functional data. (4) The data underwent bandpass filtering (0.009–0.08 Hz). (5) Lastly, the data were projected onto FreeSurfer fsaverage6 surface space.

For the GUSTO dataset (Soh et al., 2014), we processed the functional data with the following steps. (1) Removal of the first four frames. (2) Slice time correction. (3) Motion correction. (4) Alignment with structural image using boundary-based registration. (5) Five aCompCor components, six motion parameters and their temporal derivatives were regressed from the functional data. (6) The data underwent bandpass filtering (0.009–0.08 Hz). (7) Lastly, the data were then projected onto FreeSurfer fsaverage6 surface space.

For the SG70 dataset, we processed the functional data with the following steps. (1) Removal of the first four frames. (2) Slice time correction. (3) Motion correction. (4) Correcting for susceptibility-induced spatial distortion. (5) Multi-echo denoising (DuPre et al., 2021). (6) Alignment with structural image using boundary-based registration. (7) Five aCompCor components, six motion parameters and their temporal derivatives were regressed from the functional data. (8) The data underwent bandpass filtering (0.009–0.08 Hz). (9) Lastly, the data were then projected onto FreeSurfer fsaverage6 surface space.

For the SINGER dataset, we processed the functional data with the following steps (Ooi et al., 2025). (1) Removal of the first four frames. (2) Slice time correction. (3) Motion correction. (4) Correcting for susceptibility-induced spatial distortion. (5) Multi-echo denoising. (6) Alignment with structural image using boundary-based registration. (7) Five aCompCor components, six motion parameters and their temporal derivatives were regressed from the functional data. (8) The data underwent bandpass filtering (0.009–0.08 Hz). (9) Lastly, the data were then projected onto FreeSurfer fsaverage6 surface space.

For the LIFE dataset, we processed the functional data with the following steps. (1) Removal of the first four frames. (2) Slice time correction. (3) Motion correction. (4) Correcting for susceptibility-induced spatial distortion. (5) Multi-echo denoising. (6) Alignment with structural image using boundary-based registration. (7) Five aCompCor components, six motion parameters and their temporal derivatives were regressed from the functional data. (8) The data underwent bandpass filtering (0.009–0.08 Hz). (9) Lastly, the data were then projected onto FreeSurfer fsaverage6 surface space.

For the ADNI dataset (Jack et al., 2008; Petersen et al., 2010), we processed the functional data with the following steps. (1) Slice time correction. (2) Motion correction. (3) Alignment with structural image using boundary-based registration. (4) Five aCompCor components, six motion parameters, and their temporal derivatives were regressed from the functional data. (5) Lastly, the data were then projected onto FreeSurfer fsaverage6 surface space.

For the MACC Harmonization dataset (Hilal et al., 2020), we processed the functional data with the following steps. (1) Removal of the first four frames. (2) Slice time correction. (3) Motion correction. (4) Alignment with structural image using boundary-based registration. (5) Five aCompCor components, six motion parameters and their temporal derivatives were regressed from the functional data. (6) Lastly, the data were then projected onto FreeSurfer fsaverage6 surface space.

For the devCCNP dataset (Zuo & Consortium, 2023), we processed the functional data with the following steps. (1) Removal of the first four frames. (2) Slice time correction. (3) Motion correction. (4) Alignment with structural image using boundary-based registration. (5) Five aCompCor components, six motion parameters and their temporal derivatives were regressed from the functional data. (6) The data underwent bandpass filtering (0.009–0.08 Hz). (7) Lastly, the data were then projected onto FreeSurfer fsaverage6 surface space.

For the TCP dataset (Chopra et al., 2025), we processed the functional data with the following steps. (1) Removal of the first four frames. (2) Slice time correction. (3) Motion correction. (4) Correcting for susceptibility-induced spatial distortion. (5) Alignment with structural image using boundary-based registration. (6) Five aCompCor components, six motion parameters and their temporal derivatives were regressed from the functional data. (7) The data underwent bandpass filtering (0.009–0.08 Hz). (8) Lastly, the data were then projected onto FreeSurfer fsaverage6 surface space.

#### 4.1.3 The computation of FC, FCD and SC

Let us first consider the HCP dataset. For each run of each participant, the fMRI time courses were averaged within each Desikan–Killiany region of interest (ROI; Desikan et al., 2006) to create a 68 × 1200 matrix. 68 × 68 functional connectivity (FC) matrices were then computed by correlating the time courses among all pairs of time courses.

Functional connectivity dynamics (FCD) were computed by defining a 60-second window (equivalent to 83 time points or TRs) and sliding the window frame by frame, resulting in 1118 sliding windows (Zalesky et al., 2014; Hansen et al., 2015; Leonardi & Van De Ville, 2015). FC was computed within each sliding window for each participant’s run. Each sliding window FC matrix was then vectorized by only considering the upper triangular entries. The vectorized FCs were correlated, resulting in a 1118 × 1118 FCD matrix for each run of each participant.

FC and FCD for the other datasets were computed in the same manner as the HCP dataset. FCD was always computed using a 60-second window. However, the TR and scan length differs for each dataset, so the final dimensions of the FCD matrices will vary across datasets.

For diffusion MRI, the processing begins with *b*0 intensity normalization, followed by the calculation of susceptibility-induced B_0_ field deviations using *b*0 images acquired in both phase-encoding directions. The full diffusion timeseries from both phase-encoding directions is then processed with the “eddy” tool, which corrects for eddy current distortions and subject motion. Gradient distortion is subsequently corrected, and the *b*0 image is aligned to the T1-weighted image using boundary-based registration (BBR). The corrected diffusion data output from the “eddy” step are resampled into the native structural space at 1.25 mm resolution and masked. Additionally, diffusion directions and gradient deviation estimates are appropriately rotated and registered into the structural space (Glasser et al., 2013). Probabilistic tractography with the Anatomically-Constrained Tractography (ACT) (Smith et al., 2012) was performed on fiber orientation distribution (FOD) images in each participant, using the second-order integration over fiber orientation distribution (iFOD2) algorithm provided by MRtrix3 (Tournier et al., 2010, 2019). 5,000,000 streamlines were sampled to generate tractograms, which were then filtered using Spherical-deconvolution informed filtering of tracks (SIFT2; Smith et al., 2015). This process generated a structural connectivity (SC) matrix for each participant, where each entry represented the weighted sum of the streamlines, approximating the mean fiber density (Smith et al., 2022) between two Desikan-Killiany ROIs.

### 4.2 DELSSOME neural network architecture

To avoid computationally intensive numerical integration (Figure 1c) in the CMA-ES evaluation, we proposed the DELSSOME (Deep Learning for Surrogate Statistics Optimization in Mean Field Modeling) to directly predict the FC+FCD cost without numerical integration. DELSSOME utilized the transformer architecture (Fig. 9; Vaswani et al., 2017). The inputs to DELSSOME were the FIC parameters, the SC matrix, the FC matrix, and the FCD-CDF. We note that FCD-CDF was converted to a probability distribution function (i.e., FCD-PDF) before input to DELSSOME.

**Figure 9.**
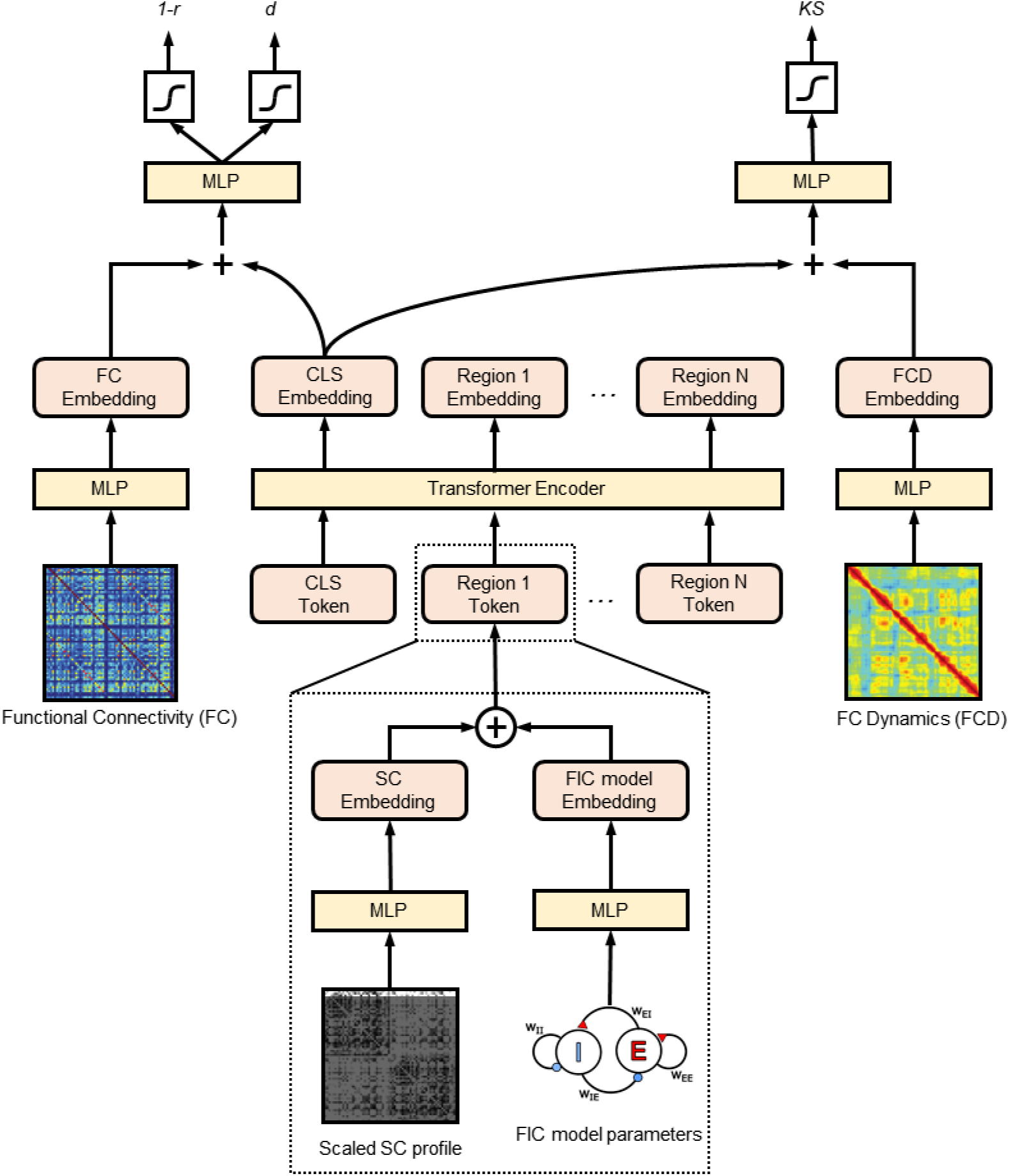
DELSSOME neural network architecture. Overview of the DELSSOME FC+FCD cost predictor. The inputs to this model are FIC model parameters, SC, empirical FC matrix and empirical FCD probability distribution function (FCD-PDF). The FIC parameters and SC profile of each region of interest (ROI) were passed through separate multi-layer perceptrons (MLPs) and then concatenated to generate a vector representing sufficient statistics of the ROI. The sequence of ROI tokens together with a prepended [CLS] token were passed through a transformer encoder. The hidden state corresponding to the [CLS] token was extracted as the FIC model embedding. The FC and FCD-PDF were also passed through separate MLPs to generate FC and FCD embeddings. The FC and FIC model embeddings were added together and passed through another MLP to predict FC costs *1-r* and *d*. The FCD and FIC model embeddings were added together and passed through another MLP to predict FCD cost *KS*.

The goal of the transformer encoder module (Fig. 9) was to encode the dynamics of the FIC model. Therefore, the input to the encoder module were the FIC parameters and SC matrix. More specifically, for each Desikan-Killiany ROI, the region’s w_EE_, w_EI_ and σ were mapped to a d/2-dimensional representation via a multi-layer perceptron (MLP). For each ROI, the region’s SC profile (i.e., corresponding row of the SC matrix) was scaled by the global coupling parameter G, and mapped to a d/2-dimensional representation through another MLP. These two d/2-dimensional vectors were concatenated to form a d-dimensional token embedding. Since there were 68 Desikan-Killiany ROIs, this process yielded a sequence of 68 d-dimensional tokens. In addition, a [CLS] token of dimensionality d was prepended to the sequence of 68 tokens, and the entire sequence of 69 tokens was passed to a transformer encoder (Devlin et al., 2019). The hidden state corresponding to the [CLS] token was extracted as the FIC model embedding.

Since the FC cost involved comparing the simulated FC and empirical FC, so the empirical FC matrix should be another input to the DELSSOME FC+FCD cost predictor. The empirical FC matrix was fed through an MLP, resulting in an FC embedding. The FC embedding was added with the FIC model embedding (from the CLS token), and fed through a final MLP to predict the FC costs *1-r* and *d*.

Similarly, since the FCD cost involved comparing the simulated FCD and empirical FCD, so the empirical FCD should also be an input to the DELSSOME FC+FCD cost predictor. The empirical FCD-PDF was fed through an MLP, resulting in an FCD embedding. The FCD embedding was added with the FIC model embedding (from the CLS token), and fed through a final MLP to predict the FCD cost *KS*. More details of the DELSSOME FC+FCD cost predictor and the hyperparameters are found in Supplementary Methods S5.

In an earlier version of DELSSOME, a within-range classifier was used to identify parameter sets producing firing rates within a predefined physiological range (2.7–3.3 Hz) before cost evaluation. However, subsequent analyses suggest that the quality of the FIC parameters was independent of the firing rate (Fig. S10). In other words, the FC+FCD cost can be low even when firing rate was as low as 2 Hz. Therefore, for simplicity, the within-range classifier was removed from the final DELSSOME framework.

Finally, although we followed previous studies (Brunel & Wang, 2001; Wong & Wang, 2006; Deco et al., 2014) to set the FIC excitatory firing rate to be 3Hz (Supplementary Methods S1), we note that another plausible firing rate is 0.41Hz (De Kock & Sakmann, 2008). When the firing rate was set to be 0.41Hz, DELSSOME CMA-ES again achieved ~50× speed up over Euler CMA-ES, with comparable model parameter estimates (Fig. S11), suggesting that DELSSOME’s effectiveness is not contingent on the specific firing-rate assumption. Intriguingly, when the firing rate was set to be 0.41Hz, the FC+FCD cost was substantially worse than when the firing rate was set to be 3Hz (Fig. 3c vs Fig. S11b), suggesting that the 3Hz assumption led to more realistic fMRI.

### 4.3 Training & evaluation of DELSSOME cost predictor

To train and evaluate the DELSSOME FC+FCD cost predictor, we divided the HCP dataset in training (N = 680), validation (N = 180) and test (N = 169) sets (Figure 10). Most prior studies have simulated large-scale circuit models at the group level (Demirtaş et al., 2019; Müller et al., 2023; Pang et al., 2023), in part due to the substantial computational demands of individual-level modeling. Accordingly, we first focused on group-level analyses (Figs. 2 to 5) before extending the framework to individual-level analyses (Figs. 6 to 8). To avoid test-set leakage, we generated group-level SC, FC, and FCD-CDF separately within the training, validation, and test sets.

**Figure 10.**
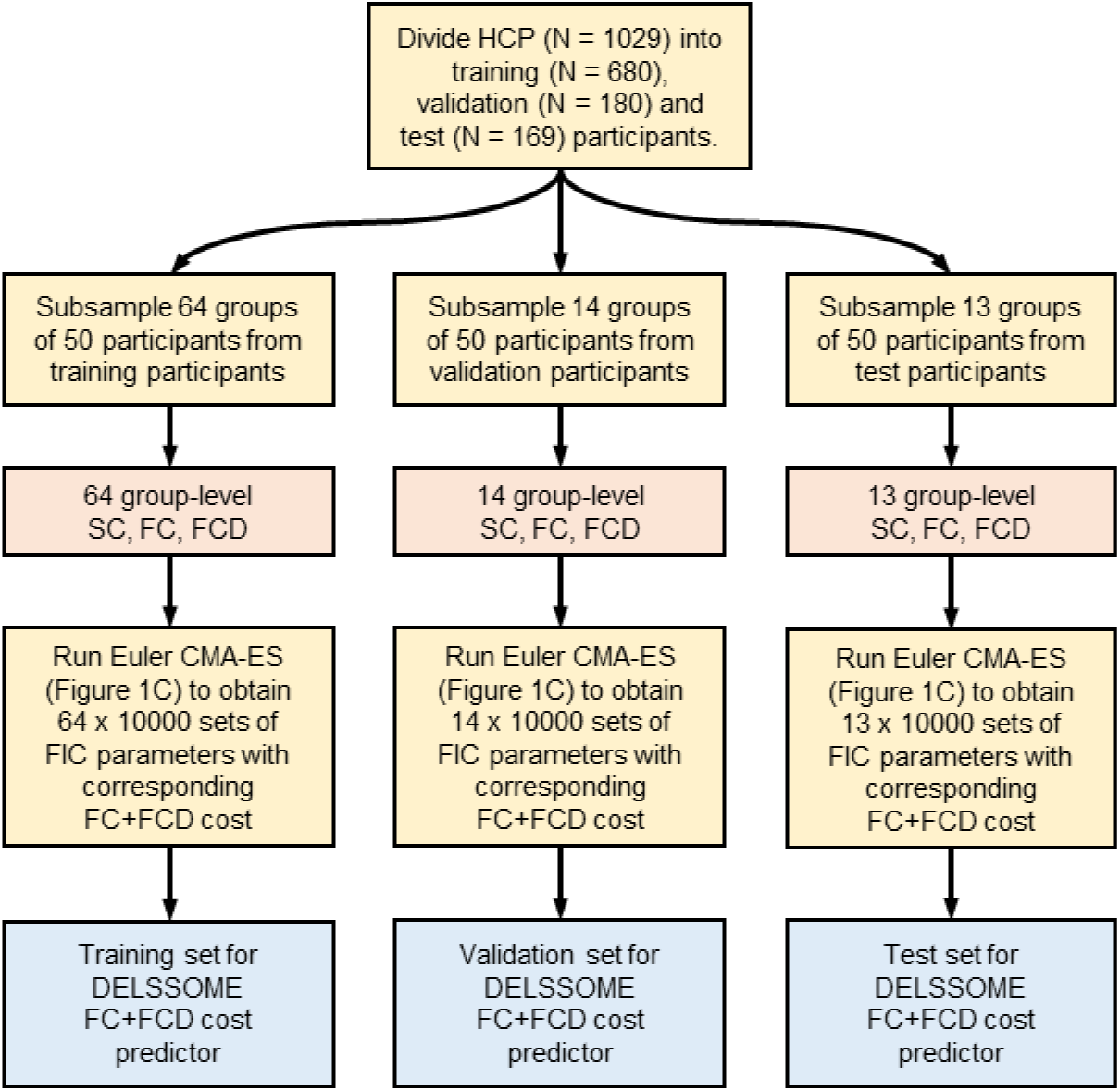
Data generation for training and evaluating DELSSOME FC+FCD cost predictor.

The 680-participant training set permits at most 13 non-overlapping groups of 50 participants. To augment the effective sample size and capture greater variability in group-level connectivity patterns, we therefore repeatedly sampled 50 participants within the training set to compute group-average SC, FC and FCD-CDF. The sampling was performed 64 times resulting in 64 group-average SC, FC and FCD-CDF in the training set. The same procedure was repeated in the validation and test sets separately, yielding 14 group-level SC, FC and FCD-CDF for the validation set, as well as 13 SC, FC and FCD-CDF for the test set.

For each triplet of group-level SC, FC and FCD-CDF, we ran CMA-ES with Euler integration for 100 epochs. Euler integration was performed with time step of 6 ms. Since each CMA-ES epoch employed 100 children, this process generated 10,000 sets of FIC (or MFM or Hopf model) parameters with corresponding ground truth FC+FCD cost, which could be used to train and evaluate the DELSSOME model. Since there were 64, 14 and 13 group-level SC, FC and FCD in the training, validation and test sets respectively, this yielded 640,000, 140,000 and 130,000 training, validation and test samples respectively.

The 640,000 training samples were used to train the DELSSOME FC+FCD cost predictor, while the 140,000 validation samples were used to tune the hyperparameters. More details of the hyperparameter tuning and final set of hyperparameters are found in Supplementary Method S5. The final trained DELSSOME cost predictor was then evaluated with the 130,000 test samples.

The details of dataset generation and training procedure of individual-level DELSSOME cost predictor can be found in Supplementary Methods S6.

### 4.4 Computational cost in training DELSSOME models

Both Euler CMA-ES and DELSSOME CMA-ES are coded in python and run on the same hardware: Intel(R) Xeon(R) Gold 6230 CPU @ 2.10GHz. While the application of trained DELSSOME models is very fast (as reported in the main text), there is significant cost in generating DELSSOME training data and training DELSSOME models.

For the FIC, MFM, and Hopf models, the generation of DELSSOME training, validation and test data (Section 4.3) required 34, 17 and 111 hours of compute time, respectively, using 91 CPU cores in parallel, running Euler CMA-ES. Training the corresponding DELSSOME models required an additional 3 hours of compute on 4× RTX 3090 GPU cores across all model setups. No model-specific hyperparameter tuning was performed; the same architecture and training configuration were used for all three models.

For the individual-level DELSSOME model, generating the training, validation and test data required approximately 90 hours on 200 CPU cores, running Euler CMA-ES. Hyperparameter tuning and model training were conducted using 50 Optuna trials, requiring 65 hours on a single RTX 3090 GPU.

### 4.5 Ethics and data availability

Use of de-identified data from all used datasets is approved by the National University of Singapore (NUS) Institutional Review Board (IRB).

The raw data for HCP (https://www.humanconnectome.org/), ABCD (https://abcdstudy.org/), TCP (https://openneuro.org/datasets/ds005237 and https://nda.nih.gov/edit_collection.html?id=3552), ADNI (https://ida.loni.usc.edu/), enhanced Nathan Kline Institute-Rockland sample (eNKI; https://fcon_1000.projects.nitrc.org/indi/enhanced/), HCP-D and HCP-A (https://www.humanconnectome.org/study/hcp-lifespan-development/data-releases) are publicly available, which can be accessed via data use agreements according to each dataset’s requirements. The PNC dataset is publicly available in the Database of Genotypes and Phenotypes (dbGaP accession phs000607.v3.p2). The SINGER, MACC, GUSTO, SG70, LIFE datasets can be obtained through each dataset’s data-transfer agreement. The devCCNP dataset can be accessed by completing the application file Data Use Agreement on CCNP (DUA-CCNP) at http://deepneuro.bnu.edu.cn/?p=163 and have it reviewed and approved by the Chinese Color Nest Consortium (CCNC).

### 4.6 Code availability

Code for this study, together with pretrained DELSSOME models can be found here (GITHUB_LINK). Co-authors (TZ and FT) reviewed the code before merging into the GitHub repository to reduce the chance of coding errors.

## Acknowledgements

Our research is supported by the NUS Yong Loo Lin School of Medicine (NUHSRO/2020/124/TMR/LOA), the Singapore National Medical Research Council (NMRC) LCG (OFLCG19May-0035), NMRC CTG-IIT (CTGIIT23jan-0001), NMRC OF-IRG (OFIRG24jan-0006; OFIRG24jul-0049), NMRC STaR (STaR20nov-0003), Singapore Ministry of Health (MOH) Centre Grant (CG21APR1009), the United States National Institutes of Health (R01MH133334 & 2R01MH120080, R00MH127293, R01MH120080, R01MH112847, U24 NS130411, R01MH113550, R01EB022573) and the Singapore National Research Foundation (NRF) Investigatorship (NRFI10-2024-0014). Additional support was provided by the Penn/CHOP Lifespan Brain Institute and the Penn AI2D center. Any opinions, findings and conclusions or recommendations expressed in this material are those of the authors and do not reflect the views of the funders.

Data were in part provided by the Human Connectome Project, WU-Minn Consortium (Principal Investigators: David Van Essen and Kamil Ugurbil; 1U54MH091657) funded by the 16 NIH Institutes and Centers that support the NIH Blueprint for Neuroscience Research; and by the McDonnell Center for Systems Neuroscience at Washington University. In RIE2025, the GUSTO dataset was supported by funding from the NRF’s Human Health and Potential (HHP) Domain, under the Human Potential Programme. The PNC was funded via RC2 grants from the National Institute of Mental Health: MH089983 and MH089924. The SG70 project was supported by Singapore National Medical Research Council [CSA-SI (MOH-000434)] and Yong Loo Lin School of Medicine, National University of Singapore. Additionally, we acknowledge Benjamin Yi Liang Goh for his processing of SG70 dataset. The SINGER project was supported by funding from the National Medical Research Council of Singapore (MOH000500).

Data used in the preparation of this article were obtained from the Adolescent Brain Cognitive Development^SM^ (ABCD) Study (https://abcdstudy.org), held in the NIMH Data Archive (NDA). This is a multisite, longitudinal study designed to recruit more than 10,000 children age 9-10 and follow them over 10 years into early adulthood. The ABCD Study® is supported by the National Institutes of Health and additional federal partners under award numbers U01DA041048, U01DA050989, U01DA051016, U01DA041022, U01DA051018, U01DA051037, U01DA050987, U01DA041174, U01DA041106, U01DA041117, U01DA041028, U01DA041134, U01DA050988, U01DA051039, U01DA041156, U01DA041025, U01DA041120, U01DA051038, U01DA041148, U01DA041093, U01DA041089, U24DA041123, U24DA041147. A full list of supporters is available at https://abcdstudy.org/federal-partners.html. A listing of participating sites and a complete listing of the study investigators can be found at https://abcdstudy.org/consortium_members/. ABCD consortium investigators designed and implemented the study and/or provided data but did not necessarily participate in the analysis or writing of this report. This manuscript reflects the views of the authors and may not reflect the opinions or views of the NIH or ABCD consortium investigators. ABCD data repository grows and changes over time. The ABCD data used in this report came from the NIMH Data Archive and were drawn from Release 4.0 (DOI: <u>10.15154/1523041</u>) and Release 5.1 (DOI: <u>10.15154/z563-zd24</u>)

Data collection and sharing for the Alzheimer’s Disease Neuroimaging Initiative (ADNI) is funded by the National Institute on Aging (National Institutes of Health Grant U19AG024904). The grantee organization is the Northern California Institute for Research and Education. In the past, ADNI has also received funding from the National Institute of Biomedical Imaging and Bioengineering, the Canadian Institutes of Health Research, and private sector contributions through the Foundation for the National Institutes of Health (FNIH) including generous contributions from the following: AbbVie, Alzheimer’s Association; Alzheimer’s Drug Discovery Foundation; Araclon Biotech; BioClinica, Inc.; Biogen; BristolMyers Squibb Company; CereSpir, Inc.; Cogstate; Eisai Inc.; Elan Pharmaceuticals, Inc.; Eli Lilly and Company; EuroImmun; F. Hoffmann-La Roche Ltd and its affiliated company Genentech, Inc.; Fujirebio; GE Healthcare; IXICO Ltd.; Janssen Alzheimer Immunotherapy Research & Development, LLC.; Johnson & Johnson Pharmaceutical Research & Development LLC.; Lumosity; Lundbeck; Merck & Co., Inc.; Meso Scale Diagnostics, LLC.; NeuroRx Research; Neurotrack Technologies; Novartis Pharmaceuticals Corporation; Pfizer Inc.; Piramal Imaging; Servier; Takeda Pharmaceutical Company; and Transition Therapeutics. Data used in preparation of this article were obtained from the Alzheimer’s Disease Neuroimaging Initiative (ADNI) database (adni.loni.usc.edu). As such, the investigators within the ADNI contributed to the design and implementation of ADNI and/or provided data but did not participate in the analysis or writing of this report. A complete listing of ADNI investigators can be found at: http://adni.loni.usc.edu/wp-content/uploads/how_to_apply/ADNI_Acknowledgement_List.pdf.

HCP-Development data were used. Research reported in this publication was supported by the National Institute Of Mental Health of the National Institutes of Health under Award Number U01MH109589 and by funds provided by the McDonnell Center for Systems Neuroscience at Washington University in St. Louis. The HCP-Development 2.0 Release data used in this report came from DOI: 10.15154/1520708. HCP-Aging data were used. Research reported in this publication was supported by the National Institute On Aging of the National Institutes of Health under Award Number U01AG052564 and by funds provided by the McDonnell Center for Systems Neuroscience at Washington University in St. Louis. The HCP-Aging 2.0 Release data used in this report came from DOI: 10.15154/1520707. The content is solely the responsibility of the authors and does not necessarily represent the official views of the National Institutes of Health.

## References

Anticevic, A., & Lisman, J. (2017). How can global alteration of excitation/inhibition balance lead to the local dysfunctions that underlie schizophrenia? Biological Psychiatry, 81(10), 818–820.

Behzadi, Y., Restom, K., Liau, J., & Liu, T. T. (2007). A component based noise correction method (CompCor) for BOLD and perfusion based fMRI. Neuroimage, 37(1), 90–101.

Bethlehem, R. A., Seidlitz, J., White, S. R., Vogel, J. W., Anderson, K. M., Adamson, C., Adler, S., Alexopoulos, G. S., Anagnostou, E., & Areces-Gonzalez, A. (2022). Brain charts for the human lifespan. Nature, 604(7906), 525–533.

Boelts, J., Deistler, M., Gloeckler, M., Tejero-Cantero, Á., Lueckmann, J.-M., Moss, G., Steinbach, P., Moreau, T., Muratore, F., Linhart, J., Durkan, C., Vetter, J., Miller, B. K., Herold, M., Ziaeemehr, A., Pals, M., Gruner, T., Bischoff, S., Krouglova, N., … Macke, J. H. (2025). *sbi reloaded: A toolkit for simulation-based inference workflows* (arXiv:2411.17337). arXiv. 10.48550/arXiv.2411.17337

Breakspear, M., Jirsa, V., & Deco, G. (2010). Computational models of the brain: From structure to function. In Neuroimage (Vol. 52, Issue 3, pp. 727–730). Elsevier. https://www.sciencedirect.com/science/article/pii/S1053811910007949

Brunel, N., & Wang, X.-J. (2001). Effects of Neuromodulation in a Cortical Network Model of Object Working Memory Dominated by Recurrent Inhibition. Journal of Computational Neuroscience, 11(1), 63–85. 10.1023/A:1011204814320

Calkins, M. E., Merikangas, K. R., Moore, T. M., Burstein, M., Behr, M. A., Satterthwaite, T. D., Ruparel, K., Wolf, D. H., Roalf, D. R., Mentch, F. D., Qiu, H., Chiavacci, R., Connolly, J. J., Sleiman, P. M. A., Gur, R. C., Hakonarson, H., & Gur, R. E. (2015). The Philadelphia Neurodevelopmental Cohort: Constructing a deep phenotyping collaborative. Journal of Child Psychology and Psychiatry, 56(12), 1356–1369. 10.1111/jcpp.12416

Casey, B. J., Cannonier, T., Conley, M. I., Cohen, A. O., Barch, D. M., Heitzeg, M. M., Soules, M. E., Teslovich, T., Dellarco, D. V., & Garavan, H. (2018). The adolescent brain cognitive development (ABCD) study: Imaging acquisition across 21 sites. Developmental Cognitive Neuroscience, 32, 43–54.

Caspary, D. M., Hughes, L. F., & Ling, L. L. (2013). Age-related GABAA receptor changes in rat auditory cortex. Neurobiology of Aging, 34(5), 1486–1496.

Chai, X. J., Castañón, A. N., Öngür, D., & Whitfield-Gabrieli, S. (2012). Anticorrelations in resting state networks without global signal regression. Neuroimage, 59(2), 1420–1428.

Chamberlain, J. D., Gagnon, H., Lalwani, P., Cassady, K. E., Simmonite, M., Seidler, R. D., Taylor, S. F., Weissman, D. H., Park, D. C., & Polk, T. A. (2021). GABA levels in ventral visual cortex decline with age and are associated with neural distinctiveness. Neurobiology of Aging, 102, 170–177.

Chopra, S., Cocuzza, C. V., Lawhead, C., Ricard, J. A., Labache, L., Patrick, L. M., Kumar, P., Rubenstein, A., Moses, J., & Chen, L. (2025). The Transdiagnostic Connectome Project: An open dataset for studying brain-behavior relationships in psychiatry. Scientific Data, 12(1), 923.

Cranmer, K., Brehmer, J., & Louppe, G. (2020). The frontier of simulation-based inference. Proceedings of the National Academy of Sciences, 117(48), 30055–30062. 10.1073/pnas.1912789117

De Kock, C. P. J., & Sakmann, B. (2008). High frequency action potential bursts (≥ 100 Hz) in L2/3 and L5B thick tufted neurons in anaesthetized and awake rat primary somatosensory cortex. The Journal of Physiology, 586(14), 3353–3364. 10.1113/jphysiol.2008.155580

Deco, G., Kringelbach, M. L., Arnatkeviciute, A., Oldham, S., Sabaroedin, K., Rogasch, N. C., Aquino, K. M., & Fornito, A. (2021). Dynamical consequences of regional heterogeneity in the brain’s transcriptional landscape. Science Advances, 7(29), eabf4752. 10.1126/sciadv.abf4752

Deco, G., Ponce-Alvarez, A., Hagmann, P., Romani, G. L., Mantini, D., & Corbetta, M. (2014). How Local Excitation-Inhibition Ratio Impacts the Whole Brain Dynamics. Journal of Neuroscience, 34(23), 7886–7898. 10.1523/JNEUROSCI.5068-13.2014

Deco, G., Ponce-Alvarez, A., Mantini, D., Romani, G. L., Hagmann, P., & Corbetta, M. (2013). Resting-state functional connectivity emerges from structurally and dynamically shaped slow linear fluctuations. Journal of Neuroscience, 33(27), 11239–11252.

Deistler, M., Boelts, J., Steinbach, P., Moss, G., Moreau, T., Gloeckler, M., Rodrigues, P. L. C., Linhart, J., Lappalainen, J. K., Miller, B. K., Gonçalves, P. J., Lueckmann, J.-M., Schröder, C., & Macke, J. H. (2025). *Simulation-Based Inference: A Practical Guide* (arXiv:2508.12939). arXiv. 10.48550/arXiv.2508.12939

Deistler, M., Kadhim, K. L., Pals, M., Beck, J., Huang, Z., Gloeckler, M., Lappalainen, J. K., Schröder, C., Berens, P., & Gonçalves, P. J. (2025). Jaxley: Differentiable simulation enables large-scale training of detailed biophysical models of neural dynamics. Nature Methods, 1–9.

Demirtaş, M., Burt, J. B., Helmer, M., Ji, J. L., Adkinson, B. D., Glasser, M. F., Van Essen, D. C., Sotiropoulos, S. N., Anticevic, A., & Murray, J. D. (2019). Hierarchical heterogeneity across human cortex shapes large-scale neural dynamics. Neuron, 101(6), 1181–1194.

Desikan, R. S., Ségonne, F., Fischl, B., Quinn, B. T., Dickerson, B. C., Blacker, D., Buckner, R. L., Dale, A. M., Maguire, R. P., & Hyman, B. T. (2006). An automated labeling system for subdividing the human cerebral cortex on MRI scans into gyral based regions of interest. Neuroimage, 31(3), 968–980.

Devlin, J., Chang, M.-W., Lee, K., & Toutanova, K. (2019). *BERT: Pre-training of Deep Bidirectional Transformers for Language Understanding* (arXiv:1810.04805). arXiv. 10.48550/arXiv.1810.04805

DuPre, E., Salo, T., Ahmed, Z., Bandettini, P., Bottenhorn, K., Caballero-Gaudes, C., Dowdle, L., Gonzalez-Castillo, J., Heunis, S., Kundu, P., Laird, A., Markello, R., Markiewicz, C., Moia, S., Staden, I., Teves, J., Uruñuela, E., Vaziri-Pashkam, M., Whitaker, K., & Handwerker, D. (2021). TE-dependent analysis of multi-echo fMRI with tedana. Journal of Open Source Software, 6(66), 3669. 10.21105/joss.03669

Froudist-Walsh, S., Bliss, D. P., Ding, X., Rapan, L., Niu, M., Knoblauch, K., Zilles, K., Kennedy, H., Palomero-Gallagher, N., & Wang, X.-J. (2021). A dopamine gradient controls access to distributed working memory in the large-scale monkey cortex. Neuron, 109(21), 3500–3520.

Gao, F., Edden, R. A., Li, M., Puts, N. A., Wang, G., Liu, C., Zhao, B., Wang, H., Bai, X., & Zhao, C. (2013). Edited magnetic resonance spectroscopy detects an age-related decline in brain GABA levels. Neuroimage, 78, 75–82.

Ghafourpour, L., Duruisseaux, V., Tolooshams, B., Wong, P. H., Anastassiou, C. A., & Anandkumar, A. (2025). *NOBLE -- Neural Operator with Biologically-informed Latent Embeddings to Capture Experimental Variability in Biological Neuron Models* (arXiv:2506.04536). arXiv. 10.48550/arXiv.2506.04536

Glasser, M. F., Sotiropoulos, S. N., Wilson, J. A., Coalson, T. S., Fischl, B., Andersson, J. L., Xu, J., Jbabdi, S., Webster, M., & Polimeni, J. R. (2013). The minimal preprocessing pipelines for the Human Connectome Project. Neuroimage, 80, 105–124.

Gonçalves, P. J., Lueckmann, J.-M., Deistler, M., Nonnenmacher, M., Öcal, K., Bassetto, G., Chintaluri, C., Podlaski, W. F., Haddad, S. A., & Vogels, T. P. (2020). Training deep neural density estimators to identify mechanistic models of neural dynamics. Elife, 9, e56261.

Greve, D. N., & Fischl, B. (2009). Accurate and robust brain image alignment using boundary-based registration. Neuroimage, 48(1), 63–72.

Hagler Jr, D. J., Hatton, S., Cornejo, M. D., Makowski, C., Fair, D. A., Dick, A. S., Sutherland, M. T., Casey, B. J., Barch, D. M., & Harms, M. P. (2019). Image processing and analysis methods for the Adolescent Brain Cognitive Development Study. Neuroimage, 202, 116091.

Hansen, E. C., Battaglia, D., Spiegler, A., Deco, G., & Jirsa, V. K. (2015). Functional connectivity dynamics: Modeling the switching behavior of the resting state. Neuroimage, 105, 525–535.

Hansen, N. (2006). The CMA Evolution Strategy: A Comparing Review. In J. A. Lozano, P. Larrañaga, I. Inza, & E. Bengoetxea (Eds.), Towards a New Evolutionary Computation (Vol. 192, pp. 75–102). Springer Berlin Heidelberg. 10.1007/3-540-32494-1_4

Harms, M. P., Somerville, L. H., Ances, B. M., Andersson, J., Barch, D. M., Bastiani, M., Bookheimer, S. Y., Brown, T. B., Buckner, R. L., & Burgess, G. C. (2018). Extending the Human Connectome Project across ages: Imaging protocols for the Lifespan Development and Aging projects. Neuroimage, 183, 972–984.

Hilal, S., Tan, C. S., Van Veluw, S. J., Xu, X., Vrooman, H., Tan, B. Y., Venketasubramanian, N., Biessels, G. J., & Chen, C. (2020). Cortical cerebral microinfarcts predict cognitive decline in memory clinic patients. Journal of Cerebral Blood Flow & Metabolism, 40(1), 44–53. 10.1177/0271678X19835565

Hodgkin, A. L., & Huxley, A. F. (1952). A quantitative description of membrane current and its application to conduction and excitation in nerve. The Journal of Physiology, 117(4), 500.

Honey, C. J., Sporns, O., Cammoun, L., Gigandet, X., Thiran, J. P., Meuli, R., & Hagmann, P. (2009). Predicting human resting-state functional connectivity from structural connectivity. Proceedings of the National Academy of Sciences, 106(6), 2035–2040. 10.1073/pnas.0811168106

Jack, C. R., Bernstein, M. A., Fox, N. C., Thompson, P., Alexander, G., Harvey, D., Borowski, B., Britson, P. J., L. Whitwell, J., Ward, C., Dale, A. M., Felmlee, J. P., Gunter, J. L., Hill, D. L. G., Killiany, R., Schuff, N., Fox-Bosetti, S., Lin, C., Studholme, C., … Weiner, M. W. (2008). The Alzheimer’s disease neuroimaging initiative (ADNI): MRI methods. Journal of Magnetic Resonance Imaging, 27(4), 685–691. 10.1002/jmri.21049

Jernigan, T. L., Brown, S. A., & Dowling, G. J. (2018). The adolescent brain cognitive development study. Journal of Research on Adolescence: The Official Journal of the Society for Research on Adolescence, 28(1), 154.

Kong, X., Kong, R., Orban, C., Wang, P., Zhang, S., Anderson, K., Holmes, A., Murray, J. D., Deco, G., van den Heuvel, M., & Yeo, B. T. T. (2021). Sensory-motor cortices shape functional connectivity dynamics in the human brain. Nature Communications, 12(1), Article 1. 10.1038/s41467-021-26704-y

Kovachki, N., Li, Z., Liu, B., Azizzadenesheli, K., Bhattacharya, K., Stuart, A., & Anandkumar, A. (2023). Neural operator: Learning maps between function spaces with applications to pdes. Journal of Machine Learning Research, 24(89), 1–97.

Kringelbach, M. L., Cruzat, J., Cabral, J., Knudsen, G. M., Carhart-Harris, R., Whybrow, P. C., Logothetis, N. K., & Deco, G. (2020). Dynamic coupling of whole-brain neuronal and neurotransmitter systems. Proceedings of the National Academy of Sciences, 117(17), 9566–9576. 10.1073/pnas.1921475117

Lam, N. H., Borduqui, T., Hallak, J., Roque, A., Anticevic, A., Krystal, J. H., Wang, X.-J., & Murray, J. D. (2022). Effects of altered excitation-inhibition balance on decision making in a cortical circuit model. Journal of Neuroscience, 42(6), 1035–1053.

Larsen, B., Cui, Z., Adebimpe, A., Pines, A., Alexander-Bloch, A., Bertolero, M., Calkins, M. E., Gur, R. E., Gur, R. C., Mahadevan, A. S., Moore, T. M., Roalf, D. R., Seidlitz, J., Sydnor, V. J., Wolf, D. H., & Satterthwaite, T. D. (2022). A developmental reduction of the excitation:inhibition ratio in association cortex during adolescence. Science Advances, 8(5), eabj8750. 10.1126/sciadv.abj8750

Larsen, B., Sydnor, V. J., Keller, A. S., Yeo, B. T., & Satterthwaite, T. D. (2023). A critical period plasticity framework for the sensorimotor–association axis of cortical neurodevelopment. Trends in Neurosciences, 46(10), 847–862.

Lauterborn, J. C., Scaduto, P., Cox, C. D., Schulmann, A., Lynch, G., Gall, C. M., Keene, C. D., & Limon, A. (2021). Increased excitatory to inhibitory synaptic ratio in parietal cortex samples from individuals with Alzheimer’s disease. Nature Communications, 12(1), 2603.

Leonardi, N., & Van De Ville, D. (2015). On spurious and real fluctuations of dynamic functional connectivity during rest. Neuroimage, 104, 430–436.

Li, Z., Kovachki, N., Azizzadenesheli, K., Liu, B., Bhattacharya, K., Stuart, A., & Anandkumar, A. (2021). *Fourier Neural Operator for Parametric Partial Differential Equations* (arXiv:2010.08895). arXiv. 10.48550/arXiv.2010.08895

Moon, J.-Y., Lee, U., Blain-Moraes, S., & Mashour, G. A. (2015). General relationship of global topology, local dynamics, and directionality in large-scale brain networks. PLoS Computational Biology, 11(4), e1004225.

Müller, E. J., Munn, B. R., Redinbaugh, M. J., Lizier, J., Breakspear, M., Saalmann, Y. B., & Shine, J. M. (2023). The non-specific matrix thalamus facilitates the cortical information processing modes relevant for conscious awareness. Cell Reports, 42(8). https://www.cell.com/cell-reports/fulltext/S2211-1247(23)00855-0?uuid=uuid%3Aed1296ea-49e1-4d66-bbb9-6fb1139cf505

Nooner, K. B., Colcombe, S. J., Tobe, R. H., Mennes, M., Benedict, M. M., Moreno, A. L., Panek, L. J., Brown, S., Zavitz, S. T., & Li, Q. (2012). The NKI-Rockland sample: A model for accelerating the pace of discovery science in psychiatry. Frontiers in Neuroscience, 6, 152.

Ooi, L. Q. R., Orban, C., Zhang, S., Nichols, T. E., Tan, T. W. K., Kong, R., Marek, S., Dosenbach, N. U., Laumann, T. O., & Gordon, E. M. (2025). Longer scans boost prediction and cut costs in brain-wide association studies. Nature, 644(8077), 731–740.

Pang, J. C., Aquino, K. M., Oldehinkel, M., Robinson, P. A., Fulcher, B. D., Breakspear, M., & Fornito, A. (2023). Geometric constraints on human brain function. Nature, 618(7965), 566–574.

Petersen, R. C., Aisen, P. S., Beckett, L. A., Donohue, M. C., Gamst, A. C., Harvey, D. J., Jack, C. R., Jagust, W. J., Shaw, L. M., Toga, A. W., Trojanowski, J. Q., & Weiner, M. W. (2010). Alzheimer’s Disease Neuroimaging Initiative (ADNI): Clinical characterization. Neurology, 74(3), 201–209. 10.1212/WNL.0b013e3181cb3e25

Petitet, P., Spitz, G., Emir, U. E., Johansen-Berg, H., & O’Shea, J. (2021). Age-related decline in cortical inhibitory tone strengthens motor memory. Neuroimage, 245, 118681.

Ponce-Alvarez, A., & Deco, G. (2024). The Hopf whole-brain model and its linear approximation. Scientific Reports, 14(1), 2615.

Porges, E. C., Woods, A. J., Edden, R. A., Puts, N. A., Harris, A. D., Chen, H., Garcia, A. M., Seider, T. R., Lamb, D. G., & Williamson, J. B. (2017). Frontal gamma-aminobutyric acid concentrations are associated with cognitive performance in older adults. Biological Psychiatry: Cognitive Neuroscience and Neuroimaging, 2(1), 38–44.

Raissi, M., Perdikaris, P., & Karniadakis, G. E. (2019). Physics-informed neural networks: A deep learning framework for solving forward and inverse problems involving nonlinear partial differential equations. Journal of Computational Physics, 378, 686–707.

Richardson, B. D., Ling, L. L., Uteshev, V. V., & Caspary, D. M. (2013). Reduced GABAA receptor-mediated tonic inhibition in aged rat auditory thalamus. Journal of Neuroscience, 33(3), 1218–1227.

Saberi, A., Wischnewski, K. J., Jung, K., Lotter, L. D., Schaare, H. L., Banaschewski, T., Barker, G. J., Bokde, A. L. W., Desrivières, S., Flor, H., Grigis, A., Garavan, H., Gowland, P., Heinz, A., Brühl, R., Martinot, J.-L., Martinot, M.-L. P., Artiges, E., Nees, F., … Valk, S. L. (2025). Adolescent maturation of cortical excitation-inhibition ratio based on individualized biophysical network modeling. Science Advances, 11(23), eadr8164. 10.1126/sciadv.adr8164

Satterthwaite, T. D., Elliott, M. A., Ruparel, K., Loughead, J., Prabhakaran, K., Calkins, M. E., Hopson, R., Jackson, C., Keefe, J., & Riley, M. (2014). Neuroimaging of the Philadelphia neurodevelopmental cohort. Neuroimage, 86, 544–553.

Schaefer, A., Kong, R., Gordon, E. M., Laumann, T. O., Zuo, X.-N., Holmes, A. J., Eickhoff, S. B., & Yeo, B. T. (2018). Local-global parcellation of the human cerebral cortex from intrinsic functional connectivity MRI. Cerebral Cortex, 28(9), 3095–3114.

Sciamanna, M., Virte, M., Masoller, C., & Gavrielides, A. (2012). Hopf bifurcation to square-wave switching in mutually coupled semiconductor lasers. Physical Review E, 86(1), 016218. 10.1103/PhysRevE.86.016218

Simmonite, M., Carp, J., Foerster, B. R., Ossher, L., Petrou, M., Weissman, D. H., & Polk, T. A. (2019). Age-related declines in occipital GABA are associated with reduced fluid processing ability. Academic Radiology, 26(8), 1053–1061.

Smith, R. E., Raffelt, D., Tournier, J.-D., & Connelly, A. (2022). Quantitative streamlines tractography: Methods and inter-subject normalisation. Aperture Neuro, 2, 1–25.

Smith, R. E., Tournier, J.-D., Calamante, F., & Connelly, A. (2012). Anatomically-constrained tractography: Improved diffusion MRI streamlines tractography through effective use of anatomical information. Neuroimage, 62(3), 1924–1938.

Smith, R. E., Tournier, J.-D., Calamante, F., & Connelly, A. (2015). SIFT2: Enabling dense quantitative assessment of brain white matter connectivity using streamlines tractography. Neuroimage, 119, 338–351.

Soh, S.-E., Tint, M. T., Gluckman, P. D., Godfrey, K. M., Rifkin-Graboi, A., Chan, Y. H., Stünkel, W., Holbrook, J. D., Kwek, K., & Chong, Y.-S. (2014). Cohort profile: Growing Up in Singapore Towards healthy Outcomes (GUSTO) birth cohort study. International Journal of Epidemiology, 43(5), 1401–1409.

Stasinopoulos, D. M., & Rigby, R. A. (2008). Generalized additive models for location scale and shape (GAMLSS) in R. Journal of Statistical Software, 23, 1–46.

Sun, L., Zhao, T., Liang, X., Xia, M., Li, Q., Liao, X., Gong, G., Wang, Q., Pang, C., & Yu, Q. (2025). Human lifespan changes in the brain’s functional connectome. Nature Neuroscience, 1–11.

Sydnor, V. J., Larsen, B., Bassett, D. S., Alexander-Bloch, A., Fair, D. A., Liston, C., Mackey, A. P., Milham, M. P., Pines, A., & Roalf, D. R. (2021). Neurodevelopment of the association cortices: Patterns, mechanisms, and implications for psychopathology. Neuron, 109(18), 2820–2846.

Thomas Yeo, B. T., Krienen, F. M., Sepulcre, J., Sabuncu, M. R., Lashkari, D., Hollinshead, M., Roffman, J. L., Smoller, J. W., Zöllei, L., Polimeni, J. R., Fischl, B., Liu, H., & Buckner, R. L. (2011). The organization of the human cerebral cortex estimated by intrinsic functional connectivity. Journal of Neurophysiology, 106(3), 1125–1165. 10.1152/jn.00338.2011

Tolley, N., Rodrigues, P. L., Gramfort, A., & Jones, S. R. (2024). Methods and considerations for estimating parameters in biophysically detailed neural models with simulation based inference. PLOS Computational Biology, 20(2), e1011108.

Tournier, J. D., Calamante, F., & Connelly, A. (2010). Improved probabilistic streamlines tractography by 2nd order integration over fibre orientation distributions. Proceedings of the International Society for Magnetic Resonance in Medicine, 1670. https://archive.ismrm.org/2010/1670.html

Tournier, J. D., Smith, R., Raffelt, D., Tabbara, R., Dhollander, T., Pietsch, M., Christiaens, D., Jeurissen, B., Yeh, C.-H., & Connelly, A. (2019). MRtrix3: A fast, flexible and open software framework for medical image processing and visualisation. Neuroimage, 202, 116137.

Van Essen, D. C., Smith, S. M., Barch, D. M., Behrens, T. E., Yacoub, E., Ugurbil, K., & Consortium, W.-M. H. (2013). The WU-Minn human connectome project: An overview. Neuroimage, 80, 62–79.

Vaswani, A., Shazeer, N., Parmar, N., Uszkoreit, J., Jones, L., Gomez, A. N., Kaiser, \Lukasz, & Polosukhin, I. (2017). Attention is all you need. Advances in Neural Information Processing Systems, 30. https://proceedings.neurips.cc/paper/7181-attention-is-all

Volkow, N. D., Koob, G. F., Croyle, R. T., Bianchi, D. W., Gordon, J. A., Koroshetz, W. J., Pérez-Stable, E. J., Riley, W. T., Bloch, M. H., & Conway, K. (2018). The conception of the ABCD study: From substance use to a broad NIH collaboration. Developmental Cognitive Neuroscience, 32, 4–7.

Wang, P., Kong, R., Kong, X., Liégeois, R., Orban, C., Deco, G., van den Heuvel, M. P., & Thomas Yeo, B. T. (2019). Inversion of a large-scale circuit model reveals a cortical hierarchy in the dynamic resting human brain. Science Advances, 5(1), eaat7854. 10.1126/sciadv.aat7854

Wong, K.-F., & Wang, X.-J. (2006). A recurrent network mechanism of time integration in perceptual decisions. Journal of Neuroscience, 26(4), 1314–1328.

Yan, X., Kong, R., Xue, A., Yang, Q., Orban, C., An, L., Holmes, A. J., Qian, X., Chen, J., & Zuo, X.-N. (2023). Homotopic local-global parcellation of the human cerebral cortex from resting-state functional connectivity. NeuroImage, 273, 120010.

Zalesky, A., Fornito, A., Cocchi, L., Gollo, L. L., & Breakspear, M. (2014). Time-resolved resting-state brain networks. Proceedings of the National Academy of Sciences, 111(28), 10341–10346. 10.1073/pnas.1400181111

Zhang, S., Larsen, B., Sydnor, V. J., Zeng, T., An, L., Yan, X., Kong, R., Kong, X., Gur, R. C., Gur, R. E., Moore, T. M., Wolf, D. H., Holmes, A. J., Xie, Y., Zhou, J. H., Fortier, M. V., Tan, A. P., Gluckman, P., Chong, Y. S., … Yeo, B. T. T. (2024). In vivo whole-cortex marker of excitation-inhibition ratio indexes cortical maturation and cognitive ability in youth. Proceedings of the National Academy of Sciences, 121(23), e2318641121. 10.1073/pnas.2318641121

Zuo, X.-N., & Consortium, C. (2023). Developing Chinese Color Nest Project (devCCNP) Lite. *(No Title)*. https://cir.nii.ac.jp/crid/1883961343045027840

