## Supplementary Material for "Optimizing Biophysical Large-Scale Brain Circuit Models With Deep Neural Networks"

### Supplemental Material

This supplemental material consists of Supplemental Methods, Supplemental Tables and Supplemental Figures to complement the Methods and Results sections in the main text.

### Supplemental Methods

#### S1. Details of the FIC model

The feedback inhibition control (FIC) model (Deco et al., 2014) was derived from the mean field reduction of a spiking neuronal network model (Brunel & Wang, 2001; Wong & Wang, 2006). The detailed derivation of the FIC model can be found in a previous study (Deco et al., 2014). The differential equations governing the $j$-th cortical region are shown below:

$$\begin{aligned} I_{j}^{\left( E \right)}=W_{E}I_{0}+w_{EE}J_{NMDA}S_{j}^{\left( E \right)}+GJ_{NMDA}\sum_{k} C_{jk}S_{k}^{\left( E \right)}-w_{IE}S_{j}^{\left( I \right)}\#\left( 1 \right) \end{aligned}$$

$$\begin{aligned} I_{j}^{\left( I \right)}=W_{I}I_{0}+w_{EI}J_{NMDA}S_{j}^{\left( E \right)}-w_{II}S_{j}^{\left( I \right)} \#\left( 2 \right) \end{aligned}$$

$$\begin{aligned} r_{j}^{\left( E \right)}=\phi\left( I_{j}^{\left( E \right)} \right)=\frac{a_{E}I_{j}^{\left( E \right)}-b_{E}}{1-\exp\left( -d_{E}\left( a_{E}I_{j}^{\left( E \right)}-b_{E} \right) \right)} \#\left( 3 \right) \end{aligned}$$

$$\begin{aligned} r_{j}^{\left( I \right)}=\phi\left( I_{j}^{\left( I \right)} \right)=\frac{a_{I}I_{j}^{\left( I \right)}-b_{I}}{1-\exp\left( -d_{I}\left( a_{I}I_{j}^{\left( I \right)}-b_{I} \right) \right)} \#\left( 4 \right) \end{aligned}$$

$$\begin{aligned} \frac{dS_{j}^{\left( E \right)}}{dt}=-\frac{S_{j}^{\left( E \right)}}{\tau_{E}}+\left( 1-S_{j}^{\left( E \right)} \right)\gamma r_{j}^{\left( E \right)}+\sigma\nu_{j}\left( t \right) \#\left( 5 \right) \end{aligned}$$

$$\begin{aligned} \frac{dS_{j}^{\left( I \right)}}{dt}=-\frac{S_{j}^{\left( I \right)}}{\tau_{I}}+r_{j}^{\left( I \right)}+\sigma\nu_{j}\left( t \right) \#\left( 6 \right) \end{aligned}$$

where $S$, $r$, and $I$ denote synaptic gating variable, firing rate, and synaptic current respectively. The superscripts $E$ and $I$ represent the excitatory and inhibitory neuronal populations respectively.

The input current $I_{j}^{(E)}$ of the excitatory population of the $j$-th cortical ROI is the sum of four inputs (Equation 1). The first input is the external input current $W_{E}I_{0}$, which might include subcortical delays. The second input is the intra-regional excitatory-to-excitatory current governed by the excitatory-to-excitatory recurrent connection strength $w_{EE}$ scaled by the synaptic coupling constant $J_{NMDA}$. The third input is the inter-regional input, which is controlled by the SC matrix ($C_{jk}$ is the connectivity between regions $j$ and $k$) and scaled by the global constant $G$. The fourth input is the intra-regional negative feedback from the inhibitory population governed by the inhibitory-to-excitatory connection strength $w_{IE}$.

The input current $I_{j}^{(I)}$ of the inhibitory population of the $j$-th cortical ROI is the sum of three inputs (Equation 2). The first input is the external input current $W_{I}I_{0}$. The second input is the intra-regional positive feedback from the excitatory population governed by the excitatory-to-inhibitory connection strength *w_EI_* scaled by the synaptic coupling constant $J_{NMDA}$. The third input is the intra-regional inhibitory-to-inhibitory current governed by the inhibitory-to-inhibitory recurrent connection strength $w_{II}$.

The excitatory input current $I_{j}^{(E)}$ and inhibitory input current $I_{j}^{(I)}$ are transformed into firing rates via the input-output functions specified in Equations 3 and 4. Following previous studies (Deco et al., 2014), the parameters of the input-output function were set to be excitatory gain $a_{E}=310$n/C, inhibitory gain $a_{I}=615$n/C, $b_{E}=125$Hz, $b_{I}=177$Hz, $d_{E}=0.16$s and $d_{I}=0.087$s. Finally, the rate of change of the synaptic gating variables $S_{j}^{(E)}$ and $S_{j}^{(I)}$ are computed via equations 5 and 6. Following previous studies (Deco et al., 2014), the kinetic parameters for synaptic activities $\tau_{E}$, $\tau_{I}$and $\gamma$ were set to 100ms, 10ms and 0.641 respectively. $\nu_{j}\left( t \right)$ corresponds to uncorrelated standard Gaussian noise with the noise amplitude being controlled by $\sigma$.

Following the original study (Deco et al., 2014), $w_{II}$, $W_{E}$, $W_{I}$, $I_{0}$ and $J_{NMDA}$ were set to 1, 1, 0.7 0.382nA and 0.15nA respectively in the current study. The inhibitory-to-excitatory connection strength $w_{IE}$ was computed analytically to ensure that the excitatory firing rate is maintained to be around 3Hz (Brunel & Wang, 2001; Deco et al., 2014).

The local circuit parameters, including $w_{EE}$ (excitatory-to-excitatory recurrent connection strength), $w_{EI}$ (excitatory-to-inhibitory connection strength), $\sigma$ (noise amplitude) and $G$ (global SC scaling constant) are unspecified and will be inferred by fitting to empirical fMRI data. Given a fixed set of model parameters, equations 1 to 6 were used to simulate the time courses of excitatory and inhibitory synaptic gating variables ($S_{j}^{(E)}$ and $S_{j}^{(I)}$) of each ROI with a fixed local circuit parameter set. The regional E/I ratio was defined as the average temporal ratio between $S_{j}^{(E)}$ and $S_{j}^{(I)}$. The mean cortical E/I ratio was then derived from averaging regional E/I ratios across all cortical ROIs. Additionally, the simulated excitatory synaptic gating variables ($S_{j}^{(E)}$) were fed into the Balloon-Windkessel hemodynamic model to generate fMRI BOLD signals (Stephan et al., 2007; Deco et al., 2014). The simulated fMRI BOLD signals were utilized to generate simulated static FC and FCD.

#### S2. Optimizing the FIC model with CMA-ES in the HCP dataset

As mentioned in the previous section, we seek to optimize $w_{EE}$ (excitatory-to-excitatory recurrent connection strength), $w_{EI}$ (excitatory-to-inhibitory connection strength), $\sigma$ (noise amplitude) and $G$ (global SC scaling constant). We assumed that $w_{EE}$, $w_{EI}$ and $\sigma$ were spatially heterogeneous. Assuming a 68-region Desikan-Killiany parcellation (Desikan et al., 2006), this resulted in a different $w_{EE}$, $w_{EI}$ and $\sigma$ for each brain region, yielding 68 × 3 + 1 = 205 parameters to be estimated.

To further constrain the parameter space (Zhang et al., 2024), w_EE_, w_EI_, and $\sigma$ were parameterized as linear combinations of the principal resting-state functional connectivity gradient (Margulies et al., 2016) and the T1w/T2w myelin map (Glasser & Van Essen, 2011).

$$\begin{aligned} w_{EE,j}=a+b\times{T1w/T2w \mathrm{ratio}}_{j}+c\times{FC gradient}_{j} \#\left( 7 \right) \end{aligned}$$

$$\begin{aligned} w_{EI,j} =d+e\times{T1w/T2w \mathrm{ratio}}_{j}+f\times{FC gradient}_{j} \#\left( 8 \right) \end{aligned}$$

$$\begin{aligned} \sigma_{j}=g+h\times{T1w/T2w \mathrm{ratio}}_{j}+i\times{FC gradient}_{j}\#\left( 9 \right) \end{aligned}$$

where *j* denotes the *j*-th region. By employing this parameterization strategy, the number of “free” numbers was reduced to 3 × 3 + 1 = 10 parameters (including $G$).

The 10 parameters were optimized using CMA-ES by minimizing dissimilarity of the simulated FC and FC to empirical FC and FCD. The agreement between empirical and simulated FC matrices was defined as the Pearson’s correlation (*r*) between the z-transformed upper triangular entries of the two matrices. Larger *r* indicates more similar static FC. However, Pearson’s correlation does not account for scale difference, so we also computed the absolute difference (*d*) between the means of the empirical and simulated FC matrices (Demirtaş et al., 2019). A smaller d indicates more similar static FC.

We note that there is no temporal correspondence between simulated and empirical FCD matrices, so we cannot simply use the Euclidean distance to measure dissimilarity. Instead, the dissimilarity between simulated and empirical FCD matrices was quantified by using the Kolmogorov-Smirnov (*KS*) distance. Here, the KS distance was defined as the maximum distance between the cumulative distribution functions (CDFs) constructed by collapsing the upper triangular entries of simulated and empirical FCD matrices (Hansen et al., 2015; Kong et al., 2021). Hence, a small *KS* distance indicated 2 similar CDFs, therefore 2 similar FCD matrices. Because the *KS* distance was computed by collapsing the upper triangular entries of the FCD matrices, no temporal correspondence was assumed.

The overall FC+FCD cost was defined as (1 – *r*) + *d* + *KS*. A smaller cost indicates better agreement between simulated and empirical fMRI. Throughout the study, we note that FC+FCD cost was computed at the group level. For more details about how group-level FC, FCD and SC are computed, see Supplementary Methods S3.

In the case of evaluating DELSSOME CMA-ES in the HCP dataset (Fig. 3), recall that the HCP test participants (N = 169) were further divided into the FIC model inversion training (N = 57), validation (N = 56) and test (N = 56) sets (Fig. 3a). Group-average FC and FCD were computed separately in the training, validation and test sets. Group-average SC, FC gradient and T1w/T2w ratio was computed only in the HCP training set (N = 680) and were used throughout the analysis.

Euler CMA-ES was run on the FIC model inversion training set for 100 epochs. FC+FCD cost was computed using FIC model inversion training set FC and FCD. The best candidate parameter set from each epoch was selected, yielding 100 candidate parameter sets. The 100 candidate parameter sets were then evaluated in the FIC model inversion validation set using the FIC model inversion validation set FC and FCD. Finally, the top parameter set from the validation set was evaluated in the FIC model inversion test set using the FIC model inversion test set FC and FCD.

When computing FC+FCD cost using Euler integration, the FIC model was simulated three times, and the average simulated FC and FCD were used to compute the FC+FCD cost. Euler integration in the FIC model inversion training and validation sets was performed with timestep of 6ms, while Euler integration in the FIC model inversion test sets was performed with timestep of 0.5ms.

DELSSOME CMA-ES utilized the same procedure as Euler CMA-ES, except that we used DELSSOME instead of Euler integration in the FIC model inversion training set. However, we still used Euler integration in the FIC model inversion validation and test sets.

#### S3. Group-level FC, FCD and SC

Across different sections of this study, group-average FC, FCD and SC were computed based on different groups of participants. Our procedure followed that of our previous studies (Kong et al., 2021; Zhang et al., 2024). More specifically, to compute group-level FC across a group of participants, we applied Fisher-r-to-z transformation to the FC matrix of each run of each participant. The FC matrices were then averaged across runs within each participant, followed by averaging across all participants and then applying the inverse Fisher-r-to-z transformation.

In the case of FCD, recall that the FCD each participant was a W × W matrix, where W is the number of sliding windows (W = 1118 in HCP and 109 in PNC). Unlike static FC, FCD matrices could not be directly averaged across participants due to the lack of temporal correspondence during resting-state. As discussed in the previous section, we used the Kolmogorov–Smirnov (KS) distance to measure the distance between the two FCD matrices, consistent with previous studies (Zhang et al., 2024). The KS distance between two FCD matrices was defined as the maximum distance between the cumulative distribution functions (CDFs) obtained by collapsing the upper triangular entries of simulated and empirical FCD matrices, so no temporal correspondence was assumed. Therefore, given the FCD matrices of a group of participants, instead of averaging the FCD matrices, we averaged the CDF of the FCD matrices.

To generate a group-level SC matrix, a thresholding procedure was first applied to remove false positives (de Reus & van den Heuvel, 2013). More specifically, if <50% of participants had a non-zero value in a particular entry in the SC matrix, then the entry is set to zero in all individual-level SC matrices. Then averaging across participants with non-zero streamlines, log-transforming the averaged values, and setting the main diagonal entries to zero were performed. Group-level SCs were computed separately for a particular group of participants, with the maximum value normalized to 0.02.

#### S4. Optimizing the FIC model with CMA-ES in the PNC dataset

To evaluate whether DELSSOME CMA-ES generalizes to a new dataset, the DELSSOME models trained from the HCP dataset were applied directly to the Philadelphia Neurodevelopment Cohort (PNC) dataset (Figs. 4 and S5). Because there was no diffusion data and T1w/T2w ratio in the PNC dataset, group-average SC, FC gradient and T1w/T2w ratio from the HCP training set (N = 680) were used.

In the case of associating E/I ratio with age, recall that PNC participants were sorted according to age and divided into 29 age groups (with 30 or 31 participants in each group). Within each age group, 15 participants were randomly selected as the validation set, while the remaining participants were assigned to the training set. Group-average FC and FCD were computed separately in the training and validation sets.

For each age group, DELSSOME CMA-ES was applied to the training set for 50 epochs with FC+FCD cost computed using the training set FC and FCD. The procedure was repeated five times with different random initializations, resulting in 250 candidate parameter sets. The 250 parameter sets were evaluated in the validation set with Euler integration using the validation set FC and FCD. The best candidate parameter set was then used to generate an excitation/inhibition (E/I) ratio map using Euler integration.

When computing FC+FCD cost using Euler integration, the FIC model was simulated three times, and the average simulated FC and FCD were used to compute the FC+FCD cost. When computing E/I ratio, the FIC model was also simulated three times, and the E/I ratio estimates were averaged across the three simulations. Euler integration in the training set and validation set was performed with timestep of 6ms, while Euler integration in the computation of the final E/I ratio was performed with timestep of 0.5ms.

Furthermore, consistent with our previous study (Zhang et al., 2024), during CMA-ES, we additionally imposed the constraints that the spatial correlation between w_EI_ and T1w/T2w ratio should be negative, while the spatial correlation between w_EI_ and FC gradient should be positive. Euler CMA-ES followed the same procedure as DELSSOME CMA-ES, except that Euler integration was used throughout.

In the case of associating E/I ratio with cognitive performance, the same DELSSOME CMA-ES and Euler CMA-ES procedures were used as the age analysis, except that instead of participants divided into different age groups, we now have participants divided into 14 high-performance and 14 low-performance groups. Each high-performance group was age-matched to a corresponding low-performance group. Each low-performance or high-performance group comprised 31 or 32 participants.

For each group, 15 participants were randomly assigned to the validation set, while the remaining participants were assigned to the training set. For each group, 250 candidate parameter sets were generated from the training set using DELSSOME CMA-ES. The 250 parameter sets were evaluated in the validation set with Euler integration. The best candidate parameter set was then used to generate an excitation/inhibition (E/I) ratio map using Euler integration. We then compared the E/I ratio between the high and low performance groups. The same procedure was repeated with Euler CMA-ES.

#### S5. DELSSOME FC+FCD cost predictor

As mentioned in the main text, the inputs to the DELSSOME FC+FCD cost predictor were the 10 FIC parameters and the structural connectivity (SC) matrix, as well as the empirical FC and FCD matrices (Fig. 8b). The model was trained on the HCP training samples (N = 640,000), and hyperparameters were empirically determined on the HCP validation samples (N = 140,000). We emphasized that the HCP test samples (N = 130,000) were not used to train or tune the hyperparameters.

For the 68-region Desikan–Killiany parcellation, equations 7 to 9 (Supplementary Methods S2)

expanded the 9 FIC parameters into 3 local circuit parameters ($w_{EE},w_{EI},\sigma$) for each region, while the global coupling parameter G scales the 68×68 SC matrix. For each Desikan-Killiany ROI, the region’s three parameters were mapped to a d/2-dimensional representation via a single-layer MLP, followed by LayerNorm. For each ROI, the region’s SC profile (i.e., corresponding row of the SC matrix) was scaled by the global coupling parameter G, and mapped to a d/2-dimensional representation through another single-layer MLP, followed by LayerNorm. The two d/2-dimensional representations were concatenated to form a d-dimensional token embedding. Since there were 68 Desikan-Killiany ROIs, this process yielded a sequence of 68 d-dimensional tokens. In addition, a [CLS] token of dimensionality d was prepended to the sequence of 68 tokens, and the entire sequence of 69 tokens was passed to a transformer encoder (Devlin et al., 2019). The hidden state corresponding to the [CLS] token was extracted as the FIC model embedding.

The transformer encoder consisted of 4 layers, each with 8 attention heads. The model dimension was d=64, and the hidden dimension of the feed-forward layers is 4d. GeLU is used as the activation function throughout.

Since the FC cost involved comparing the simulated FC and empirical FC, so the empirical FC matrix was another input to the DELSSOME FC+FCD cost predictor. The upper triangle entries of the 68 × 68 FC matrix (2278 entries) were vectorized and fed into a single-layer MLP resulting in d-dimensional vector, followed by LayerNorm.

The resulting FC embedding vector (of length d) and the FIC model embedding vector (of length d) were added together and passed through a single-layer MLP (with sigmoid activation) to predict the FC costs *1-r* and *d* respectively.

Similarly, the FCD cost involved comparing the simulated FCD and empirical FCD, so the empirical FCD was another input to the DELSSOME FC+FCD cost predictor. The empirical FCD was represented as a probability distribution function (PDF) of FCD values (which ranged from -1 to 1). The range was discretized into 10,000 equal sized bins, so the FCD was essentially a vector of length 10,000 that summed to one. The FCD vector was fed into a single-layer MLP, resulting in d-dimensional vector. LayerNorm was applied before and after projection.

The resulting FCD embedding vector (of length d) and the FIC model embedding vector (of length d) were added together and passed through a single-layer MLP (with sigmoid activation) to predict the FCD cost *KS*.

For the 68-region Desikan–Killiany parcellation, all biophysical models (FIC, MFM, Hopf) were trained with d=64, sharing the same DELSSOME architecture.

**Training details.** For each of the three terms in the FC+FCD cost (1−r, d, KS), the squared error between the predicted and empirical values was computed. The overall training loss was the mean squared error (MSE) across the three terms. We used the Adam optimizer (Kingma, 2014) with initial learning rates of 1e-4, 1e-3, 1e-2 and 1e-1, together with an exponential learning rate scheduler (γ=0.98). Models were trained for 30 epochs with a batch size of 256. The final model was selected from the epoch with the lowest validation MSE.

**100-region Yan parcellation.** For the 100-region Yan parcellation, we used d=128 instead of d=64. Since there were 100 regions, the number of tokens increased to 101 (including [CLS]). The SC profile embeddings and FC embeddings were updated to reflect the new matrix dimensions (upper triangle of 100×100 instead of 68×68). All other architectural and training procedures remained unchanged.

#### S6. Details of individual-level DELSSOME for the FIC model

We used a total of 14 datasets comprising more than 12,005 participants to construct a normative trajectory of the excitation–inhibition (E/I) ratio across the lifespan.

We began by evaluating DELSSOME individual-level cost predictor using 12 of the 14 datasets. For each dataset, we randomly selected 40 training, 20 validation, and 20 test participants. In the case of the HCP-YA dataset, care was taken to only select participants from the HCP-YA test set (Fig. 3a), which was not previously used to train the group-level DELSSOME model.

For each participant in the training and validation sets, we ran Euler CMA-ES for 100 epochs using a time step of 5 ms. Each CMA-ES epoch produced 100 children processes, yielding 10,000 candidate FIC parameter sets per participant, each paired with the corresponding ground-truth FC+FCD cost. Consequently, each dataset contributed 400,000 training and 200,000 validation samples. Across the 12 datasets used for training and validation, this resulted in a total of 4,800,000 training samples and 2,400,000 validation samples. The training set was used to train the model, while the validation set was used to tune the hyperparameters.

Since individual-level FC and FCD-PDF were significantly noisier than group-level FC and FCD-PDF, we did not use the same hyperparameters as group-level DELSSOME. Instead, the individual-level DELSSOME cost predictor was trained using hyperparameters selected via 50 trials of the Optuna python package (Akiba et al., 2019), requiring approximately 65 hours of a single RTX3090 GPU. The final architecture and optimization settings were as follows: transformer hidden dimension 128 with 16 attention heads and 5 transformer layers; exponential learning rate schedule with initial learning rate 2.2×10⁻⁴ and decay factor γ = 0.97; all MLPs in the model were implemented as 2-layer MLPs with 146 and 58 units respectively. The loss function was the mean squared error (MSE) between predicted and ground-truth values of 1–*r*, *d*, and *KS*.

For each of the 20 test participants per dataset, we compared Euler CMA-ES with DELSSOME CMA-ES. Each approach generated 10,000 candidate parameter sets per participant. The top 100 parameter sets from each method were then re-evaluated using Euler integration with a 5 ms step size, and the single best-performing parameter set was further evaluated using a 0.5 ms step size for more accurate cost estimation. Statistical significance was assessed using a two-sided paired-sample t-test based on the 20 test participants (d.o.f. = 19). An overall two-sided paired-sample t-test was also computed based on the 12 datasets (d.o.f. = 11).

To compute the normative lifespan trajectory of the E/I ratio, we applied DELSSOME CMA-ES to every participant across all 14 datasets. For each participant, DELSSOME CMA-ES was run for 100 epochs, generating 10,000 candidate parameter sets. The best 100 candidates were subsequently validated using Euler integration with a 5 ms step size, and the best single parameter set was re-evaluated using a 0.5 ms step size. The resulting E/I ratio served as that participant’s estimate used in constructing the normative curve.

After estimating the E/I ratio for each participant, we fitted the E/I ratios using the generalized additive model for location, scale, and shape (GAMLSS; Stasinopoulos & Rigby, 2008), following established procedures in large multi-site studies (Bethlehem et al., 2022; Sun et al., 2025). GAMLSS provides a flexible framework for modeling complex outcome distributions by decomposing a distribution into up to four parameters, which are location ($\mu$), scale ($\sigma$), skewness ($\nu$), and kurtosis ($\tau$). Each parameter is linked to its own regression equation. In this formulation, outcome *Y* follows a distribution $F(\mu,\sigma,\nu,\tau)$, and each parameter is expressed as a combination of fixed effects, random effects, and smooth nonlinear functions. For example, location $\mu$ is modeled as ${g_{\mu}\left( \mu\right)=X}_{\mu}\beta_{\mu}+Z_{\mu}\gamma_{\mu}+\sum_{i} s_{\mu,i}(x_{i})$, where $X_{\mu}\beta_{\mu}$ represents typical covariates and corresponding coefficients, $Z_{\mu}\gamma_{\mu}$ represents random effects and corresponding coefficients, and $s_{\mu,i}(\mu)$ denotes smoothing functions applied to i-th covariate. The same equations apply to $\sigma,\nu$ and $\tau$. Random effects in different distributional components are assumed to be independent.

Once the model is fitted, the centile of an observation is obtained using the cumulative distribution function (CDF) of the specified outcome distribution. For an observed value $y$, the centile is given by $q=F(y\mid\mu(X,Z),\sigma(X,Z),\nu(X,Z),\tau(X,Z))$. These centiles depend on the study random effects and therefore reflect site-specific variation. To obtain harmonized, reference-normalized values, we identify the value on the reference distribution. It is defined as the same model with random effects removed ($Z=0$), which corresponds to the same centile, yielding $w=F^{-1}(q\mid\mu(X),\sigma(X),\nu(X),\tau(X))$. This procedure maps each participant’s value onto the centile structure of the reference population, effectively harmonizing measurements across sites. These harmonized values were used for visualization (Fig. 7a).

We modeled E/I ratios using the GAMLSS (Generalized Additive Models for Location, Scale, and Shape) framework, which allows distributional parameters to vary as functions of covariates. Specifically, age was modeled as a smooth nonlinear effect using cubic B-spline basis functions, with 4 degrees of freedom for the location parameter (μ) and 3 degrees of freedom for the scale parameter (σ). Sex was included as a fixed effect, and scanner site was modeled as a random effect to account for inter-site variability. No interaction terms were included. The skewness (ν) and kurtosis (τ) parameters were assumed to be constant (i.e., not modeled as functions of covariates). To determine the optimal distributional family and spline complexity, we conducted a hyperparameter search across multiple continuous distributions (including SHASH, Johnson’s SU [JSU], generalized gamma, and normal distributions). Model selection was based on the Bayesian Information Criterion (BIC). The best-fitting model, adopted for all final analyses, used the SHASHo2 distribution with a cubic B-spline age effect on μ with 4 degrees of freedom.

#### S7. Details of mean field model (MFM)

The MFM was derived from a mean-field reduction of a detailed spiking neuronal network model (Deco et al., 2013). For each cortical region of interest (ROI), neural activity is governed by the following nonlinear stochastic differential equations:

$$\begin{aligned} \frac{dS_{i}}{dt}=-\frac{S_{i}}{\tau_{s}}+r\left( 1-S_{i} \right)H\left( x_{i} \right)+\sigma v_{i}\left( t \right)\#\left( 10 \right) \end{aligned}$$

$$\begin{aligned} H\left( x_{i} \right)=\frac{ax_{i}-b}{1-\exp\left( -d\left( ax_{i}-b \right) \right)} \#\left( 11 \right) \end{aligned}$$

$$\begin{aligned} x_{i}=wJS_{i}+GJ\sum_{j} C_{ij}S_{j}+I \#\left( 12 \right) \end{aligned}$$

where $S_{i},H(x_{i})$ and $x_{i}$ denote the average synaptic gating variable, population firing rate, and total input current of the i-th ROI, respectively. The total input current $x_{i}$ reflects the sum of three contributions: (1) Intra-regional input, controlled by the recurrent connection strength *w*. (2) Inter-regional input, determined by the structural connectivity (SC) matrix, where $C_{ij}$ represents the SC between regions i and j, scaled globally by a coupling factor G. (3) External input, a constant input current *I*, potentially reflecting contributions from subcortical relays.

Following previous studies (Deco et al., 2013; Wang et al., 2019), the synaptic coupling was fixed at J=0.2609 nA. The parameters of the input–output function $H\left( x_{i} \right)$ were set to a=270 (n/C), b=108 Hz, and d=0.154 s. Kinetic parameters for synaptic activity were fixed at r=0.641 and $\tau_{s}$=0.1s. The noise term $v_{i}\left( t \right)$ represents uncorrelated Gaussian white noise with amplitude σ.

The simulated neural activities $S_{i}$ were passed to the Balloon–Windkessel hemodynamic model to generate simulated BOLD signals for each ROI (Stephan et al., 2007; Deco et al., 2013).

As in equations 7–9, the parameters *w*, *I*, and σ were further parameterized as linear combinations of the principal resting-state functional connectivity gradient and the T1w/T2w myelin map:

$$\begin{aligned} w_{j}=a+b\times{T1w/T2w \mathrm{ratio}}_{j}+c\times{FC gradient}_{j} \#\left( 13 \right) \end{aligned}$$

$$\begin{aligned} I_{j} =d+e\times{T1w/T2w \mathrm{ratio}}_{j}+f\times{FC gradient}_{j} \#\left( 14 \right) \end{aligned}$$

$$\begin{aligned} \sigma_{j}=g+h\times{T1w/T2w \mathrm{ratio}}_{j}+i\times{FC gradient}_{j} \#\left( 15 \right) \end{aligned}$$

where *j* denotes the *j*-th region. This parameterization reduced the number of free parameters to 3 × 3 + 1 = 10, including the global scaling parameter *G*.

#### S8. Details of the Hopf model

Here, we followed the Hopf model from Ponce-Alvarez and Deco (Ponce-Alvarez & Deco, 2024). The dynamics of an isolated node are governed by the normal form of a supercritical Hopf bifurcation. Extending this to a network of *N* coupled nodes, the dynamics are described by:

$$\frac{dz_{j}}{dt}=\left( a_{j}+i\omega_{j} \right)z_{j}-\left| z_{j} \right|^{2}z_{j}+g\sum_{k=1}^{N} C_{jk}\left( z_{k}-z_{j} \right)+\eta_{j}$$

where $z_{j}=x_{j}+iy_{j},\eta_{j}=\eta_{x_{j}}+i\eta_{y_{j}}$, $\left| z_{j} \right|^{2}=x_{j}^{2}+y_{j}^{2}$. The intrinsic angular frequency is denoted by $\omega\left( rad\cdot s^{-1} \right)$, and *a* is the bifurcation parameter (s^-1^). The parameter *g* (s^-1^) scales the contribution of the SC matrix *C*. The additive white noise term $\eta$ satisfies $\left\langle\eta(t) \right\rangle=0$ and $\left\langle\eta\left( t \right)\eta\left( t^{'} \right) \right\rangle=\sigma^{2}\delta\left( t-t^{'} \right)$, where $\sigma$ is the noise amplitude (s^-1/2^) and $\left\langle\cdot\right\rangle$ denotes averaging across stochastic realizations.

Similar to equations 7 to 9, the parameters *a*, $\omega$, and σ were further parameterized as linear combinations of the principal resting-state functional connectivity gradient and the T1w/T2w myelin map:

$$\begin{aligned} a_{j}=a+b\times{T1w/T2w \mathrm{ratio}}_{j}+c\times{FC gradient}_{j} \#\left( 16 \right) \end{aligned}$$

$$\begin{aligned} \omega_{j} =d+e\times{T1w/T2w \mathrm{ratio}}_{j}+f\times{FC gradient}_{j} \#\left( 17 \right) \end{aligned}$$

$$\begin{aligned} \sigma_{j}=g+h\times{T1w/T2w \mathrm{ratio}}_{j}+i\times{FC gradient}_{j} \#\left( 18 \right) \end{aligned}$$

where *j* denotes the *j*-th region. This parameterization reduced the number of free parameters to 3 × 3 + 1 = 10, including the global scaling parameter *G*.

Consistent with previous studies (Ponce-Alvarez and Deco, 2024), the simulated timeseries $z_{j}$ were directly used to compute FC and FCD, after downsampling to match the TR in the HCP dataset.

#### S9. Benchmarking of simulation-based inference

Following previous studies in neuroscience (Gonçalves et al., 2020; Tolley et al., 2024), we employed simulation-based inference (SBI) using Sequential Neural Posterior Estimation (SNPE) coupled with a Masked Autoregressive Flow (MAF) density estimator. This approach enables flexible, likelihood-free inference for complex biophysical models, where analytical likelihood functions are intractable but simulations can be readily generated.

**Overview of the SNPE framework**. SNPE (Greenberg et al., 2019) learns an explicit neural approximation to the posterior distribution $p\left( \theta| x \right)$ by training a conditional density estimator on pairs of simulated parameters $\theta$ and corresponding model outputs *x*. At each training iteration, parameters are sampled from a proposal distribution and fed into the generative model to produce simulated data. The neural network then minimizes a loss function that encourages accurate posterior reconstruction given the simulations. We adopt the implementation provided in the *sbi* Python package (Boelts et al., 2025), using SNPE-C (Greenberg et al., 2019), which combines sequential training and a likelihood-ratio–based objective for improved stability.

**Density estimator: Masked Autoregressive Flow (MAF)**. The posterior density $q_{\phi}\left( \theta| x \right)$ was modeled using a Masked Autoregressive Flow (MAF; Papamakarios et al., 2017), a flexible normalizing flow architecture that transforms a simple base distribution (e.g., Gaussian) into a complex multimodal target via a sequence of invertible transformations. Each transformation layer employs an autoregressive structure to ensure tractable computation of the Jacobian determinant, allowing efficient optimization of the log-likelihood. In our configuration, the MAF was instantiated with the following architecture: 64 units per hidden layer, consistent with DELSSOME cost predictor, 8 sequential flow layers, the embedding network consisting of a two-tower embedder combining functional connectivity (FC) and functional connectivity dynamics (FCD) features. We used standard multivariate normal as the base distribution. This architecture provided sufficient representational power while maintaining numerical stability and moderate computational cost following previous studies (Gonçalves et al., 2020; Tolley et al., 2024).

**Training procedure and hyperparameters**. The inference process was initialized with a uniform prior over a predefined parameter range. For each group of simulations *k*, parameter sets $\theta_{k}\in\mathbb{R}^{100\times10}$ and corresponding summary features $x_{k}$ were generated. Each $x_{k}$ concatenated group-level FC and FCD-PDF features:

$$x_{k}=\left[ FC_{k},FCD PDF_{k} \right]$$

Simulated parameter–data pairs were appended to the SNPE object with invalid simulations excluded. The model was then trained with the following settings: 20% of simulations held out for validation. The combined loss was used for balancing log-likelihood and calibration objectives. We used the Adam optimizer with initial learning rates of 1e-4, 1e-3, 1e-2 and 1e-1. Models were trained for 30 epochs with a batch size of 256. The final model was selected from the epoch with the lowest validation loss.

For benchmarking with DELSSOME CMA-ES, we sampled 10,000 times from the resulting posterior. The 100 samples with highest log probability were fed into validation set and the one with best validation cost was used for test. Euler integration was utilized for both validation and test sets. The test cost was reported. The benchmarking procedure were repeated 50 times, consistent with DELSSOME CMA-ES.

### Supplemental Tables

**Table S1.** Summary of sites, scanner, sample size, TR and scan duration.

| Dataset | Site | Scanner model | N | TR (s) | Scan duration (s) |
| --- | --- | --- | --- | --- | --- |
| ABCD | ABCD_CHLA | Philips Achieva dStream | 81 | 0.8 | 300 |
| ABCD | ABCD_CUB | Siemens Prisma fit | 280 | 0.8 | 300 |
| ABCD | ABCD_FIU | Siemens Prisma | 233 | 0.8 | 300 |
| ABCD | ABCD_LIBR | GE MR750 DV25 | 209 | 0.8 | 300 |
| ABCD | ABCD_LIBR | GE MR750 DV26 | 188 | 0.8 | 300 |
| ABCD | ABCD_MSSM | GE MR750 DV25 | 5 | 0.8 | 300 |
| ABCD | ABCD_MUSC | Siemens Prisma fit | 147 | 0.8 | 300 |
| ABCD | ABCD_OHSU | Siemens Prisma fit | 209 | 0.8 | 300 |
| ABCD | ABCD_ROC | Siemens Prisma fit | 132 | 0.8 | 300 |
| ABCD | ABCD_SRI | GE MR750 DV25 | 31 | 0.8 | 300 |
| ABCD | ABCD_SRI | GE MR750 DV26 | 68 | 0.8 | 300 |
| ABCD | ABCD_UCLA | Siemens Prisma fit | 119 | 0.8 | 300 |
| ABCD | ABCD_UCSD | GE MR750 DV25 | 112 | 0.8 | 300 |
| ABCD | ABCD_UCSD | GE MR750 DV26 | 195 | 0.8 | 300 |
| ABCD | ABCD_UFL | Siemens Prisma | 178 | 0.8 | 300 |
| ABCD | ABCD_UMB | Siemens Prisma fit | 248 | 0.8 | 300 |
| ABCD | ABCD_UMICH | GE MR750 DV25 | 135 | 0.8 | 300 |
| ABCD | ABCD_UMICH | GE MR750 DV26 | 121 | 0.8 | 300 |
| ABCD | ABCD_UMN | Siemens Prisma | 244 | 0.8 | 300 |
| ABCD | ABCD_UMN | Siemens Prisma fit | 48 | 0.8 | 300 |
| ABCD | ABCD_UPMC | Siemens Prisma fit | 105 | 0.8 | 300 |
| ABCD | ABCD_UTAH | Siemens Prisma | 540 | 0.8 | 300 |
| ABCD | ABCD_UVM | Philips Achieva dStream | 181 | 0.8 | 300 |
| ABCD | ABCD_UWM | GE MR750 DV25 | 34 | 0.8 | 300 |
| ABCD | ABCD_UWM | GE MR750 DV26 | 118 | 0.8 | 300 |
| ABCD | ABCD_VCU | Philips Ingenia | 106 | 0.8 | 300 |
| ABCD | ABCD_WUSTL | Siemens Prisma | 249 | 0.8 | 300 |
| ABCD | ABCD_WUSTL | Siemens Prisma fit | 60 | 0.8 | 300 |
| ABCD | ABCD_YALE | Siemens Prisma | 48 | 0.8 | 300 |
| ABCD | ABCD_YALE | Siemens Prisma fit | 185 | 0.8 | 300 |
| ADNI | ADNI1 | Philips Intera | 15 | 3.001* | 420.1* |
| ADNI | ADNI1 | Siemens Prisma fit | 9 | 3.000* | 591.0* |
| ADNI | ADNI10 | Philips GEMINI | 4 | 3 | 420 |
| ADNI | ADNI10 | Philips Ingenuity | 2 | 3 | 420 |
| ADNI | ADNI11 | Siemens Verio | 9 | 3 | 591 |
| ADNI | ADNI12 | GE Signa HDxt | 2 | 2.925 | 468 |
| ADNI | ADNI13 | Philips Achieva | 5 | 3.000* | 420.0* |
| ADNI | ADNI13 | Philips Achieva dStream | 1 | 3 | 591 |
| ADNI | ADNI14 | Philips Achieva | 2 | 3 | 591 |
| ADNI | ADNI15 | Siemens Prisma fit | 10 | 3 | 591 |
| ADNI | ADNI16 | GE DISCOVERY MR750 | 17 | 3.000* | 600.0* |
| ADNI | ADNI17 | Siemens Prisma fit | 3 | 0.607 | 592.4 |
| ADNI | ADNI18 | Siemens Prisma fit | 7 | 0.607* | 592.4* |
| ADNI | ADNI19 | Siemens Prisma fit | 8 | 3.000* | 591.0* |
| ADNI | ADNI2 | Siemens Prisma | 18 | 3 | 591 |
| ADNI | ADNI20 | GE DISCOVERY MR750 | 18 | 3 | 600 |
| ADNI | ADNI21 | GE DISCOVERY MR750w | 9 | 3 | 600 |
| ADNI | ADNI21 | Philips Achieva | 7 | 3 | 420 |
| ADNI | ADNI22 | Philips Achieva | 1 | 3 | 591 |
| ADNI | ADNI23 | Siemens Prisma fit | 3 | 3 | 591 |
| ADNI | ADNI25 | Siemens Prisma fit | 16 | 3 | 591 |
| ADNI | ADNI26 | Siemens Skyra | 3 | 0.788 | 591 |
| ADNI | ADNI27 | Siemens Prisma | 23 | 0.607* | 592.4* |
| ADNI | ADNI28 | Siemens Prisma fit | 24 | 3 | 591 |
| ADNI | ADNI3 | GE DISCOVERY MR750 | 2 | 3 | 600 |
| ADNI | ADNI30 | Philips Intera | 4 | 3.000* | 420.0* |
| ADNI | ADNI30 | Siemens Prisma | 1 | 0.607 | 592.4 |
| ADNI | ADNI33 | Siemens Prisma fit | 18 | 0.607* | 592.4* |
| ADNI | ADNI38 | Siemens Prisma | 10 | 3.000* | 591.0* |
| ADNI | ADNI39 | GE SIGNA Premier | 4 | 3 | 600 |
| ADNI | ADNI4 | Philips Ingenia | 17 | 3.000* | 591.0* |
| ADNI | ADNI40 | GE DISCOVERY MR750 | 9 | 3 | 600 |
| ADNI | ADNI40 | Philips Achieva | 9 | 3.000* | 420.0* |
| ADNI | ADNI41 | Philips Achieva | 5 | 3 | 591 |
| ADNI | ADNI43 | Siemens Skyra fit | 3 | 3 | 591 |
| ADNI | ADNI43 | Siemens Verio | 7 | 3 | 591 |
| ADNI | ADNI44 | Siemens TrioTim | 4 | 3 | 591 |
| ADNI | ADNI45 | Philips Ingenia | 2 | 3 | 591 |
| ADNI | ADNI46 | GE DISCOVERY MR750 | 2 | 3 | 600 |
| ADNI | ADNI47 | GE DISCOVERY MR750 | 21 | 3 | 600 |
| ADNI | ADNI49 | GE DISCOVERY MR750 | 12 | 3 | 600 |
| ADNI | ADNI49 | Philips Ingenia | 4 | 3 | 420 |
| ADNI | ADNI5 | Siemens Prisma | 10 | 0.607 | 592.4 |
| ADNI | ADNI50 | Philips Achieva | 4 | 3.000* | 420.0* |
| ADNI | ADNI50 | Philips Achieva dStream | 11 | 3.000* | 591.0* |
| ADNI | ADNI51 | Philips Ingenia | 3 | 3.000* | 591.0* |
| ADNI | ADNI52 | GE DISCOVERY MR750w | 9 | 3 | 600 |
| ADNI | ADNI54 | Philips Achieva | 3 | 3 | 420 |
| ADNI | ADNI55 | Siemens Verio | 9 | 3 | 591 |
| ADNI | ADNI58 | Siemens Prisma fit | 25 | 3 | 591 |
| ADNI | ADNI59 | Philips Achieva | 3 | 3 | 420 |
| ADNI | ADNI59 | Siemens Prisma fit | 28 | 0.607* | 592.4* |
| ADNI | ADNI6 | GE DISCOVERY MR750w | 5 | 3 | 600 |
| ADNI | ADNI60 | Philips Ingenia | 5 | 3 | 591 |
| ADNI | ADNI61 | Philips Ingenia | 3 | 3 | 591 |
| ADNI | ADNI62 | Philips Achieva dStream | 4 | 3 | 591 |
| ADNI | ADNI63 | Siemens Prisma | 3 | 0.607 | 592.4 |
| ADNI | ADNI7 | Siemens MAGNETOM Prisma | 1 | 0.607 | 292.6 |
| ADNI | ADNI8 | Siemens Prisma fit | 8 | 3 | 591 |
| ADNI | ADNI9 | Philips Achieva | 7 | 3.000* | 420.0* |
| GUSTO | GUSTO_NUS | Siemens Prisma 1 | 336 | 2.62 | 303.9 |
| GUSTO | GUSTO_NUS | Siemens Prisma 2 | 191 | 2.62 | 469 |
| HCP-A | HCP-LS_Harvard | Siemens Prisma | 161 | 0.8 | 382.4 |
| HCP-A | HCP-LS_UCLA | Siemens Prisma | 147 | 0.8 | 382.4 |
| HCP-A | HCP-LS_UMinn | Siemens Prisma | 197 | 0.8 | 382.4 |
| HCP-A | HCP-LS_WashU | Siemens Prisma | 207 | 0.8 | 382.4 |
| HCP-D | HCP-LS_Harvard | Siemens Prisma | 209 | 0.8 | 382.4 |
| HCP-D | HCP-LS_UCLA | Siemens Prisma | 114 | 0.8 | 382.4 |
| HCP-D | HCP-LS_UMinn | Siemens Prisma | 159 | 0.8 | 382.4 |
| HCP-D | HCP-LS_WashU | Siemens Prisma | 130 | 0.8 | 382.4 |
| HCP-YA | HCP-YA1 | Siemens Prisma | 1029 | 0.72 | 864 |
| LIFE | NUS | Siemens Prisma fit | 240 | 1 | 596 |
| MACC | MACC_NUS | Siemens Trio | 115 | 2.4 | 297.6 |
| PNC | PNC1 | Siemens Trio | 885 | 3 | 372 |
| SG70 | NUS | Siemens Prisma fit | 942 | 1 | 596 |
| SINGER | NTU | Siemens Prisma fit | 225 | 1 | 596 |
| SINGER | NUS | Siemens Prisma fit | 634 | 1 | 596 |
| TCP | TCP1 | Siemens Prisma | 58 | 0.8 | 387.2 |
| TCP | TCP2 | Siemens Prisma | 33 | 0.8 | 387.2 |
| devCCNP | CCNP_CKG | Siemens Trio | 156 | 2.5 | 450 |
| devCCNP | CCNP_PEK | GE MR750 | 48 | 2 | 352 |
| eNKI | NKI1 | Siemens Trio | 689 | 1.4 | 560 |

Note: * indicates multiple TR/scan duration combinations exist for this site-scanner combination. The most common values are shown.

### Supplemental Figures

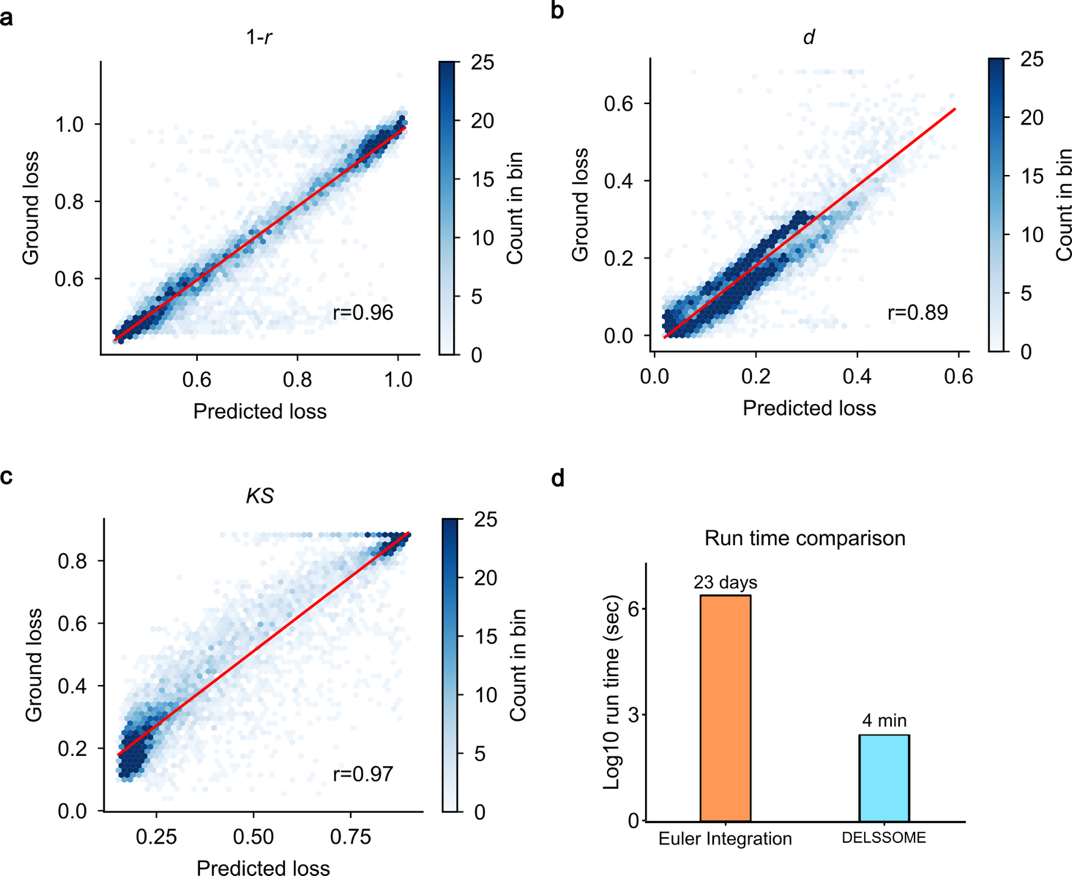

**Figure S1. Test performance of DELSSOME FC+FCD cost predictor based on the 100-region Yan parcellation.** This figure is the same as Fig. 2, but uses the 100-region Yan parcellation (Yan et al., 2023) instead of the Desikan-Killiany parcellation (Desikan et al., 2006). **a.** Test performance of DELSSOME prediction of static FC cost (*1 - r*). **b.** Test performance of DELSSOME prediction of static FC cost (*d*). **c.** Test performance of DELSSOME prediction of FCD cost (*KS*). The correlation between the predicted and ground truth loss were at least 0.89. In all the analyses, ground truth was defined based on Euler integration, while the DELSSOME models avoided the Euler integration. **d.** Run time (log scale) of DELSSOME versus Euler integration in evaluating FIC model realism. DELSSOME offers a 2000× speed up over Euler integration.

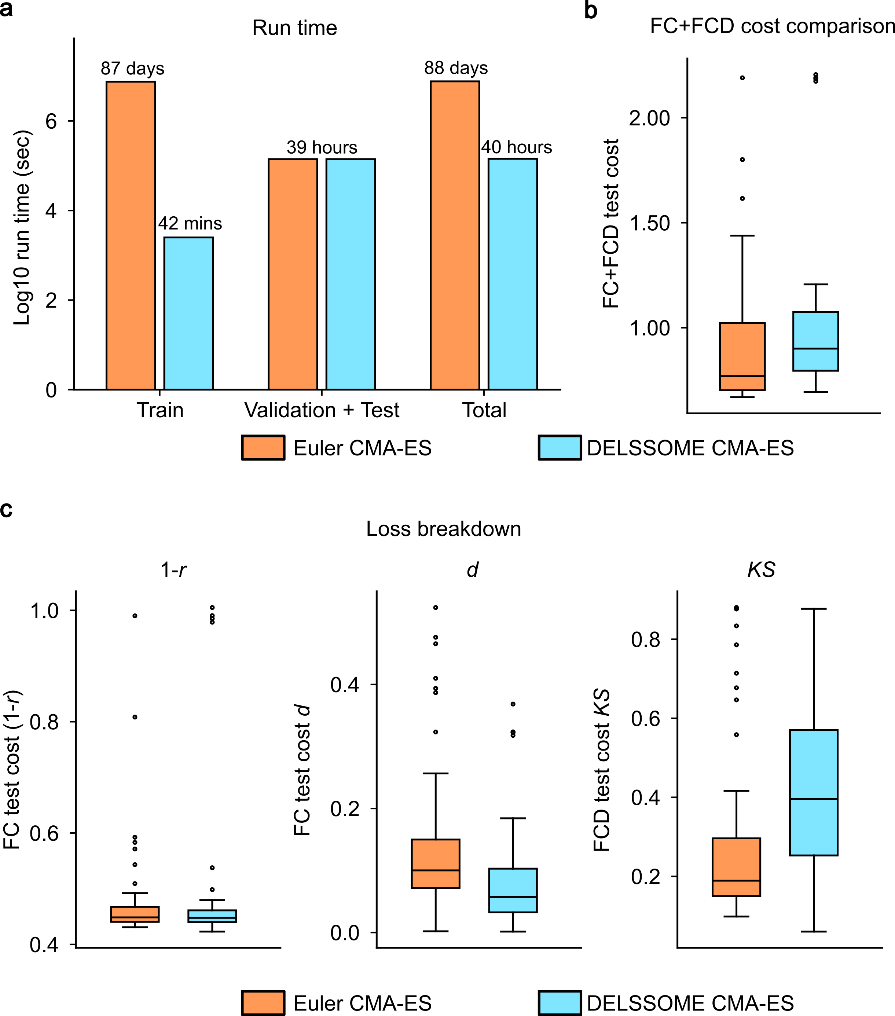

**Figure S2. Comparison of Euler CMA-ES and DELSSOME CMA-ES in the HCP-YA dataset using the 100-region Yan parcellation.** This figure is similar to Fig. 3 but uses the Yan parcellation (Yan et al., 2023). **a.** Run time (log scale) of DELSSOME CMA-ES versus Euler CMA-ES. DELSSOME CMA-ES offers a 3000× speed up over Euler CMA-ES in the training phase. If we also included validation and test phases in the run time, DELSSOME CMA-ES offers a 50× speed up over Euler CMA-ES. **b.** Total FC+FCD test cost comparison between DELSSOME CMA-ES and Euler CMA-ES. The cost is comparable (two-sample t-test, p=0.19). Each boxplot contains 50 data points corresponding to the 50 repetitions of the procedure in panel (a). **c.** Breakdown of the FC+FCD test cost from panel C into the two FC costs (*1-r* and *d*) and one FCD cost (*KS*). DELSSOME significantly sped up the estimation of the FIC model parameters without any degradation in estimation quality.

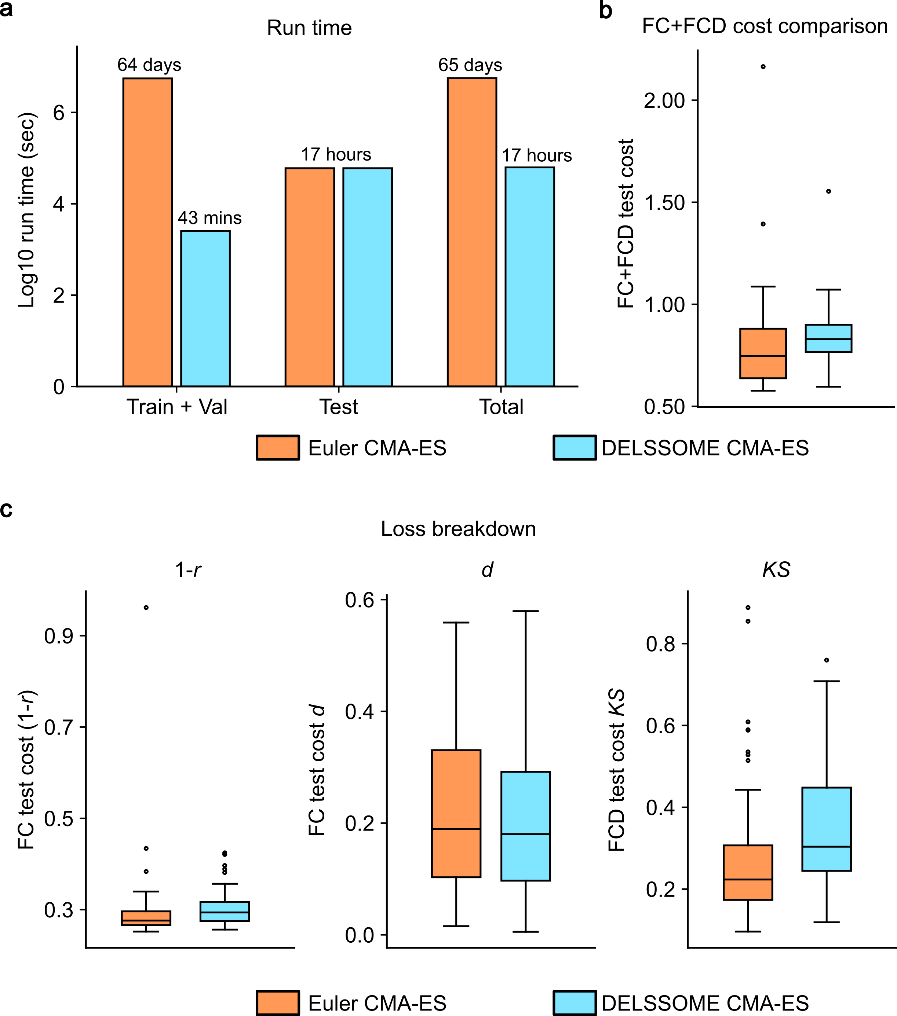

**Figure S3. Comparison of Euler CMA-ES and DELSSOME CMA-ES in the HCP-YA test set while using DELSSOME cost predictor in the validation phase**. **a**. Run time (log scale) of DELSSOME CMA-ES versus Euler CMA-ES. DELSSOME CMA-ES offers a 2000× speed up over Euler CMA-ES in the training and validation phases. If we also included the test phase in the run time, then DELSSOME CMA-ES offers a 100× speed up over Euler CMA-ES. **b**. Total FC+FCD test cost comparison between DELSSOME CMA-ES and Euler CMA-ES when using DELSSOME cost predictor in the validation phase. The cost was comparable (two-sample t-test p = 0.24). Each boxplot contains 50 data points corresponding to the 50 repetitions of the benchmarking procedure. **c**. Breakdown of the FC+FCD test cost from panel b into the two FC costs (*1-r* and *d*) and one FCD cost (*KS*).

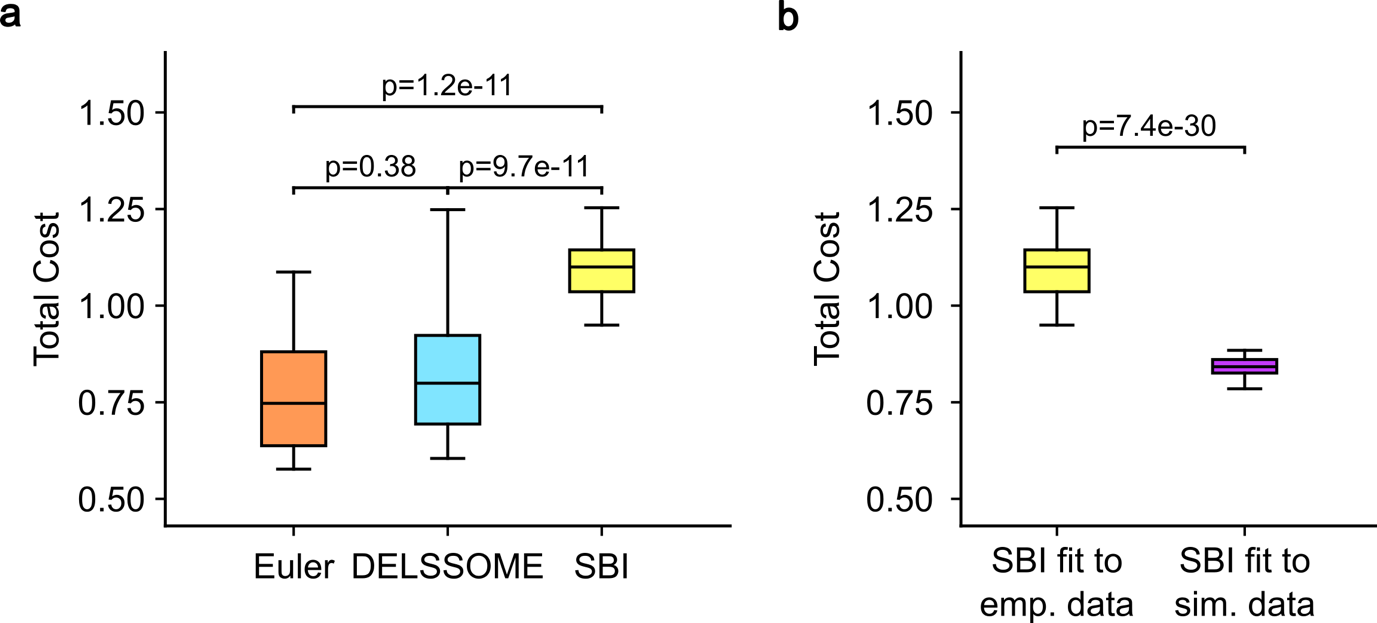

**Figure S4. Benchmarking DELSSOME against simulation-based inference (SBI) for FIC model parameter estimation.** **a.** Given empirical FC and FCD, DELSSOME CMA-ES produced FIC model parameter estimates that were comparable in quality to Euler CMA-ES. However, SBI inferred model parameters that were significantly worse than DELSSOME CMA-ES and Euler CMA-ES. Each boxplot contains 50 data points corresponding to 50 repetitions of the procedure. P-values were computed using two-sided paired-sample t-tests. **b.** SBI produces good FIC model parameter estimates when provided with simulated FC and FCD (purple), but performed substantially worse when applied to empirical FC and FCD (yellow). Note that the yellow boxplots in (a) and (b) are identical. Each boxplot contains 50 data points corresponding to 50 repetitions of the procedure. P values were obtained with a two-sided two-sample t-tests. Together, these results suggest that SBI performs well when simulated and empirical data are closely matched, but degrades when there is a mismatch between simulations and empirical data. See Supplementary Methods S9 for additional details.

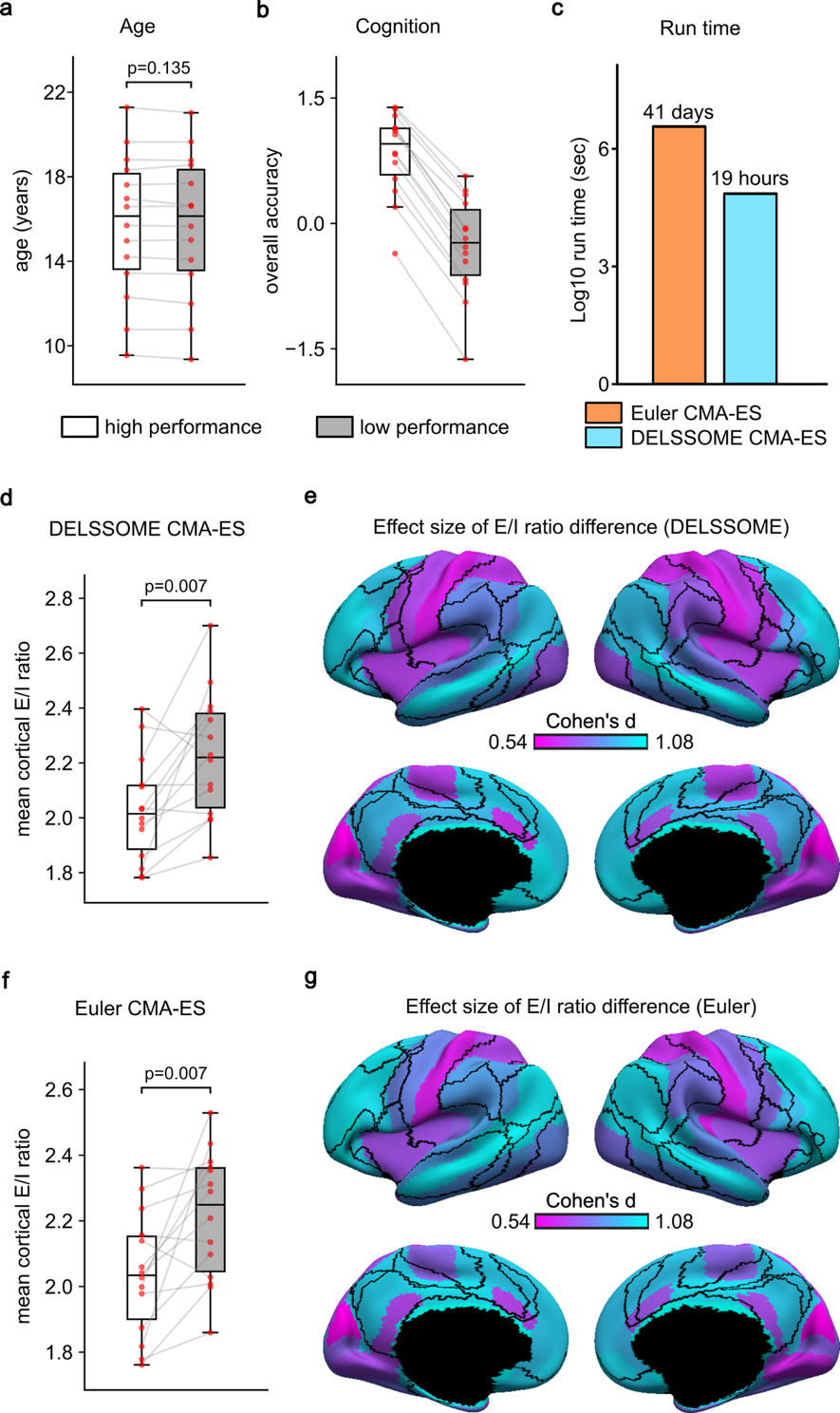

**Figure S5. DELSSOME CMA-ES reveals lower E/I ratio in youth with high cognitive performance, consistent with Euler CMA-ES. a.** Age of low and high cognitive performance groups. **b.** Cognitive performance (“overall accuracy”) of high and low performance groups. **c.** Run time comparison between DELSSOME CMA-ES and Euler CMA-ES. DELSSOME CMA-ES offers a 50× speed-up over Euler CMA-ES. **d.** Comparison of mean cortical E/I ratio between high-performance and low-performance groups estimated by DELSSOME CMA-ES. **e.** Regional differences in cortical E/I ratio between high-performance and low-performance groups estimated by DELSSOME CMA-ES. **f.** Comparison of mean cortical E/I ratio between high-performance and low-performance groups estimated by Euler CMA-ES. **g.** Regional differences in cortical E/I ratio between high-performance and low-performance groups estimated by Euler CMA-ES. 66 out of 68 regional differences were significant after FDR correction with q < 0.05 for both approaches. Pearson’s correlation between the 28 pairs of E/I ratio (from panels d and f) was 0.85. The analysis in this figure followed our previous study (Zhang et al., 2024). For more methodological details, see Supplementary Methods S4.

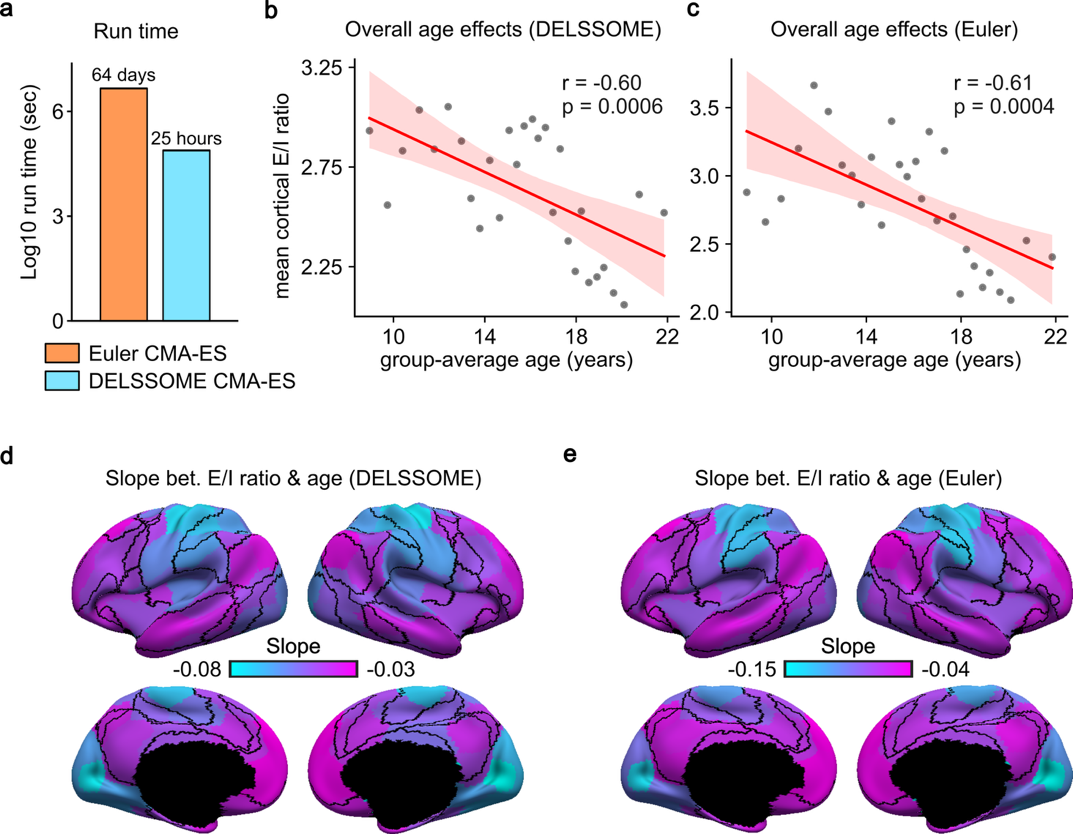

**Figure S6. DELSSOME CMA-ES generalizes to the Philadephia Neurodevelopmental Cohort (PNC) based on the 100-region Yan parcellation without further tuning**. **a**. Run time comparison between DELSSOME CMA-ES and Euler CMA-ES. DELSSOME CMA-ES offers a 50× speed-up over Euler CMA-ES. **b**. Correlation between age and mean cortical E/I ratio estimated by DELSSOME CMA-ES. **c**. Correlation between age and mean cortical E/I ratio estimated by Euler CMA-ES. **d**. Regression slope between age and regional E/I ratio estimated by DELSSOME CMA-ES. **e**. Regression slope between age and regional E/I ratio estimated by Euler CMA-ES. All slopes in panels d and e are negative. 100 out of 100 regions in panel d and 98 out of 100 regions in panel e were significant after multiple comparisons correction with false discovery rate (FDR) q < 0.05.

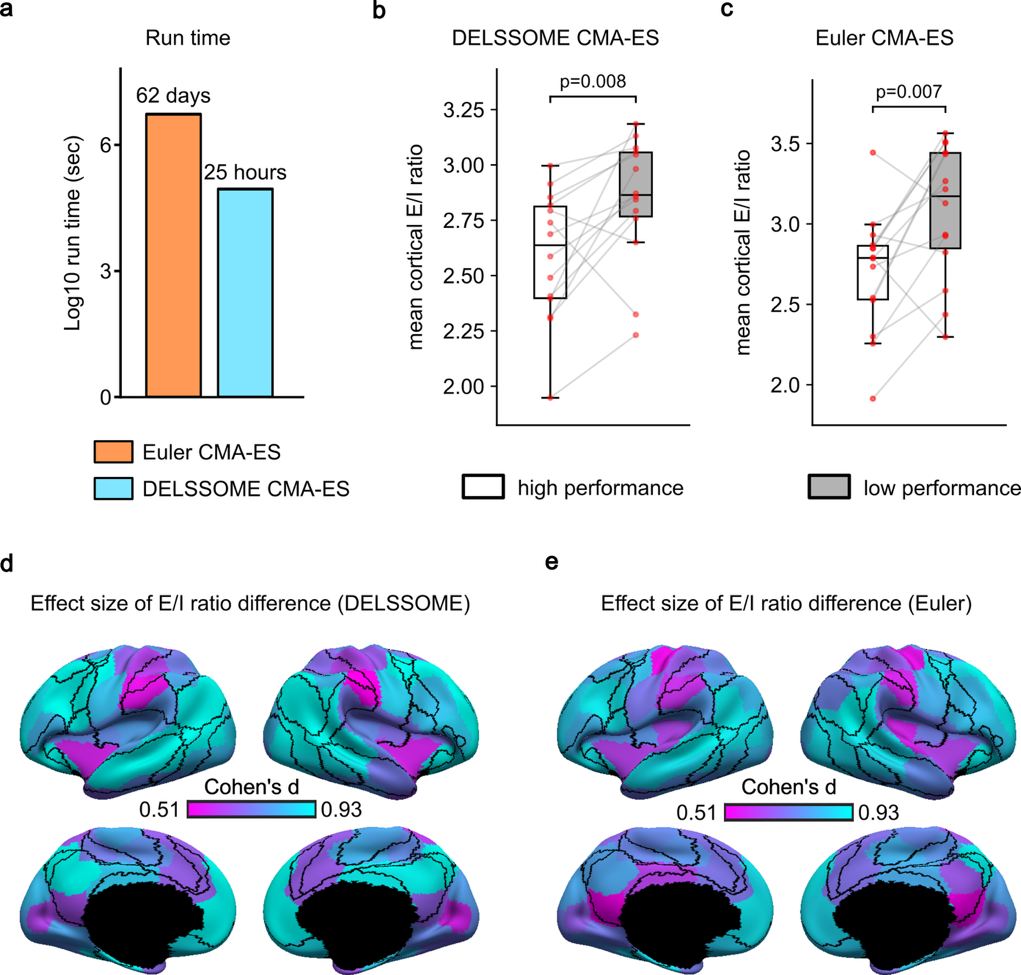

**Figure S7. DELSSOME CMA-ES reveals lower E/I ratio in youth with high cognitive performance, consistent with Euler CMA-ES, based on the 100-region Yan parcellation**. **a**. Run time comparison between DELSSOME CMA-ES and Euler CMA-ES. DELSSOME CMA-ES offers a 50× speed-up over Euler CMA-ES. **b**. Comparison of mean cortical E/I ratio between high-performance and low-performance groups estimated by DELSSOME CMA-ES. **c**. Comparison of mean cortical E/I ratio between high-performance and low-performance groups estimated by Euler CMA-ES. **d**. Regional differences in cortical E/I ratio between high-performance and low-performance groups estimated by DELSSOME CMA-ES. **e**. Regional differences in cortical E/I ratio between high-performance and low-performance groups estimated by Euler CMA-ES. 93 out of 100 regions in panel d, while 95 out of 100 regions in panel e were significant after FDR correction with q < 0.05.

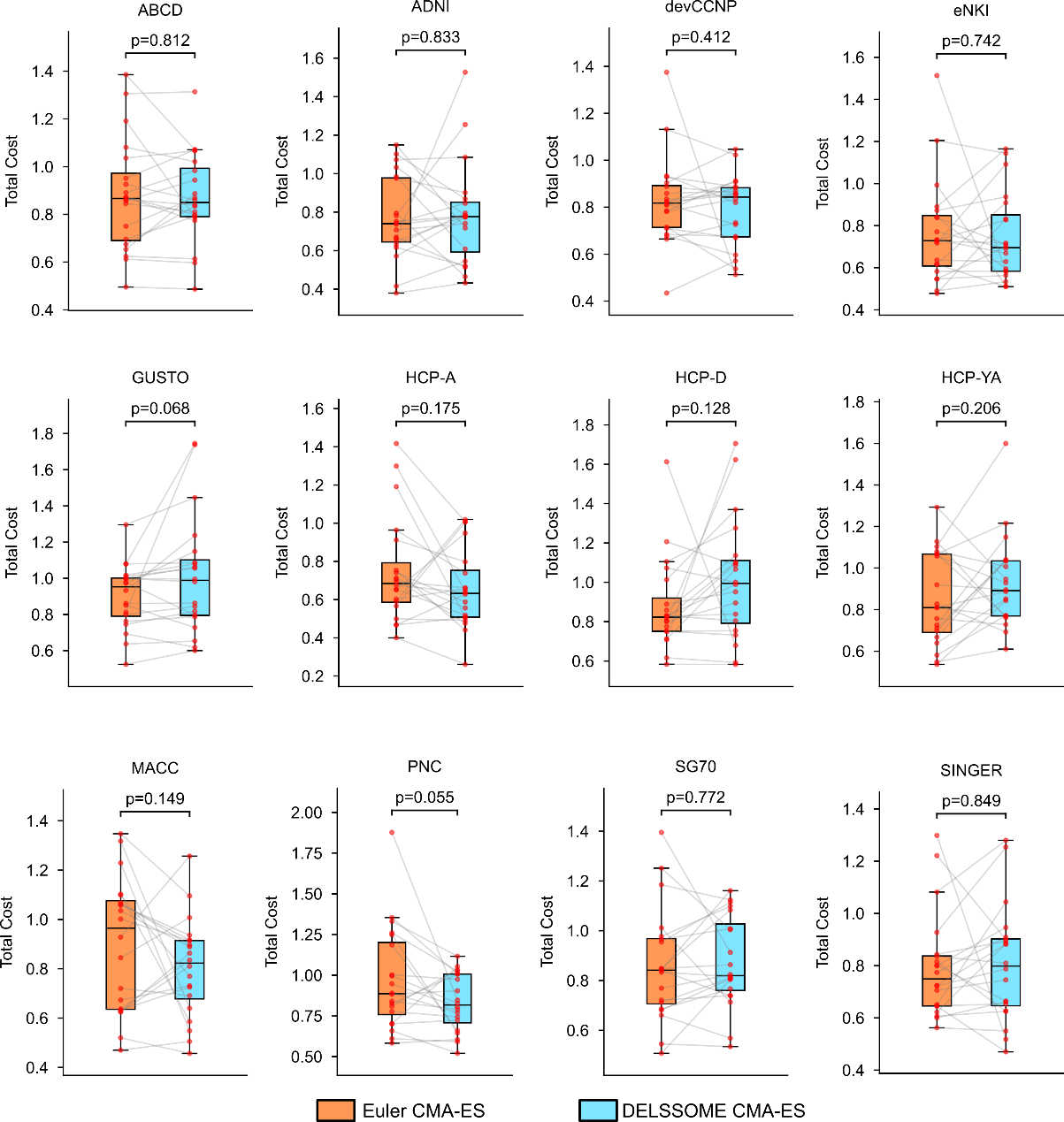

**Figure S8. Comparison of Euler CMA-ES and DELSSOME CMA-ES in individual-level FIC parameter optimization for 12 datasets**. **a**. For every dataset, DELSSOME CMA-ES achieves comparable performance to Euler CMA-ES with p > 0.05 even without multiple comparisons correction. Each red dot represents the cost of a test participant. The lines connect paired results from Euler CMA-ES and DELSSOME CMA-ES.

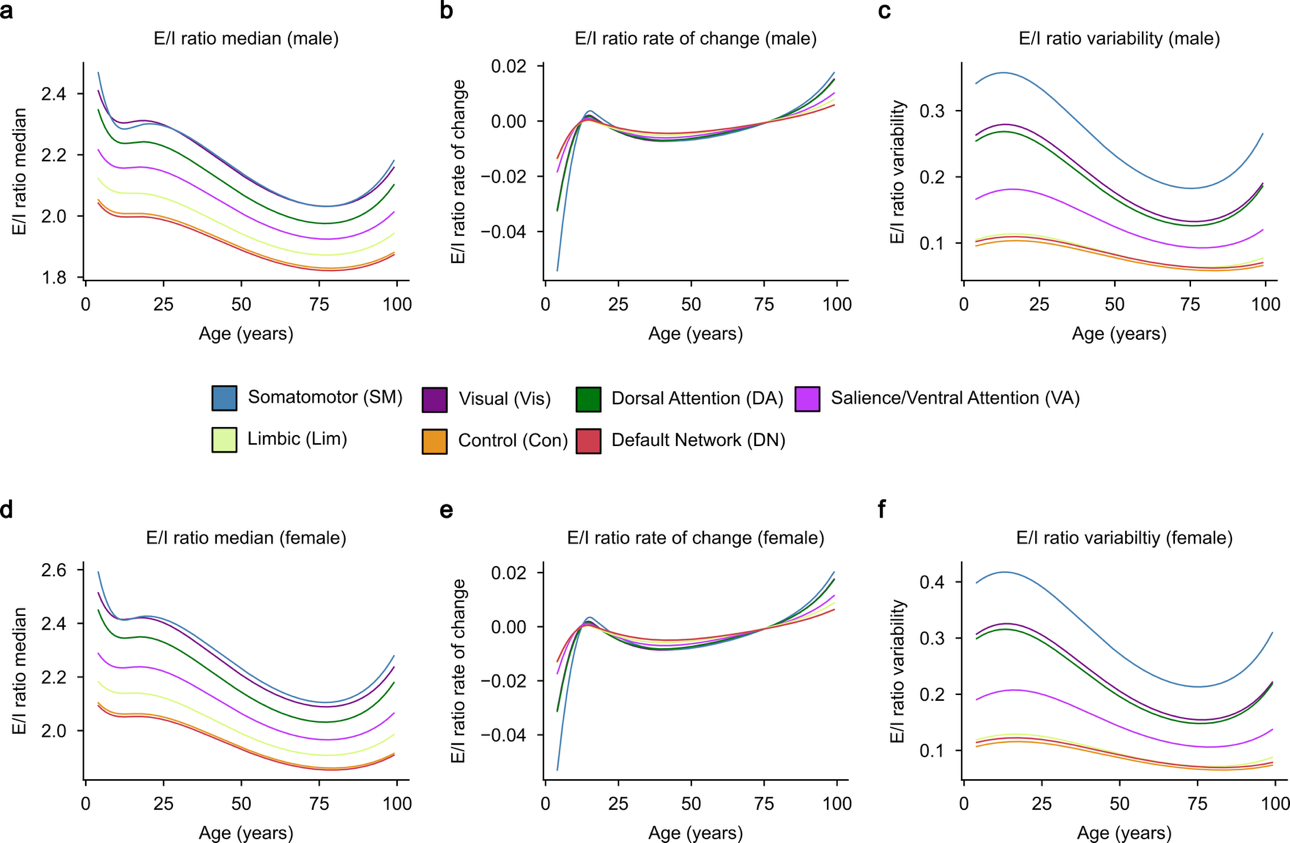

**Figure S9. Normative trajectories of cortical excitation-inhibition (E/I) ratio for each network in males and females. a.** Normative trajectories of E/I ratio (median) for each network in males. **b.** Rate of change of the median E/I trajectories (first derivative of the median curve with respect to age) for each network in males. **c.** Trajectories of between-individual variability in E/I ratio (scale parameter σ from the fitted GAMLSS) in males. **d.** Same as panel a, but for females. **e.** Same as panel b, but for females. **f.** Same as panel c, but for females.

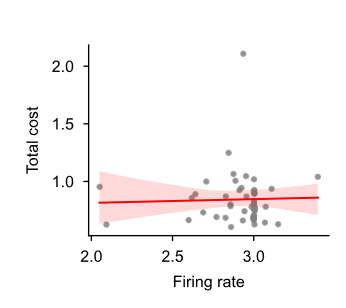

**Figure S10. Quality of FIC model parameters is independent of firing rate.** Scatter plot of FC+FCD test cost versus firing rate across the 50 repetitions of the benchmarking procedure in the HCP test set. Each dot represents the test cost and firing rate that correspond to the parameter set with lowest validation cost for each repetition. No significant correlation was observed (Pearson *r* = 0.03, *p* = 0.83). In an early version of DELSSOME, a within-range classifier was used to identify parameter sets producing firing rates within a predefined physiological range (2.7–3.3 Hz) before cost evaluation. Given this current result, the within-range classifier was removed from the final DELSSOME framework.

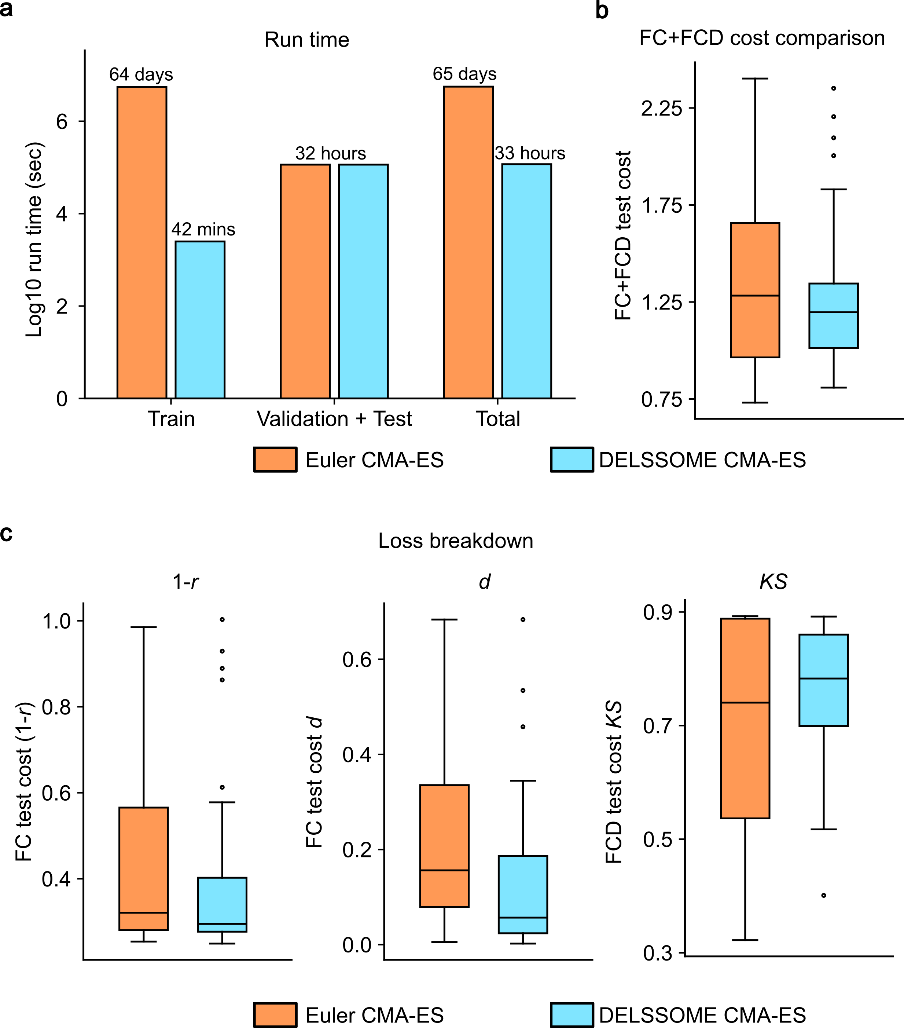

**Figure S11. DELSSOME performance with alternative firing-rate fixed point of 0.41Hz.** Comparison of Euler CMA-ES and DELSSOME CMA-ES in the HCP test set with the firing-rate fixed point set to 0.41 Hz (based on L2/3 excitatory neuron spontaneous rates; De Kock & Sakmann, 2008) instead of the default ~3 Hz. **a**. Run time (log scale) of DELSSOME CMA-ES versus Euler CMA-ES. The same acceleration pattern is observed at the alternative fixed point. **b**. Total FC+FCD test cost comparison between DELSSOME CMA-ES and Euler CMA-ES (two-sample t-test, p = 0.14). Each boxplot contains 50 data points corresponding to the 50 repetitions of the benchmarking procedure. **c**. Breakdown of the FC+FCD test cost from panel b into the two FC costs (1−*r* and *d*) and one FCD cost (*KS*).

### References

Akiba, T., Sano, S., Yanase, T., Ohta, T., & Koyama, M. (2019). Optuna: A Next-generation Hyperparameter Optimization Framework. *Proceedings of the 25th ACM SIGKDD International Conference on Knowledge Discovery & Data Mining*, 2623–2631. https://doi.org/10.1145/3292500.3330701

Bethlehem, R. A., Seidlitz, J., White, S. R., Vogel, J. W., Anderson, K. M., Adamson, C., Adler, S., Alexopoulos, G. S., Anagnostou, E., & Areces-Gonzalez, A. (2022). Brain charts for the human lifespan. *Nature*, *604*(7906), 525–533.

Boelts, J., Deistler, M., Gloeckler, M., Tejero-Cantero, Á., Lueckmann, J.-M., Moss, G., Steinbach, P., Moreau, T., Muratore, F., Linhart, J., Durkan, C., Vetter, J., Miller, B. K., Herold, M., Ziaeemehr, A., Pals, M., Gruner, T., Bischoff, S., Krouglova, N., … Macke, J. H. (2025). *sbi reloaded: A toolkit for simulation-based inference workflows* (arXiv:2411.17337). arXiv. https://doi.org/10.48550/arXiv.2411.17337

Brunel, N., & Wang, X.-J. (2001). Effects of Neuromodulation in a Cortical Network Model of Object Working Memory Dominated by Recurrent Inhibition. *Journal of Computational Neuroscience*, *11*(1), 63–85. https://doi.org/10.1023/A:1011204814320

De Kock, C. P. J., & Sakmann, B. (2008). High frequency action potential bursts (≥ 100 Hz) in L2/3 and L5B thick tufted neurons in anaesthetized and awake rat primary somatosensory cortex. *The Journal of Physiology*, *586*(14), 3353–3364. https://doi.org/10.1113/jphysiol.2008.155580

de Reus, M. A., & van den Heuvel, M. P. (2013). Estimating false positives and negatives in brain networks. *Neuroimage*, *70*, 402–409.

Deco, G., Ponce-Alvarez, A., Hagmann, P., Romani, G. L., Mantini, D., & Corbetta, M. (2014). How Local Excitation-Inhibition Ratio Impacts the Whole Brain Dynamics. *Journal of Neuroscience*, *34*(23), 7886–7898. https://doi.org/10.1523/JNEUROSCI.5068-13.2014

Deco, G., Ponce-Alvarez, A., Mantini, D., Romani, G. L., Hagmann, P., & Corbetta, M. (2013). Resting-state functional connectivity emerges from structurally and dynamically shaped slow linear fluctuations. *Journal of Neuroscience*, *33*(27), 11239–11252.

Demirtaş, M., Burt, J. B., Helmer, M., Ji, J. L., Adkinson, B. D., Glasser, M. F., Van Essen, D. C., Sotiropoulos, S. N., Anticevic, A., & Murray, J. D. (2019). Hierarchical heterogeneity across human cortex shapes large-scale neural dynamics. *Neuron*, *101*(6), 1181–1194.

Desikan, R. S., Ségonne, F., Fischl, B., Quinn, B. T., Dickerson, B. C., Blacker, D., Buckner, R. L., Dale, A. M., Maguire, R. P., & Hyman, B. T. (2006). An automated labeling system for subdividing the human cerebral cortex on MRI scans into gyral based regions of interest. *Neuroimage*, *31*(3), 968–980.

Devlin, J., Chang, M.-W., Lee, K., & Toutanova, K. (2019). *BERT: Pre-training of Deep Bidirectional Transformers for Language Understanding* (arXiv:1810.04805). arXiv. https://doi.org/10.48550/arXiv.1810.04805

Glasser, M. F., & Van Essen, D. C. (2011). Mapping human cortical areas in vivo based on myelin content as revealed by T1-and T2-weighted MRI. *Journal of Neuroscience*, *31*(32), 11597–11616.

Gonçalves, P. J., Lueckmann, J.-M., Deistler, M., Nonnenmacher, M., Öcal, K., Bassetto, G., Chintaluri, C., Podlaski, W. F., Haddad, S. A., & Vogels, T. P. (2020). Training deep neural density estimators to identify mechanistic models of neural dynamics. *Elife*, *9*, e56261.

Greenberg, D., Nonnenmacher, M., & Macke, J. (2019). Automatic posterior transformation for likelihood-free inference. *International Conference on Machine Learning*, 2404–2414. https://proceedings.mlr.press/v97/greenberg19a.html

Hansen, E. C., Battaglia, D., Spiegler, A., Deco, G., & Jirsa, V. K. (2015). Functional connectivity dynamics: Modeling the switching behavior of the resting state. *Neuroimage*, *105*, 525–535.

Kingma, D. P. (2014). Adam: A method for stochastic optimization. *arXiv Preprint arXiv:1412.6980*. https://scholar.google.com/scholar?cluster=16194105527543080940&hl=en&inst=569367360547434339&inst=3212728378801010220&oi=scholarr

Kong, X., Kong, R., Orban, C., Wang, P., Zhang, S., Anderson, K., Holmes, A., Murray, J. D., Deco, G., van den Heuvel, M., & Yeo, B. T. T. (2021). Sensory-motor cortices shape functional connectivity dynamics in the human brain. *Nature Communications*, *12*(1), Article 1. https://doi.org/10.1038/s41467-021-26704-y

Margulies, D. S., Ghosh, S. S., Goulas, A., Falkiewicz, M., Huntenburg, J. M., Langs, G., Bezgin, G., Eickhoff, S. B., Castellanos, F. X., Petrides, M., Jefferies, E., & Smallwood, J. (2016). Situating the default-mode network along a principal gradient of macroscale cortical organization. *Proceedings of the National Academy of Sciences*, *113*(44), 12574–12579. https://doi.org/10.1073/pnas.1608282113

Papamakarios, G., Pavlakou, T., & Murray, I. (2017). Masked autoregressive flow for density estimation. *Advances in Neural Information Processing Systems*, *30*. https://proceedings.neurips.cc/paper_files/paper/2017/hash/6c1da886822c67822bcf3679d04369fa-Abstract.html

Ponce-Alvarez, A., & Deco, G. (2024). The Hopf whole-brain model and its linear approximation. *Scientific Reports*, *14*(1), 2615.

Stasinopoulos, D. M., & Rigby, R. A. (2008). Generalized additive models for location scale and shape (GAMLSS) in R. *Journal of Statistical Software*, *23*, 1–46.

Stephan, K. E., Weiskopf, N., Drysdale, P. M., Robinson, P. A., & Friston, K. J. (2007). Comparing hemodynamic models with DCM. *Neuroimage*, *38*(3), 387–401.

Sun, L., Zhao, T., Liang, X., Xia, M., Li, Q., Liao, X., Gong, G., Wang, Q., Pang, C., & Yu, Q. (2025). Human lifespan changes in the brain’s functional connectome. *Nature Neuroscience*, 1–11.

Tolley, N., Rodrigues, P. L., Gramfort, A., & Jones, S. R. (2024). Methods and considerations for estimating parameters in biophysically detailed neural models with simulation based inference. *PLOS Computational Biology*, *20*(2), e1011108.

Wang, P., Kong, R., Kong, X., Liégeois, R., Orban, C., Deco, G., van den Heuvel, M. P., & Thomas Yeo, B. T. (2019). Inversion of a large-scale circuit model reveals a cortical hierarchy in the dynamic resting human brain. *Science Advances*, *5*(1), eaat7854. https://doi.org/10.1126/sciadv.aat7854

Wong, K.-F., & Wang, X.-J. (2006). A recurrent network mechanism of time integration in perceptual decisions. *Journal of Neuroscience*, *26*(4), 1314–1328.

Yan, X., Kong, R., Xue, A., Yang, Q., Orban, C., An, L., Holmes, A. J., Qian, X., Chen, J., & Zuo, X.-N. (2023). Homotopic local-global parcellation of the human cerebral cortex from resting-state functional connectivity. *NeuroImage*, *273*, 120010.

Zhang, S., Larsen, B., Sydnor, V. J., Zeng, T., An, L., Yan, X., Kong, R., Kong, X., Gur, R. C., Gur, R. E., Moore, T. M., Wolf, D. H., Holmes, A. J., Xie, Y., Zhou, J. H., Fortier, M. V., Tan, A. P., Gluckman, P., Chong, Y. S., … Yeo, B. T. T. (2024). In vivo whole-cortex marker of excitation-inhibition ratio indexes cortical maturation and cognitive ability in youth. *Proceedings of the National Academy of Sciences*, *121*(23), e2318641121. https://doi.org/10.1073/pnas.2318641121
